# Bioorthogonal caging-group-free photoactivatable probes for minimal-linkage-error nanoscopy

**DOI:** 10.1101/2023.06.21.545866

**Authors:** Ayse Aktalay, Richard Lincoln, Lukas Heynck, Maria Augusta do R. B. F. Lima, Alexey N. Butkevich, Mariano L. Bossi, Stefan W. Hell

## Abstract

Here we describe highly compact, click compatible, and photoactivatable dyes for super-resolution fluorescence microscopy (nanoscopy). By combining the photoactivatable xanthone (PaX) core with a tetrazine group, we achieve minimally sized and highly sensitive molecular dyads for selective labeling of unnatural amino acids introduced by genetic code expansion. We exploit the excited state quenching properties of the tetrazine group to attenuate the photoactivation rates of the PaX, and further reduce the overall fluorescence emission of the photogenerated fluorophore, providing two mechanisms of selectivity to reduce off-target signal. Coupled with MINFLUX nanoscopy, we demonstrate our dyads in the minimal-linkage-error imaging of vimentin filaments, demonstrating the molecular scale precision in fluorophore positioning.

## INTRODUCTION

Advancements in live-cell super-resolution fluorescence microscopy (nanoscopy) methods beget greater demands on the fluorophores and labels used for imaging, to fully leverage the improvements in optical resolution.^1-2^ To distinguish adjacent fluorophores at molecular-scale proximities, super-resolution methods rely on the sequential ‘on’–’off’ transitioning of fluorophores to generate distinct molecular states. This is commonly achieved with cyanines^3^ in complex imaging buffers that are incompatible with live-cell imaging.^4^ Cyanines further suffer from intermolecular energy transfer between dark states, limiting the molecular distances that can be separated.^5^ These constraints may be addressed by employing alternative stochastic blinking fluorophores,^6-12^ photoswitchable dyes,^13-14^ or photoactivatable (caged) dyes.^15-17^

Furthermore, as the improvements in obtainable optical resolution reach single nanometer precision, as it is the case for MINFLUX (minimal photon fluxes)^18-21^ and MINSTED nanoscopy techniques,^22-23^ the displacement of the fluorophore from the point of interest (referred to as linkage error) becomes a critical parameter.^1, 24-25^ Genetic code expansion has emerged as a powerful platform for site-specific labeling of proteins with small organic fluorophores with minimal linkage error.^2, 26^ Through incorporation of unnatural amino acids (UAAs) with bioorthogonal reactivity to fluorescent probes, proteins of interest can be labeled in a fast and specific manner.^27^ The most prominent example is the combination of “clickable” UAAs and tetrazine-functionalized fluorophores for highly-specific labeling by live-cell compatible strain-promoted inverse electron-demand Diels-Alder cycloaddition (SPIEDAC) reactions (Figure 1).^28^ A further advantage of tetrazine-fluorophore dyads is fluorogenicity, which arises if the fluorescence signal is partially or fully quenched by the unreacted tetrazine (Tz) fragment^29^ and then restored upon reaction with the strained alkyne.^30^ This “turn-on” effect has been extensively utilized to develop labels for no-wash imaging.^10, 31-33^ Despite their significant potential, only a handful of Tz-fluorophore dyads are compatible with fluorescence nanoscopy methods with sufficient optical resolution to observe the advantages in reduced linkage error,^2, 9, 26^ and fewer still are commercially available.^5^ To date, no Tz-fluorophore dyads have been utilized to label UAAs for MINFLUX imaging—the closest example is an azide plus variant of the cyanine dye Alexa647, in combination with copper-mediated click chemistry, and a dedicated buffer to achieve blinking.^26^

**Figure 1. a–c.**
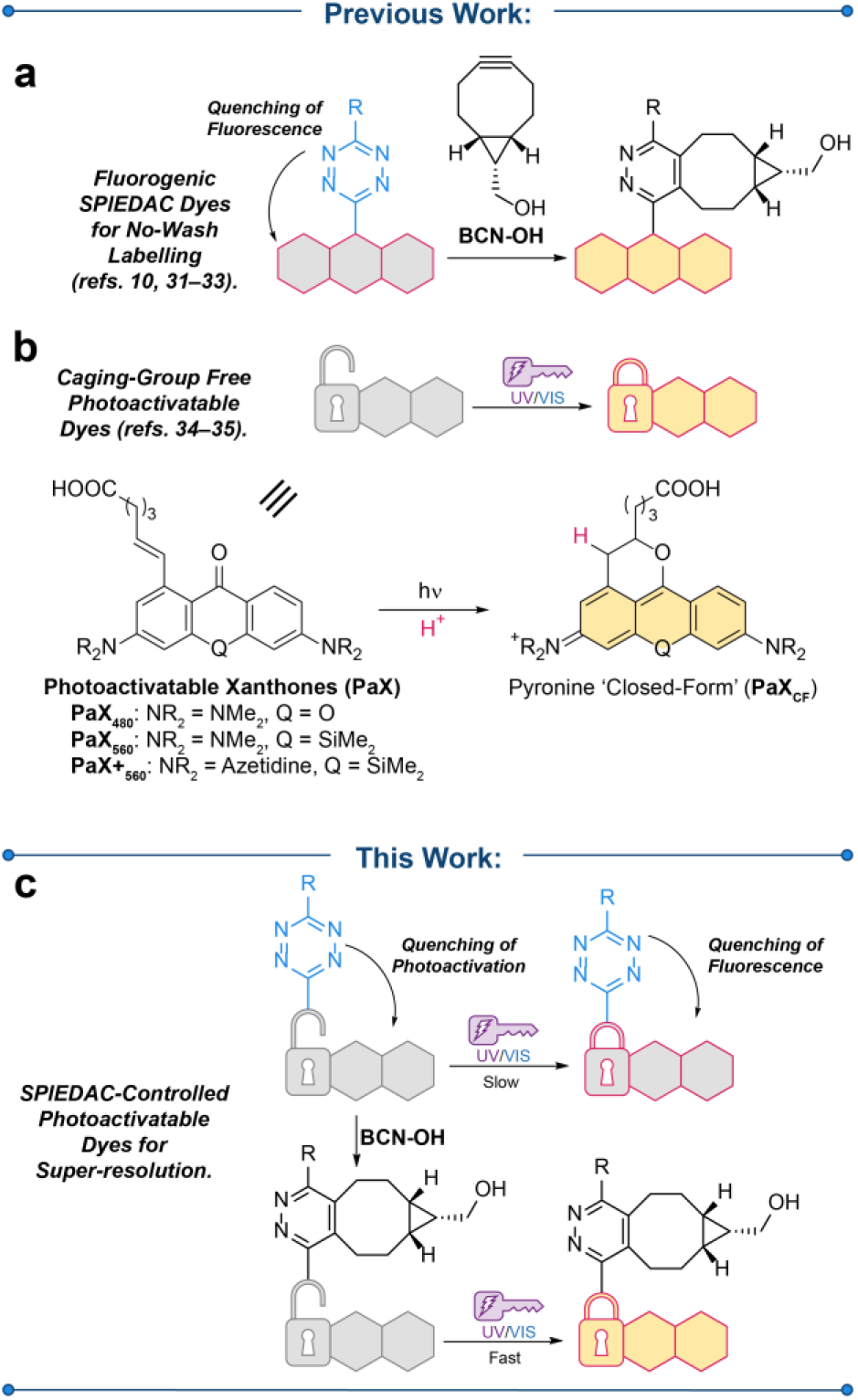
Rational design of a photoactivatable tetrazine dyad for super-resolution microscopy. Previous efforts to build fluorogenic labeling strategies utilizing (a) strain-promoted inverse electron demand Diels–Alder (SPIEDAC) reactions and (b) caging-group free photoactivatable fluorophores, including photoactivatable (PaX) dyes which undergo photoactivation to a highly fluorescent ‘closed form’ (PaXCF). This work (c) utilizes SPIEDAC to control both the brightness and the switching rates of photoactivatable fluorophores for nanoscopy.

We recently reported a new class of caging-group-free photoactivatable xanthone (PaX) analogues that upon UV (one photon) or NIR (two photon) irradiation convert rapidly and cleanly into highly fluorescent pyronine dyes in a triplet mediated reaction.^34-35^ Due to their small, uncharged structures, these PaX labels can be used for fixed or live-cell STED (stimulated emission depletion) and PALM (photoactivated light microscopy), as well as MINFLUX nanoscopy. We reasoned that tetrazine moieties, in addition to imparting fluorogenicity to UAA-specific labels, will further attenuate the rate of photoactivation of the PaX core by quenching of the xanthone excited state—precluding intersystem crossing and subsequent fluorophore formation. Thus, tetrazine functionalization of PaX dyes constitutes a suitable platform to construct compact, high-contrast photoactivatable labels with minimal linkage errors, ideally suited for MINFLUX nanoscopy.

We report herein a series of PaX dyads incorporating tetrazine moieties (PaX-Tz) for nanoscopy imaging, in which we systematically varied the xanthone analogue, tetrazine, and linking strategy between the two, in order to modulate photoactivation rates and the fluorescence contrast between the unreacted tetrazine (Tz) and reacted pyridazine (Pz) adducts of the pyronine fluorophore (Figure 1). The quenching imparted by the tetrazine is eliminated upon reaction with the target UAA, resulting in faster photoactivation and brighter fluorescence following labeling. Finally, we demonstrated the utility of these labels for STED, PALM, and MINFLUX nanoscopy of vimentin filaments incorporating UAA for truly molecular-scale visualization of protein-protein distances.

## RESULTS AND DISCUSSION

### Synthesis and characterization of PaX tetrazine labels

A number of diversely functionalized tetrazines have been reported, with improved rates of reactivity to UAAs, typically anticorrelated with their chemical stabilities.^28, 36-37^ We reasoned that an ideal combination of tetrazine (Tz) and photoactivatable xanthone (PaX) would result in quenching of the singlet excited state of the xanthone, via an energy transfer mechanism analogous to quenching observed in fluorescent tetrazine dyads,^38^ precluding intersystem crossing and, in turn, photo-assembly of the fluorophore.

In order to identify such a combination of PaX and Tz moieties, we synthesized a series of PaX-Tz dyads (Figure 2a). Constructed from the reported PaX carboxylic acids (PaX480, PaX560, and PaX+560),34 we first varied the diarylketone between an electron rich oxygen-bridged amino-xanthone (PaX480; **1, 2**) and electron-deficient silicon-bridged xanthones (PaX560, PaX+560) bearing either dimethylamine (**3, 4**) or azetidine auxochromic groups (**5, 6**) in order to elucidate electronic effects on quenching. We next varied the Tz moiety, selecting either the more-reactive unsubstituted (**1, 3, 5**) or more shelf-stable methyl-substituted (**2, 4, 6**) phenyl tetrazines. We further explored alternative linkage strategies between the Tz unit and the PaX (Figure 2b), in order to reduce the linkage error and the influence of distance on quenching.^38^ The first strategy involved ether-linked dyads (**7, 8**), derived from the corresponding 3-bromo-1,2,4,5-tetrazine^39^ and PaX560 analogue with an alcohol-terminated linker. However, we found that such constructs suffered from poor-reactivity of the Tz moiety, and were susceptible to hydrolytic cleavage of the Tz unit. In our second strategy, we constructed a PaX (**S9**, see Supplementary Information) with an acrylate moiety as the linker/radical trap, which served to generate a series of the dyads bearing either secondary (**9, 10**) or tertiary (**11, 12**) acrylamide linkages. Initial TD-DFT studies^29^ anticipated excited state energy transfer to the Tz moiety in both the singlet and triplet manifolds for the PaX structures, regardless of linkage (Figure S1a), as well as quenching of the singlet excited state of the photogenerated (i.e., ‘closed-form’) pyronine fluorophore (PaXCF, Figure 1b and Figure S1b). Recording the fluorescence lifetime of compound **4** at 490 nm before photoactivation showed significant shortening of the lifetime compared to PaX560 (Figure S2a) further affirming the ability of the Tz moiety to quench the singlet state of the diaryl ketone, and in turn, prevent photoswitching from the triplet state. After photoactivation of **4**, the fluorescence lifetime of the corresponding pyronine (**4a**) at 585 nm was also significantly shorter than that of the fluorophore photogenerated from PaX560, emphasizing a potential fluorogenic response (Figure S2b).

**Figure 2.**
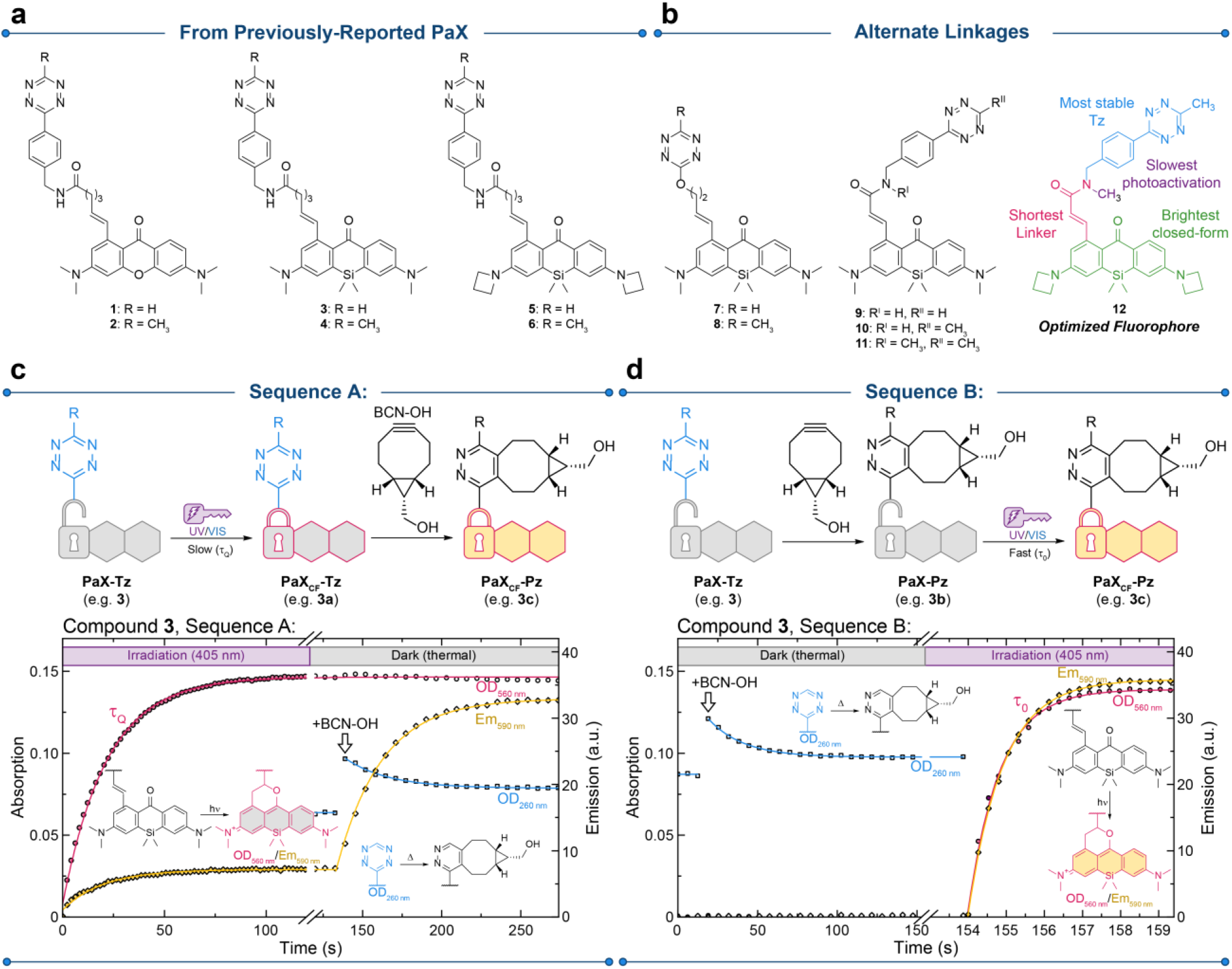
Characterization of PaX-Tz dyads. **a**, Chemical structures of the PaX-Tz dyads studied in this work derived from previously-reported PaX^34^. **b**, Alternate linkage designs. **c**, Effect of the Tz moiety on the photoactivation behavior of PaX-Tz dyes, exemplified with compound **3**. In Sequence A, the temporal evolution of the absorption and fluorescence spectra of **3** (**3**→**3a**; 1.66 μg mL^−1^) irradiated in methanol (λact = 405 nm) was monitored. BCN-OH (50 eq.) was subsequently added and the effect of Tz-quenching of the fluorescence was observed (**3a**→**3c**). **d**, In Sequence B, BCN-OH was first allowed to react with **3** (**3**→**3b**), then following the temporal evolution of the absorption and fluorescence spectra upon irradiation (**3b**→**3c**).

To evaluate the effect of the Tz moiety on the photoactivation of the PaX chromophore, irradiation experiments were first conducted in methanol, prior and after complete reaction of the Tz moiety with (1*R*,8*S*,9*s*)-bicyclo[6.1.0]non-4-yn-9-ylmethanol (BCN-OH) to give the corresponding pyridazine (Py) product. In the first experimental sequence (Sequence A, Figure 2c) the dyads were irradiated with 405 nm light until complete conversion to the corresponding ‘closed-form’ (PaXCF; e.g., **3a**, see Figure S3a), which was monitored spectroscopically at the absorption and emission maxima of the PaXCF (560 nm and 590 nm, respectively, for compound **3**). This established the rate of photoactivation in the presence of Tz quenching (i.e., the rate of conversion from PaX-Tz to PaXCF-Tz). Excess BCN-OH (50 equiv.) was next added, resulting in an enhancement of the fluorescence signal (590 nm for **3a** to **3c**) concomitant with a decrease in absorption from the Tz (260 nm) as the SPIEDAC reaction occurred. In the second experimental sequence (Sequence B, Figure 2d) the dyads were first allowed to react completely with BCN-OH yielding the pyridazine dyad (PaX-Pz; e.g., **3b**, see Figure S3a), and subsequently photoactivated to the ‘closed-form’ (PaXCF-Pz), establishing the rate of photoactivation in the absence of the Tz moiety. Figure S3b shows the example spectral changes corresponding the Sequences A and B for compound **3** and Figure S4 shows the LC-MS analysis for intermediates **3a**–**3c**.

The designed experiment allows for a quantitative comparison of relevant properties for the presented dyads. The ratio of the photoactivation speed of the PaX-Tz and PaX-Pz provides a measure of the Tz quenching of the photoactivation process for all the dyads (Figure S5a). Additionally, the ratio of apparent fluorescence quantum yields of the PaXCF-Tz and PaXCF-Pz provided a measure of the fluorogenicity of the compounds (Figure S5b) and allowed comparison of quantum yields across the family of PaXCF (Figure S5c). Lastly, LC-MS analysis was performed on the reaction mixtures and revealed no major byproducts or photoproducts of Tz quenching, unless otherwise indicated below (Figure S6).

In general, we observed an increase in photoactivation speed after reaction with BCN-OH, confirming quenching of the photoactivation pathway by the Tz moiety, and an approx. 5-fold fluorogenic response of the PaXCF-Tz upon reaction with BCN-OH. More specifically, for the compounds derived from PaX480 (**1, 2**, see Figure S7) the combination of slower photoactivation rates and lower photostability of the dye scaffold, when combined with the quenching of Tz, resulted in significant oxidative photoblueing^40-41^ from the prolonged irradiation over the course of Sequence A (Figure S7a-d), which was further confirmed in the LC-MS analysis (Figure S7e,f). This was most notable for compound **2** where the demethylated product was primarily observed. Compound **4** and the dyads derived from the azetidine functionalized PaX+560 compounds (**5, 6**) yielded comparable results to compound **3**, with compound **5** showing the greatest quenching of photoactivation and **6** showing the greatest fluorogenicity of all the dyads.

Changing the linkage strategy to an ether (**7, 8**) enabled shortening the Tz–Pax distance, however, the reduced stability of the ether resulted in slow loss of the Tz in methanol (Figure S8a), precluding their study by Sequence A. Both compounds **7** and **8** reacted quickly with BCN-OH and the resulting product was resistant to methanolysis (Figure S8b). Photoactivation of the dyad after reaction with BCN-OH was rapid and clean (Figure S8c,d). Overall, the electrophilicity of the alkoxytetrazines may preclude their use in live-cell experiments.

To prepare the acrylamide-linked PaX scaffolds (**9**–**12**), new acrylate-functionalized PaX were prepared (**S9** and **S12**, see Supporting information for details) and coupled to commercially-available tetrazine amines to yield the secondary acrylamide products (**9, 10**). To access the tertiary (*N*-methyl) acrylamides (**11, 12**), *N*-methyl-4-(6-methyl-1,2,4,5-tetrazin-3-yl)benzylamine^42^ was additionally synthesized and coupled with the acrylate PaX. Much to our surprise, the acrylamide-derived PaX scaffolds showed notably slower photoactivation rates, likely due to the greater electron deficiency of the radical trap. The secondary acrylamides further showed the establishment of an equilibrium with the nucleophilic addition of methanol solvent after photoactivation (Figure S9 for Compound **9**). For compound **10**, this was particularly apparent, with the primary photoproduct of Sequence A being non-absorbing at 570 nm and non-fluorescent while Sequence B (following reaction with BCN-OH) photoactivation was fast and yielded the desired **10c**, making this a promising candidate for no-wash labeling. Due to this, the effect of the Tz on the photoactivation rate could not be reliably determined for compounds **9** and **10**, but tertiary acrylamide-functionalized PaX scaffolds may prove to be a useful general strategy to reduce the photoactivation rate of PaX fluorophores.

With the reduced linker length of the acrylamides, we anticipated better quenching between the Tz and PaX. However, we observed only a modest fluorogenicity and photoactivation suppression for the tertiary acrylamides **11** and **12**. This may be due to the reduced flexibility of the acrylamide linker to adopt a π-stacked geometry that brings the Tz close to the pyronine fluorophore as compared to the dyads with longer linkers. Importantly, the acrylamide had no effect on the fluorescence quantum yield (ϕf) of the PaXCF-Pz, with the ϕf ultimately being determined by the nature of *N,N*-dialkylamino substituents (dimethyl vs azetidine, see Figure S5a). Compound **12** provided the ideal combination of properties for nanoscopy: an azetidine-bearing PaX scaffold for the highest ϕf of the closed form (**12c**); a secondary acrylamide linker for minimal size and slowest photoactivation, both before and after reaction with BCN-OH (for greater activation control during imaging); and a methyl-functionalized tetrazine for optimal shelf-life.

To determine if the fluorogenic effect and reactivity of our dyads translated into a more biologically relevant setting, and as a first demonstration of the close proximity of the label to its target, we reacted compound **3a** (prepared by photolysis of compound **3**) with a green fluorescent protein (GFP) variant bearing a Y35TAG point mutation for the incorporation of *endo*-BCN-*L*-lysine (Figure 3a).^43-47^ By monitoring the fluorescence of the GFP upon excitation at 470 nm, as well as the new emission band at 590 nm, we could observe Förster resonance energy transfer (FRET) from the GFP (donor) to the PaXCF (acceptor, see Figure 3b). Furthermore, by monitoring the emission of the PaXCF at 590 nm while directly exciting with 560 nm light, we could quantify the fluorescence enhancement of the fluorophore as the tetrazine moiety reacted with the BCN, observing a 4.4-fold enhancement in fluorescence, consistent with previous experiments. Mass spectrometry confirmed covalent labeling with **3a** (Figure S10).

**Figure 3.**
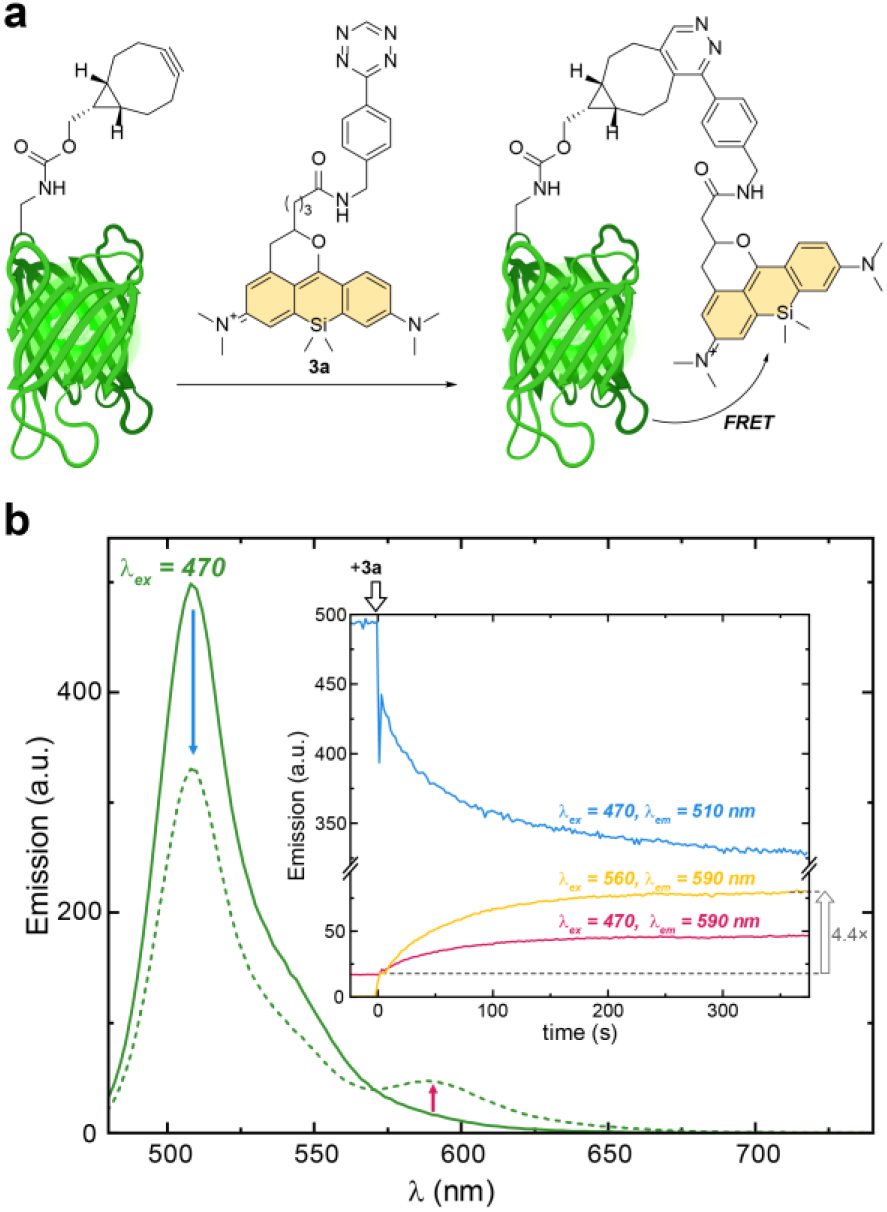
Demonstration of FRET in GFP labeled with a PaX-Tz fluorophores. **a**, Schematic representation of GFP-PaX dyad. GFP protein (1.1 eq.; for plasmid details see the Supporting Information) containing *endo*-BCN-*L*-lysine^43-47^ was reacted with **3a** in PBS at pH 7.4. **b**, Fluorescence spectra of GFP before (solid line) and after (dashed line) reaction with **3a**. Inset: Temporal evolution of fluorescence signal upon addition of **3a**, indicative of FRET from GFP to compound **3a** upon SPIEDAC reaction. Created with BioRender.com.

### Bioorthogonal probes for fluorescence nanoscopy

We next turned our attention to applying PaX-Tz dyads in fluorescence microscopy and nanoscopy studies. In order to test the permeability and specificity in live-cell labeling, we initiated our studies utilizing a two-step labeling of HaloTag-expressing cells.^10, 48^ In this protocol, U2OS cells stably expressing a vimentin-HaloTag construct^49^ were first labeled with the chloroalkane ligand HTL-BCN (Figure 4a), following an established protocol (10 μM, 30 min) to incorporate the cycloalkyne into the HaloTag, followed by the overnight labeling with the respective dyad (**1**–**12**, 200 nM). Afterwards, the cells were fixed with paraformaldehyde and imaged by confocal microscopy (Figure 4a and Figure S11). For most dyads, the labeling was specific for vimentin filaments, and showed a high contrast between before and after photoactivation. Alkoxytetrazine (**7, 8**) and secondary acrylamide (**9, 10**) dyads failed to give meaningful images, which may be attributed to cell impermeability, poor specificity, slow reaction kinetics, low chemical stability (e.g., against cellular nucleophiles), or a combination thereof.

**Figure 4.**
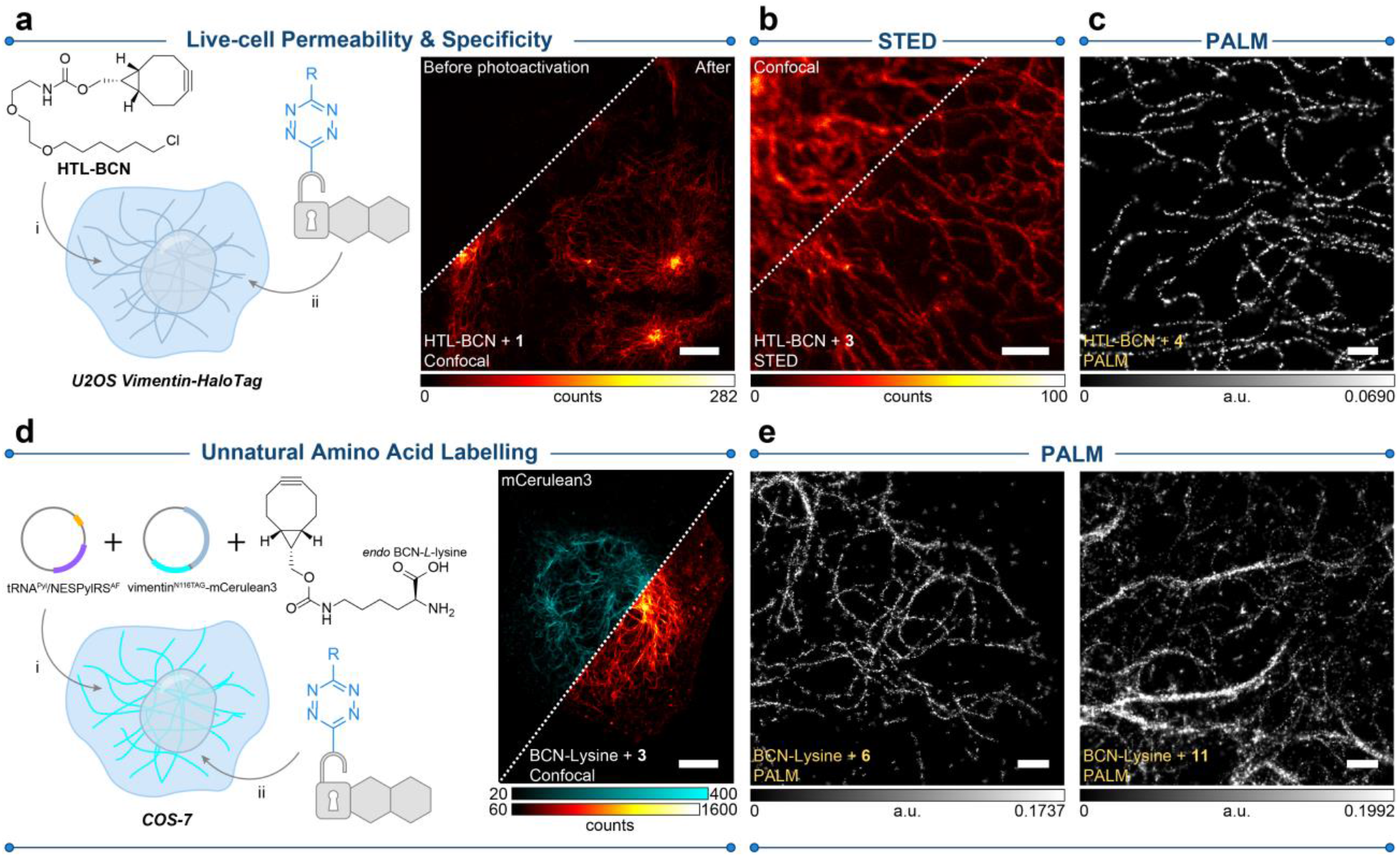
Live-cell labeling with PaX-Tz dyads for imaging. **a**, Live cell permeability and BCN-specificity were assessed by confocal imaging in U2OS cells stably expressing a vimentin-HaloTag construct in a two-step labeling strategy utilizing the HaloTag specific ligand HTL-BCN (10 μM, 30 min) and a dyad (200 nM, overnight), fixed with PFA and (e.g., **1**, right panel). Conversion to PaXCF was achieved by 405 nm illumination. **b**, STED image of cells described in **a**, labeled with compound **3**. Conversion to PaXCF was achieved with widefield illumination (AHF analysentechnik AG, 4,6-diamidino-2-phenylindole filter set F46-816). **c**, PALM image of cells described in **a**, labeled with compound **4. d**, COS-7 cells transiently expressing a vimentin-mCerulean3 fusion construct^50^ carrying a N116TAG mutation (for incorporating the UAA *endo* BCN-*L*-lysine) in vimentin.^51-52^ The plasmids were provided by the Lemke lab (EMBL, Heidelberg). See Figure S13 for detailed maps of tRNAPyl/NESPylRSAF, and vimentin^N116TAG^-mCerulean3 plasmids. Transfected cells were labeled with a dyad (500 nM, 4 h), fixed with PFA and imaged. Right: confocal image of a transfected cell showing mCerulean signal (cyan) and co-labeled with **3** (red hot). Conversion to the PaXCF was achieved by 405 nm illumination. **e**, PALM imaging of cells described in **d** labeled with compounds **6** (left) and **11** (right). Scale bars: 10 μm (**a, d**), 500 nm (**c, e**).

In our previous work, PaX derived probes (such as PaX560 conjugated antibodies and nanobodies) were compatible with STED nanoscopy, which we confirmed was also the case for clickable Tz dyad **3** utilizing depletion at 660 nm, giving results consistent with our previous report (Figure 4b). Acrylamide-derived dyads were, however, incompatible with STED. All dyads with good labelling (i.e., excluding **7**–**10**) performed remarkable well in PALM (Figure 4c and Figure S12), where we observed well-resolved structures. As expected from their photophysical properties, compounds **5, 6** and **12** bearing azetidine auxochromic groups yielded higher overall photon counts. Methyl substituted Tz derivatives (**2, 4, 6**) yielded 15–20% less photons than the unsubstituted analogues (**1, 3, 5**). We also observed that tertiary acrylamide-PaX (**11, 12**) required higher intensity 405 nm photoactivation in PALM, while also yielding more photons than their first-generation counterparts. Taken together, these results highlight the diverse applicability of PaX-Tz dyads: while the probes featuring alkene radical traps (e.g., **3**) require much less UV activation, they are STED-compatible; tertiary acrylamides **11**,**12** afford greater control in localization-based techniques. Based on these results, and the above spectroscopic characterizations, we selected compounds **11** and **12** in subsequent MINFLUX experiments (see below).

Next, COS-7 cells were prepared (Fig 4d), transiently expressing a vimentin-mCerulean3 fusion construct^50^ carrying a N116TAG mutation (for incorporating the UAA endo BCN-*L*-lysine) in the 1A coil fragment of the vimentin head domain.^51-52^ Initial staining with **3** (Figure 4d) demonstrated bright labeling of the mCerulean3-tagged vimentin filaments after photoactivation, with a minimal background that can be attributed to non-incorporated UAA, free or bound to tRNA, or unreacted PaX-Tz in lipophilic compartments. Using this strategy, cells were further imaged by PALM microscopy (Figure 4e and Figure S14) for a subset palette of our dyads.

### Minimal linkage error MINFLUX nanoscopy of vimentin filaments

To demonstrate the advantage of linkage-error-free labeling with our dyads, we turned to MINFLUX nanoscopy to visualize the vimentin filaments prepared via genetic code expansion (as described above) and stained with compound **12**. Individual fluorophores were localized by MINFLUX with a median precision of 2 nm, utilizing 100 photons per localization (Figure 5a). Measuring the average full-width half maximum (FWHM) of linearized filament segments (Figure 5b, see Figure S15 and accompanied Supporting Information for methodology) we obtained an average value of 14 nm, which was in excellent agreement with the value reported by cryogenic electron microscopy/electron tomography data,^53-56^ thus confirming the PaX-Tz labeling strategy reduced the linkage-error, in practice, to an undetectable level. To support this claim, we also recorded as a comparison MINFLUX images of the transfected cells labeled with nanobodies targeting the mCerulean3 and carrying the Tz dyad **11** attached via TCO-PEG3-maleimide coupling (Figure S16). Compound **12** failed to provide stoichiometrically-labeled nanobodies, likely due to lower aqueous solubility imparted by the azetidine groups. Also transfected and native COS-7 cells labeled by indirect immunofluorescence (using secondary antibodies bearing **12**) were imaged. The same filament thickness analysis yielded significantly larger values of 22 and ∼30 nm for the nanobody and antibody labeling, respectively. This comparison highlights the importance of the linkage errors associated with the selected labeling method. However, we also found that anti-vimentin antibodies screened failed to uniformly mark all the filaments in transfected cells (Figure S17), serendipitously revealing a second population of predominantly-untagged single filaments. While beyond the scope of this work, this result highlights the challenges of incorporating UAAs in cellular components as well as the versatile role of vimentin bundles. We further observed no difference in filament thickness by indirect immunofluorescence between transfected and wild-type COS-7 cells (Figure S18).

**Figure 5.**
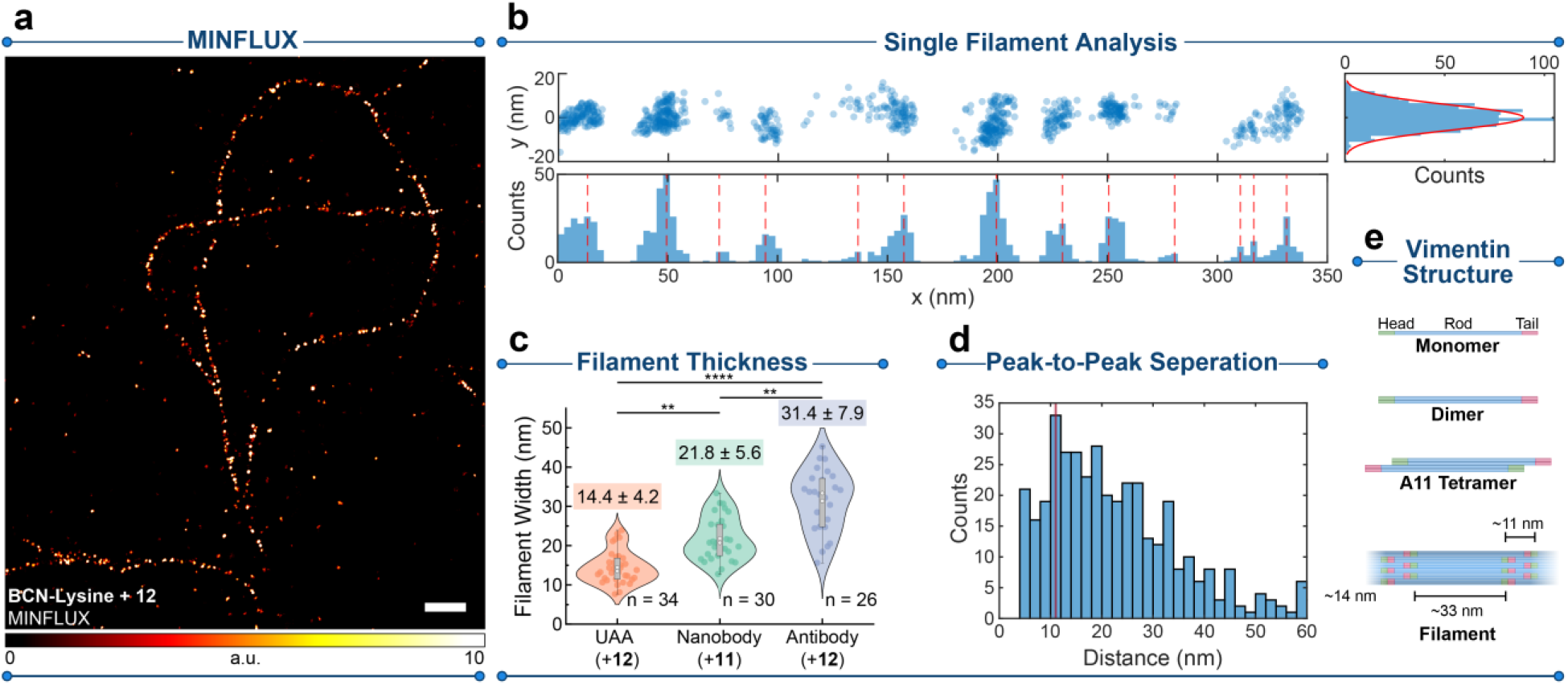
MINFLUX characterization of linkage-error-free labeling. **a**, MINFLUX image of vimentin filaments in COS-7 cells incorporating *endo*-BCN-*L*-lysine as described in Figure 3d and labeled with compound **12. b**, Single filament analysis of the linearized single filament marked in **a**; distributions of localizations along the filament length (x) and width (y) were used to determine peak-to-peak separation of localization domains and the FWHM of the filament segment, respectively. **c**, Distribution of filament thicknesses of vimentin segments labeled with different strategies, including UAA + compound **12**. anti-GFP nanobodies + compound **11**, and indirect immunofluorescence + compound **12** in transfected COS-7 cells (as described above). The distribution mean and SDs are written above or below the violin plots. **P < 0.01; ****P < 0.0001 using the Kruskal-Wallis and the Dunn method for post-hoc correction for multiple comparisons. **d**, Peak-to-peak separations of localization domains in vimentin filaments labeled with UAA. **e**, Structure of the vimentin according to recent literature with our obtained values shown.^57^ Scale bar: 200 nm.

The direct incorporation of the fluorophores into the head domain of the vimentin monomers can additionally reveal information on the multimeric structure of the mature filaments. By measuring the peak-to-peak distance of localization clusters along the filament length (Figure 5d) we observed an abundance of pairwise distances of 11 nm, consistent with previous observations on the structure of vimentin unit length filaments (ULFs) utilizing C and N-terminal tagging strategies,^57^ where values of ∼39 nm and ∼10 nm are expected for separations of head domains within single ULFs and between annealed ULFs in matured filaments, respectively (Figure 5e). However, in previous work these values could not be directly observed *in cellulo* due to limitations to resolution and linkage error, further highlighting the power of our strategy. In our case, direct tagging of N116 shifts the fluorophore position inwards, increasing the observed value, but low fluorophore density and ULF orientations further contribute to measurement errors. The additional peak at 33 nm may thus be due to the ULF head domain separations. While beyond the scope of this work, further improvements to UAA incorporation^26, 58^ as well as 3D imaging modalities can afford an even more complete visualization the of vimentin substructure.

## CONCLUSION

We show, for the first time, how the excited state quenching inherent to tetrazine dyads can be additionally utilized to impart chemical control over the activation process of fluorophores for next generation super-resolution fluorescence microscopy. We have prepared a family of highly compact, biorthogonal, live-cell compatible, clickable and photoactivatable probes for UAAs introduced by genetic code expansion, that can be applied in STED, PALM and MINFLUX nanoscopies. Combined with the ultimate resolution of MINFLUX, we visualized the substructure of vimentin filaments incorporating an UAA in the head domain of the filament monomers *in cellulo* and with molecular-scale precision. Our work demonstrates the combined potential of linkage-error-free labeling with appropriately designed caging-group-free photoactivatable fluorophores.

## SUPPORTING INFORMATION

Additional experimental details, materials, methods, and characterization data for all new compounds (PDF).

## ACKNOWLEDGEMENTS

The research was funded by Bundesministerium für Bildung und Forschung (German Federal Ministry of Education and Research), Project No. 13N14122 “3D Nano Life Cell” (to S.W.H.). R.L. is grateful to the Max Planck Society for a Nobel Laureate Fellowship. We thank Prof. S. Jakobs (Max Planck Institute for Multidisciplinary Sciences, University of Göttingen) for providing the U2OS-Vim-Halo. We thank Prof. E. Lemke (Johannes Gutenberg University, Mainz) and the EMBL for providing the pEvolv_tRNAPyl/PylRSAF, CMV_NES-PylRS(AF)_hU6tRNAPyl, and Vimentin(N116TAG)-mCerulean3 plasmids. We thank the staff of Mass Spectrometry Core Facility (Max Planck Institute for Medical Research) for recording mass spectra of small molecules, M. Tarnawski (Max Planck Institute for Medical Research, Protein Expression and Characterization Facility) for the GFP protein, the Optical Microscopy Facility (MPI MR) for the use of their fluorescence microscopes, and M. Remmel (MPI MR) for the use of a custom-built PALM microscope.

## NOTES

The authors declare the following competing financial interest(s): R.L., M.L.B., and A.N.B. are co-inventors of a patent application covering the photoactivatable dyes of this work, filed by the Max Planck Society. S.W.H. owns shares of Abberior GmbH and Abberior Instruments GmbH whose dyes and MINFLUX microscope, respectively, have also been used in this study.

## REFERENCES

1. Sahl, S. J.; Hell, S. W.; Jakobs, S., Fluorescence nanoscopy in cell biology. Nat. Rev. Mol. Cell Biol. 2017, 18 (11), 685–701.

2. Helmerich, D. A.; Beliu, G.; Taban, D.; Meub, M.; Streit, M.; Kuhlemann, A.; Doose, S.; Sauer, M., Photoswitching fingerprint analysis bypasses the 10-nm resolution barrier. Nat. Methods 2022, 19 (8), 986–994.

3. Gidi, Y.; Payne, L.; Glembockyte, V.; Michie, M. S.; Schnermann, M. J.; Cosa, G., Unifying Mechanism for Thiol-Induced Photoswitching and Photostability of Cyanine Dyes. J. Am. Chem. Soc. 2020, 142 (29), 12681–12689.

4. Dempsey, G. T.; Vaughan, J. C.; Chen, K. H.; Bates, M.; Zhuang, X., Evaluation of fluorophores for optimal performance in localization-based super-resolution imaging. Nat. Methods 2011, 8 (12), 1027–36.

5. Beliu, G.; Kurz, A. J.; Kuhlemann, A. C.; Behringer-Pliess, L.; Meub, M.; Wolf, N.; Seibel, J.; Shi, Z.-D.; Schnermann, M.; Grimm, J. B.; Lavis, L. D.; Doose, S.; Sauer, M., Bioorthogonal labeling with tetrazine-dyes for super-resolution microscopy. Communications Biology 2019, 2 (1), 261.

6. Uno, S. N.; Kamiya, M.; Morozumi, A.; Urano, Y., A green-light-emitting, spontaneously blinking fluorophore based on intramolecular spirocyclization for dual-colour super-resolution imaging. Chem. Commun. (Camb.) 2017, 54 (1), 102–105.

7. Morozumi, A.; Kamiya, M.; Uno, S. N.; Umezawa, K.; Kojima, R.; Yoshihara, T.; Tobita, S.; Urano, Y., Spontaneously Blinking Fluorophores Based on Nucleophilic Addition/Dissociation of Intracellular Glutathione for Live-Cell Super-resolution Imaging. J. Am. Chem. Soc. 2020, 142 (21), 9625–9633.

8. Gerasimaite, R. T.; Bucevicius, J.; Kiszka, K. A.; Schnorrenberg, S.; Kostiuk, G.; Koenen, T.; Lukinavicius, G., Blinking Fluorescent Probes for Tubulin Nanoscopy in Living and Fixed Cells. ACS Chem. Biol. 2021, 16 (11), 2130–2136.

9. Werther, P.; Yserentant, K.; Braun, F.; Kaltwasser, N.; Popp, C.; Baalmann, M.; Herten, D. P.; Wombacher, R., Live-Cell Localization Microscopy with a Fluorogenic and Self-Blinking Tetrazine Probe. Angew Chem Int Ed Engl 2020, 59 (2), 804–810.

10. Werther, P.; Yserentant, K.; Braun, F.; Grussmayer, K.; Navikas, V.; Yu, M.; Zhang, Z.; Ziegler, M. J.; Mayer, C.; Gralak, A. J.; Busch, M.; Chi, W.; Rominger, F.; Radenovic, A.; Liu, X.; Lemke, E. A.; Buckup, T.; Herten, D. P.; Wombacher, R., Bio-orthogonal Red and Far-Red Fluorogenic Probes for Wash-Free Live-Cell and Super-resolution Microscopy. ACS Cent Sci 2021, 7 (9), 1561–1571.

11. Remmel, M.; Scheiderer, L.; Butkevich, A. N.; Bossi, M. L.; Hell, S. W., Accelerated MINFLUX Nanoscopy, through Spontaneously Fast-Blinking Fluorophores. Small 2023, 19 (12), e2206026.

12. Zheng, Y.; Ye, Z.; Zhang, X.; Xiao, Y., Recruiting Rate Determines the Blinking Propensity of Rhodamine Fluorophores for Super-Resolution Imaging. Journal of the American Chemical Society 2023, 145 (9), 5125–5133.

13. Roubinet, B.; Weber, M.; Shojaei, H.; Bates, M.; Bossi, M. L.; Belov, V. N.; Irie, M.; Hell, S. W., Fluorescent Photoswitchable Diarylethenes for Biolabeling and Single-Molecule Localization Microscopies with Optical Superresolution. J. Am. Chem. Soc. 2017, 139 (19), 6611–6620.

14. Uno, K.; Aktalay, A.; Bossi, M. L.; Irie, M.; Belov, V. N.; Hell, S. W., Turn-on mode diarylethenes for bioconjugation and fluorescence microscopy of cellular structures. Proc. Natl. Acad. Sci. U. S. A. 2021, 118 (14), e2100165118.

15. Hauke, S.; von Appen, A.; Quidwai, T.; Ries, J.; Wombacher, R., Specific protein labeling with caged fluorophores for dual-color imaging and super-resolution microscopy in living cells. Chemical Science 2017, 8 (1), 559–566.

16. Belov, V. N.; Wurm, C. A.; Boyarskiy, V. P.; Jakobs, S.; Hell, S. W., Rhodamines NN: a novel class of caged fluorescent dyes. Angew Chem Int Ed Engl 2010, 49 (20), 3520–3.

17. Grimm, J. B.; Klein, T.; Kopek, B. G.; Shtengel, G.; Hess, H. F.; Sauer, M.; Lavis, L. D., Synthesis of a Far-Red Photoactivatable Silicon-Containing Rhodamine for Super-Resolution Microscopy. Angew Chem Int Ed Engl 2016, 55 (5), 1723–7.

18. Balzarotti, F.; Eilers, Y.; Gwosch, K. C.; Gynna, A. H.; Westphal, V.; Stefani, F. D.; Elf, J.; Hell, S. W., Nanometer resolution imaging and tracking of fluorescent molecules with minimal photon fluxes. Science 2017, 355 (6325), 606–612.

19. Gwosch, K. C.; Pape, J. K.; Balzarotti, F.; Hoess, P.; Ellenberg, J.; Ries, J.; Hell, S. W., MINFLUX nanoscopy delivers 3D multicolor nanometer resolution in cells. Nat. Methods 2020, 17 (2), 217–224.

20. Wolff, J. O.; Scheiderer, L.; Engelhardt, T.; Engelhardt, J.; Matthias, J.; Hell, S. W., MINFLUX dissects the unimpeded walking of kinesin-1. Science 2023, 379 (6636), 1004–1010.

21. Deguchi, T.; Iwanski, M. K.; Schentarra, E. M.; Heidebrecht, C.; Schmidt, L.; Heck, J.; Weihs, T.; Schnorrenberg, S.; Hoess, P.; Liu, S.; Chevyreva, V.; Noh, K. M.; Kapitein, L. C.; Ries, J., Direct observation of motor protein stepping in living cells using MINFLUX. Science 2023, 379 (6636), 1010–1015.

22. Weber, M.; Leutenegger, M.; Stoldt, S.; Jakobs, S.; Mihaila, T. S.; Butkevich, A. N.; Hell, S. W., MINSTED fluorescence localization and nanoscopy. Nat Photonics 2021, 15 (5), 361–366.

23. Weber, M.; von der Emde, H.; Leutenegger, M.; Gunkel, P.; Sambandan, S.; Khan, T. A.; Keller-Findeisen, J.; Cordes, V. C.; Hell, S. W., MINSTED nanoscopy enters the Angstrom localization range. Nat. Biotechnol. 2023, 41 (4), 569–576.

24. Liu, S.; Hoess, P.; Ries, J., Super-Resolution Microscopy for Structural Cell Biology. Annu Rev Biophys 2022, 51, 301–326.

25. Liu, S.; Hoess, P.; Ries, J., Super-Resolution Microscopy for Structural Cell Biology. Annual Review of Biophysics 2022, 51 (1), 301–326.

26. Mihaila, T. S.; Bate, C.; Ostersehlt, L. M.; Pape, J. K.; Keller-Findeisen, J.; Sahl, S. J.; Hell, S. W., Enhanced incorporation of subnanometer tags into cellular proteins for fluorescence nanoscopy via optimized genetic code expansion. Proc. Natl. Acad. Sci. U. S. A. 2022, 119 (29), e2201861119.

27. Lee, S.; Kim, J.; Koh, M., Recent Advances in Fluorescence Imaging by Genetically Encoded Non-canonical Amino Acids. J. Mol. Biol. 2022, 434 (8), 167248.

28. Oliveira, B. L.; Guo, Z.; Bernardes, G. J. L., Inverse electron demand Diels-Alder reactions in chemical biology. Chem. Soc. Rev. 2017, 46 (16), 4895–4950.

29. Chi, W.; Huang, L.; Wang, C.; Tan, D.; Xu, Z.; Liu, X., A unified fluorescence quenching mechanism of tetrazine-based fluorogenic dyes: energy transfer to a dark state. Materials Chemistry Frontiers 2021, 5 (18), 7012–7021.

30. Kozma, E.; Kele, P., Fluorogenic probes for super-resolution microscopy. Org. Biomol. Chem. 2019, 17 (2), 215–233.

31. Wieczorek, A.; Werther, P.; Euchner, J.; Wombacher, R., Green-to far-red-emitting fluorogenic tetrazine probes – synthetic access and no-wash protein imaging inside living cells. Chemical Science 2017, 8 (2), 1506–1510.

32. Lee, Y.; Cho, W.; Sung, J.; Kim, E.; Park, S. B., Monochromophoric Design Strategy for Tetrazine-Based Colorful Bioorthogonal Probes with a Single Fluorescent Core Skeleton. J. Am. Chem. Soc. 2018, 140 (3), 974–983.

33. Knorr, G.; Kozma, E.; Schaart, J. M.; Nemeth, K.; Torok, G.; Kele, P., Bioorthogonally Applicable Fluorogenic Cyanine-Tetrazines for No-Wash Super-Resolution Imaging. Bioconjug. Chem. 2018, 29 (4), 1312–1318.

34. Lincoln, R.; Bossi, M. L.; Remmel, M.; D’Este, E.; Butkevich, A. N.; Hell, S. W., A general design of caging-group-free photoactivatable fluorophores for live-cell nanoscopy. Nat. Chem. 2022, 14 (9), 1013–1020.

35. Likhotkin, I.; Lincoln, R.; Bossi, M. L.; Butkevich, A. N.; Hell, S. W., Photoactivatable Large Stokes Shift Fluorophores for Multicolor Nanoscopy. J. Am. Chem. Soc. 2023, 145 (3), 1530–1534.

36. Knall, A. C.; Slugovc, C., Inverse electron demand Diels-Alder (iEDDA)-initiated conjugation: a (high) potential click chemistry scheme. Chem. Soc. Rev. 2013, 42 (12), 5131–42.

37. Svatunek, D.; Wilkovitsch, M.; Hartmann, L.; Houk, K. N.; Mikula, H., Uncovering the Key Role of Distortion in Bioorthogonal Tetrazine Tools That Defy the Reactivity/Stability Trade-Off. J. Am. Chem. Soc. 2022, 144 (18), 8171–8177.

38. Chi, W. J.; Huang, L.; Wang, C.; Tan, D.; Xu, Z. C.; Liu, X. G., A unified fluorescence quenching mechanism of tetrazine-based fluorogenic dyes: energy transfer to a dark state. Materials Chemistry Frontiers 2021, 5 (18), 7012–7021.

39. Schnell, S. D.; Hoff, L. V.; Panchagnula, A.; Wurzenberger, M. H. H.; Klapotke, T. M.; Sieber, S.; Linden, A.; Gademann, K., 3-Bromotetrazine: labelling of macromolecules via monosubstituted bifunctional s-tetrazines. Chem. Sci. 2020, 11 (11), 3042–3047.

40. Butkevich, A. N.; Bossi, M. L.; Lukinavicius, G.; Hell, S. W., Triarylmethane Fluorophores Resistant to Oxidative Photobluing. J. Am. Chem. Soc. 2019, 141 (2), 981–989.

41. Grimm, J. B.; Xie, L.; Casler, J. C.; Patel, R.; Tkachuk, A. N.; Falco, N.; Choi, H.; Lippincott-Schwartz, J.; Brown, T. A.; Glick, B. S.; Liu, Z.; Lavis, L. D., A General Method to Improve Fluorophores Using Deuterated Auxochromes. JACS Au 2021, 1 (5), 690–696.

42. Li, X.; Wang, Y. Preparation of tetrazine-based fluorescent probes for detection of superoxide anion. CN115583920, 2023.

43. Plass, T.; Milles, S.; Koehler, C.; Schultz, C.; Lemke, E. A., Genetically encoded copper-free click chemistry. Angew Chem Int Ed Engl 2011, 50 (17), 3878–81.

44. Borrmann, A.; Milles, S.; Plass, T.; Dommerholt, J.; Verkade, J. M.; Wiessler, M.; Schultz, C.; van Hest, J. C.; van Delft, F. L.; Lemke, E. A., Genetic encoding of a bicyclo[6.1.0]nonyne-charged amino acid enables fast cellular protein imaging by metal-free ligation. ChemBioChem 2012, 13 (14), 2094–9.

45. Lang, K.; Davis, L.; Wallace, S.; Mahesh, M.; Cox, D. J.; Blackman, M. L.; Fox, J. M.; Chin, J. W., Genetic Encoding of bicyclononynes and trans-cyclooctenes for site-specific protein labeling in vitro and in live mammalian cells via rapid fluorogenic Diels-Alder reactions. J. Am. Chem. Soc. 2012, 134 (25), 10317–20.

46. Nikic, I.; Plass, T.; Schraidt, O.; Szymanski, J.; Briggs, J. A.; Schultz, C.; Lemke, E. A., Minimal tags for rapid dual-color live-cell labeling and super-resolution microscopy. Angew Chem Int Ed Engl 2014, 53 (8), 2245–9.

47. Schlesinger, O.; Chemla, Y.; Heltberg, M.; Ozer, E.; Marshall, R.; Noireaux, V.; Jensen, M. H.; Alfonta, L., Tuning of Recombinant Protein Expression in Escherichia coli by Manipulating Transcription, Translation Initiation Rates, and Incorporation of Noncanonical Amino Acids. ACS Synth Biol 2017, 6 (6), 1076–1085.

48. Murrey, H. E.; Judkins, J. C.; Am Ende, C. W.; Ballard, T. E.; Fang, Y.; Riccardi, K.; Di, L.; Guilmette, E. R.; Schwartz, J. W.; Fox, J. M.; Johnson, D. S., Systematic Evaluation of Bioorthogonal Reactions in Live Cells with Clickable HaloTag Ligands: Implications for Intracellular Imaging. J. Am. Chem. Soc. 2015, 137 (35), 11461–75.

49. Butkevich, A. N.; Ta, H.; Ratz, M.; Stoldt, S.; Jakobs, S.; Belov, V. N.; Hell, S. W., Two-Color 810 nm STED Nanoscopy of Living Cells with Endogenous SNAP-Tagged Fusion Proteins. ACS Chem. Biol. 2018, 13 (2), 475–480.

50. Markwardt, M. L.; Kremers, G. J.; Kraft, C. A.; Ray, K.; Cranfill, P. J.; Wilson, K. A.; Day, R. N.; Wachter, R. M.; Davidson, M. W.; Rizzo, M. A., An improved cerulean fluorescent protein with enhanced brightness and reduced reversible photoswitching. PLoS One 2011, 6 (3), e17896.

51. Nikic, I.; Estrada Girona, G.; Kang, J. H.; Paci, G.; Mikhaleva, S.; Koehler, C.; Shymanska, N. V.; Ventura Santos, C.; Spitz, D.; Lemke, E. A., Debugging Eukaryotic Genetic Code Expansion for Site-Specific Click-PAINT Super-Resolution Microscopy. Angew Chem Int Ed Engl 2016, 55 (52), 16172–16176.

52. Gregor, C.; Grimm, F.; Rehman, J.; Wurm, C. A.; Egner, A., Two-color live-cell STED nanoscopy by click labeling with cell-permeable fluorophores. bioRxiv 2022, 2022.09.11.507450.

53. Mucke, N.; Wedig, T.; Burer, A.; Marekov, L. N.; Steinert, P. M.; Langowski, J.; Aebi, U.; Herrmann, H., Molecular and biophysical characterization of assembly-starter units of human vimentin. J. Mol. Biol. 2004, 340 (1), 97–114.

54. Goldie, K. N.; Wedig, T.; Mitra, A. K.; Aebi, U.; Herrmann, H.; Hoenger, A., Dissecting the 3-D structure of vimentin intermediate filaments by cryo-electron tomography. J. Struct. Biol. 2007, 158 (3), 378–85.

55. Chernyatina, A. A.; Nicolet, S.; Aebi, U.; Herrmann, H.; Strelkov, S. V., Atomic structure of the vimentin central alpha-helical domain and its implications for intermediate filament assembly. Proc. Natl. Acad. Sci. U. S. A. 2012, 109 (34), 13620–5.

56. Eibauer, M.; Weber, M. S.; Turgay, Y.; Sivagurunathan, S.; Goldman, R. D.; Medalia, O., The molecular architecture of vimentin filaments. bioRxiv 2021, 2021.07.15.452584.

57. Nunes Vicente, F.; Lelek, M.; Tinevez, J. Y.; Tran, Q. D.; Pehau-Arnaudet, G.; Zimmer, C.; Etienne-Manneville, S.; Giannone, G.; Leduc, C., Molecular organization and mechanics of single vimentin filaments revealed by super-resolution imaging. Sci Adv 2022, 8 (8), eabm2696.

58. Reinkemeier, C. D.; Girona, G. E.; Lemke, E. A., Designer membraneless organelles enable codon reassignment of selected mRNAs in eukaryotes. Science 2019, 363 (6434), eaaw2644.

