## Supporting Information for "Bioorthogonal caging-group-free photoactivatable probes for minimal-linkage-error nanoscopy"

***This section includes:***

Compound 10. 45
Compound S10. 46
Compound 11. 47
Compound S12. 48
Compound S13. 49
Compound S14. 50
Compound 12. 51
SUPPLEMENTARY REFERENCES 52
NMR SPECTRA 55
Compound 1 $^{1}H$ 55
Compound 1 $^{13}C$ 56
Compound 2 $^{1}H$ 57
Compound 2 $^{13}C$ 58
Compound 3 $^{1}H$ 59
Compound 3 $^{13}C$ 60
Compound 4 $^{1}H$ 61
Compound 4 $^{13}C$ 62
Compound 5 $^{1}H$ 63
Compound 5 $^{13}C$ 64
Compound 6 $^{1}H$ 65
Compound 6 $^{13}C$ 66
Compound S6 $^{1}H$ 67
Compound S6 $^{13}C$ 68
Compound S7 $^{1}H$ 69
Compound S7 $^{13}C$ 70
Compound S8 $^{1}H$ 71
Compound S8 $^{13}C$ 72
Compound S9 $^{1}H$ 73
Compound S9 $^{13}C$ 74
Compound 10 $^{1}H$ 75
Compound 10 $^{13}C$ 76
Compound S10 $^{1}H$ 77
Compound S10 $^{13}C$ 78
Compound S13 $^{1}H$ 79

### SUPPLEMENTARY FIGURES

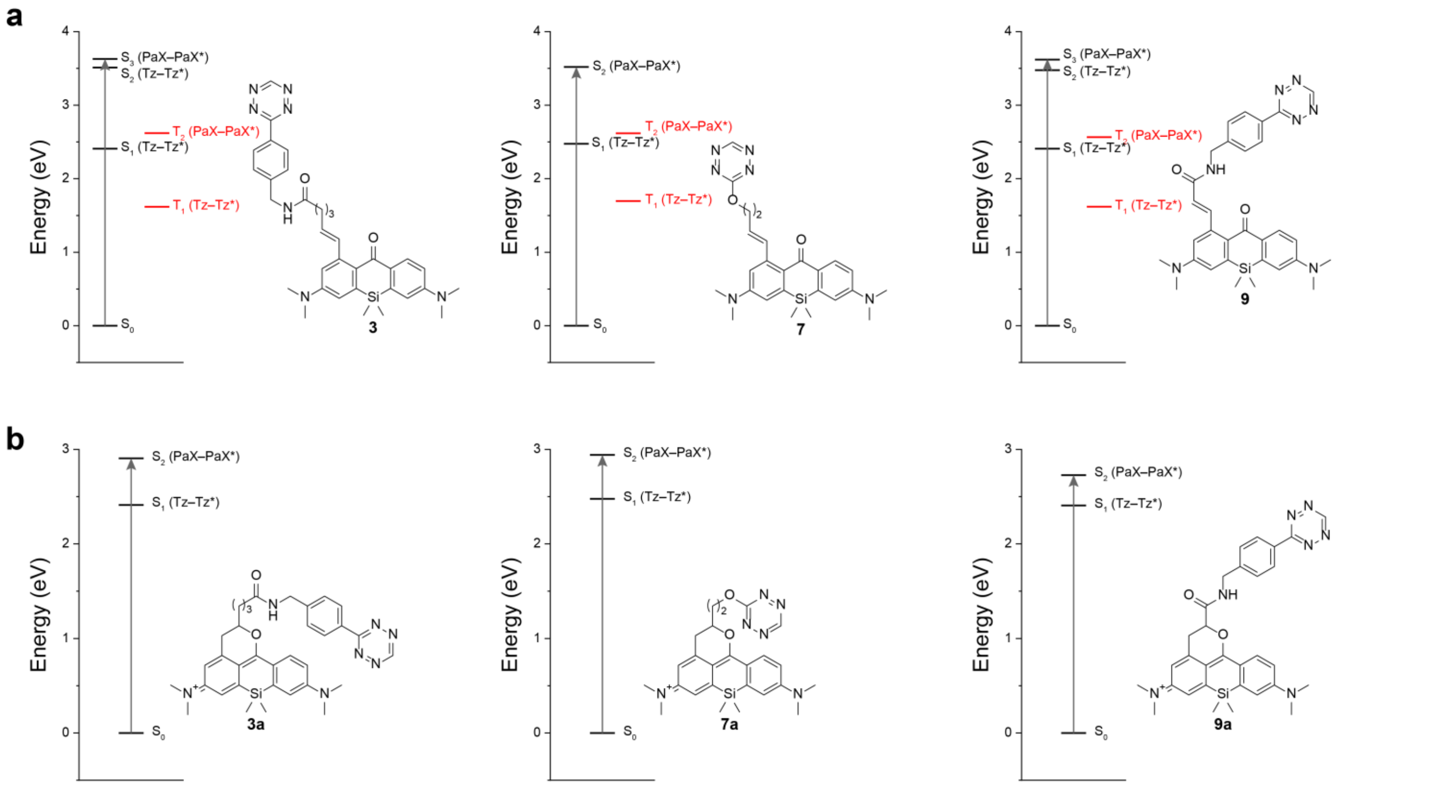

**Figure S1. a–b,** Time dependent DFT calculation of Franck-Condon transitions at the wB97XD/Def2SVP level with applied SMD model solvation of water for compounds **3**, **7** and **9** (a) and respective ‘closed-form’ structures **3a**, **7a**, and **9a** (b). Based on the orbital contributions to the excited state, the states are assigned to be centered on the tetrazine (Tz–Tz*) or the xanthone (PaX–Pax*). Tz–Tz* states with lower energy than PaX–Pax* suggests possible quenching of the PaX photoactivation (in a) or ‘closed-form’ fluorescence (in b).

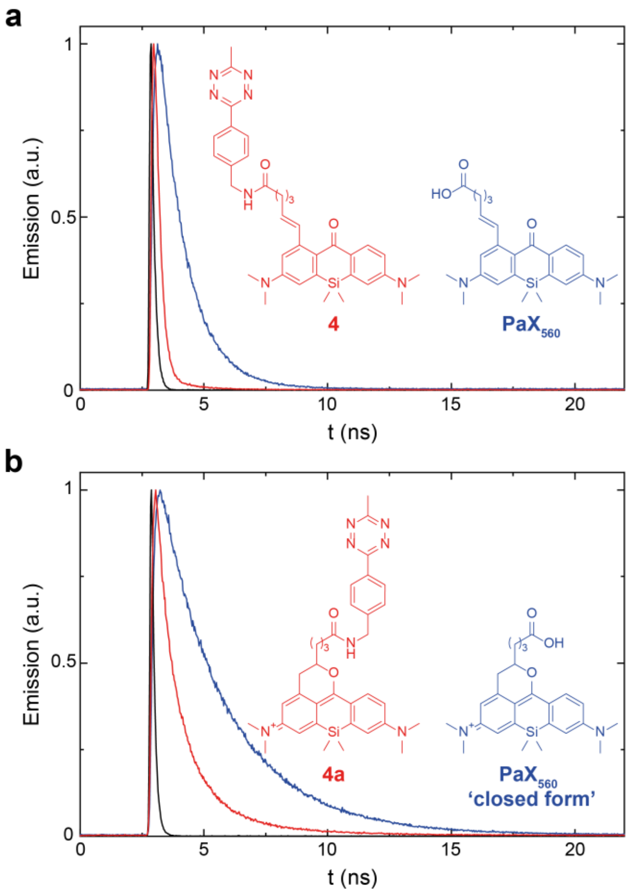

**Figure S2. a**, Fluorescence decay traces in methanol for the PaX-Tz dyad **4** (red) and the free acid of $PaX_{560}$ (blue) recorded at 490 nm ($\lambda_{ex}$ = 375 nm). **b**, Fluorescence decay traces in methanol for the 'closed form' PaX-Tz dyad 4a (red) and the 'closed form' of $PaX_{560}$ (blue) measured at 585 nm ($\lambda_{ex}$ = 375 nm) after partial activation with 405 nm. The instrument response function is shown in black.

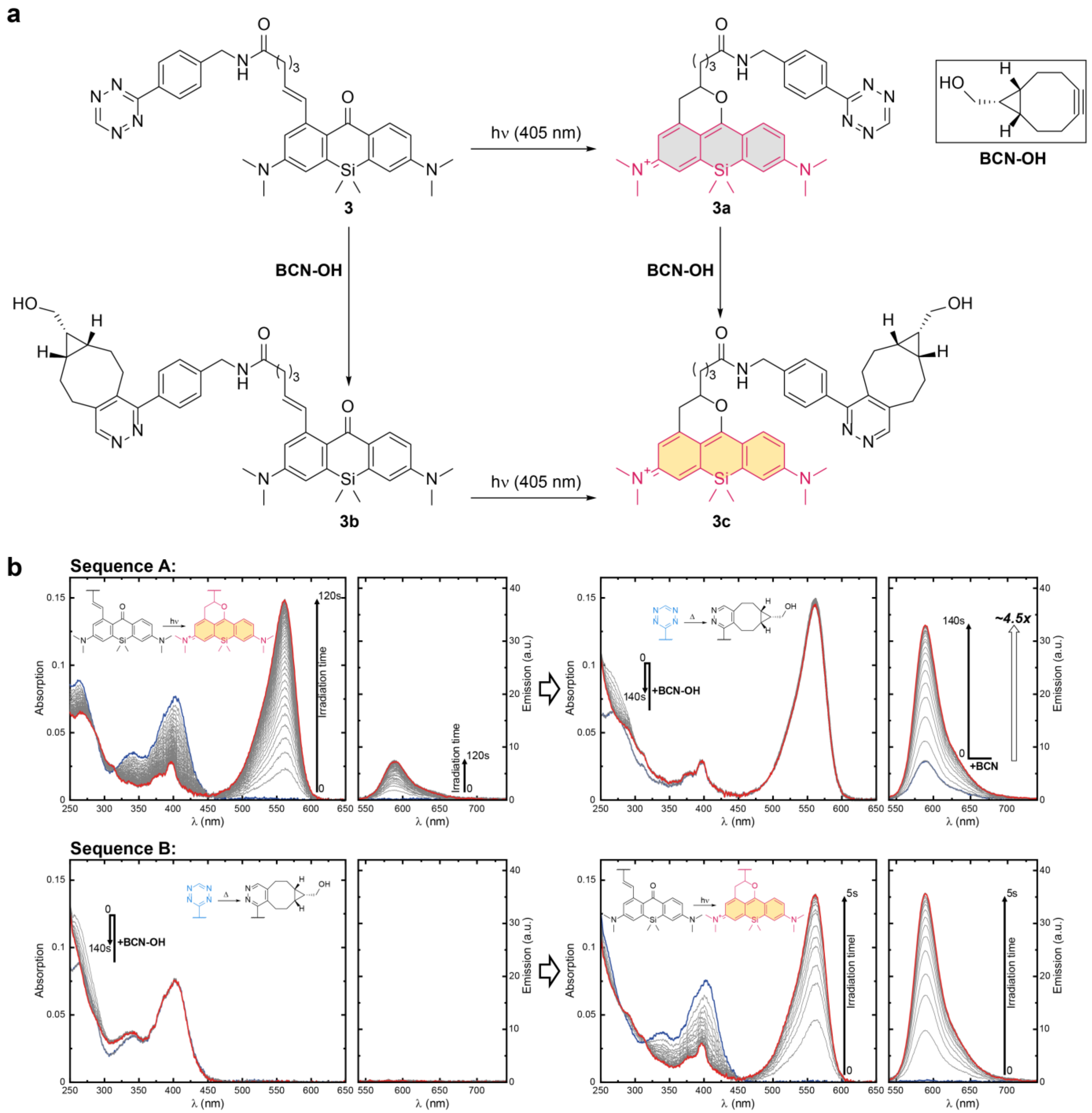

**Figure S3. a**, Chemical structures of species formed from compound **3** during Sequence A (irradiation first, then BCN-OH; clockwise from **3**) and Sequence B (BCN-OH first, then irradiation; counter clockwise from **3**). **b**, Temporal evolution of the absorption and fluorescence spectra of **3** in methanol during Sequence A (irradiation first, then BCN-OH; top row) and B (BCN-OH first, then irradiation; bottom row), $\lambda_{act} = 405$ nm.

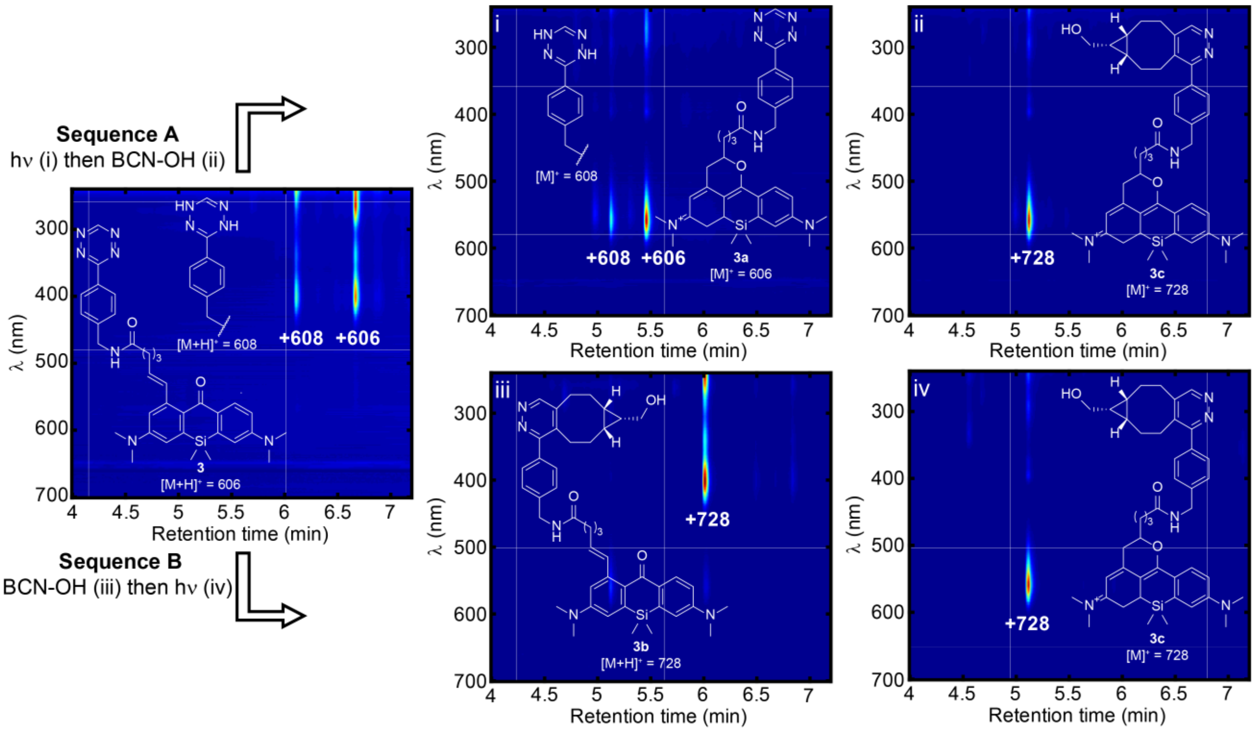

**Figure S4.** LC-MS chromatograms of the reaction mixtures of PaX **3** before and after the two stages of Sequence A (i and ii) and B (iii and iv), and the chemical structures of the major products detected.

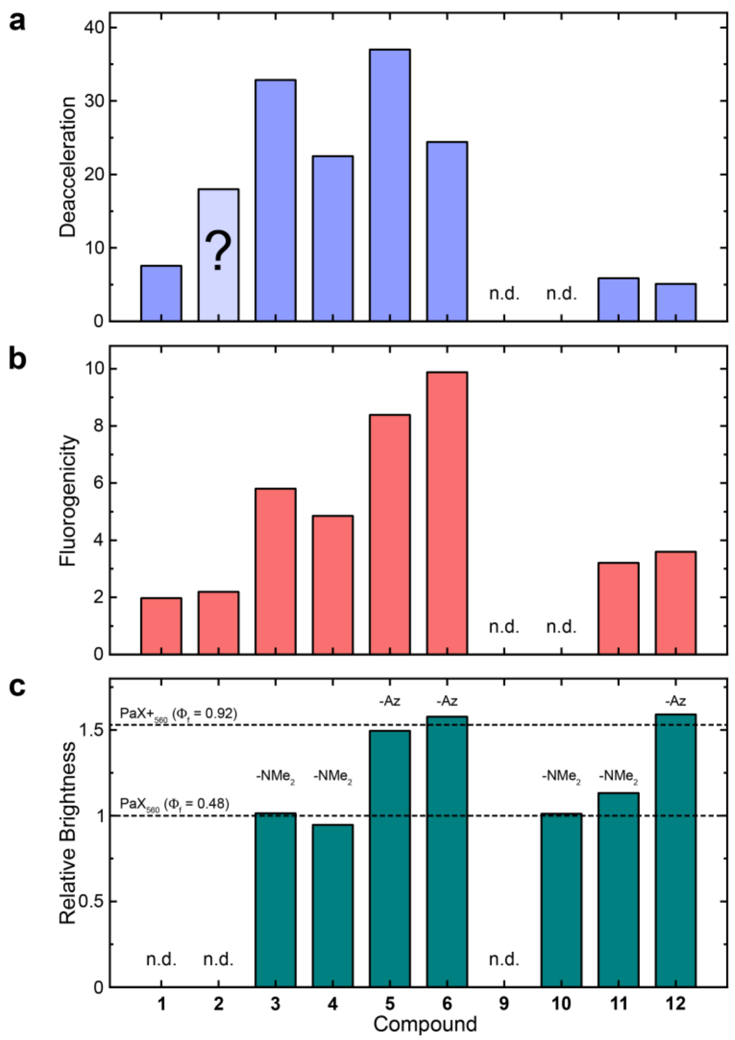

**Figure S5. a**, Effect of Tz moiety on the photoactivation rate (i.e. deacceleration) in the Tz dyads calculated from that ratio of photoactivation speeds from Sequence B vs Sequence A for each dyad. The ratio could not be determined for compounds **9** and **10**. The question mark denotes ambiguity in the value for compound **2**, due to photobleaching during Sequence A. **b**, Effect of Tz moiety on the fluorescence emission of the closed-form structures (i.e. fluorogenicity) calculated from that ratio of quantum yields from Sequence B vs Sequence A for each dyad. The ratio could not be determined for compounds 9 and 10. **c**, Relative brightness for the final pyridazine product of sequence A/B for each dyad compared to the free acids $PaX_{560}$ and $PaX+_{560}$ showing the pyrazines are equally bright to the free carboxylic acids, and the relative brightness is determined by the dimethylamino or azetidine groups.

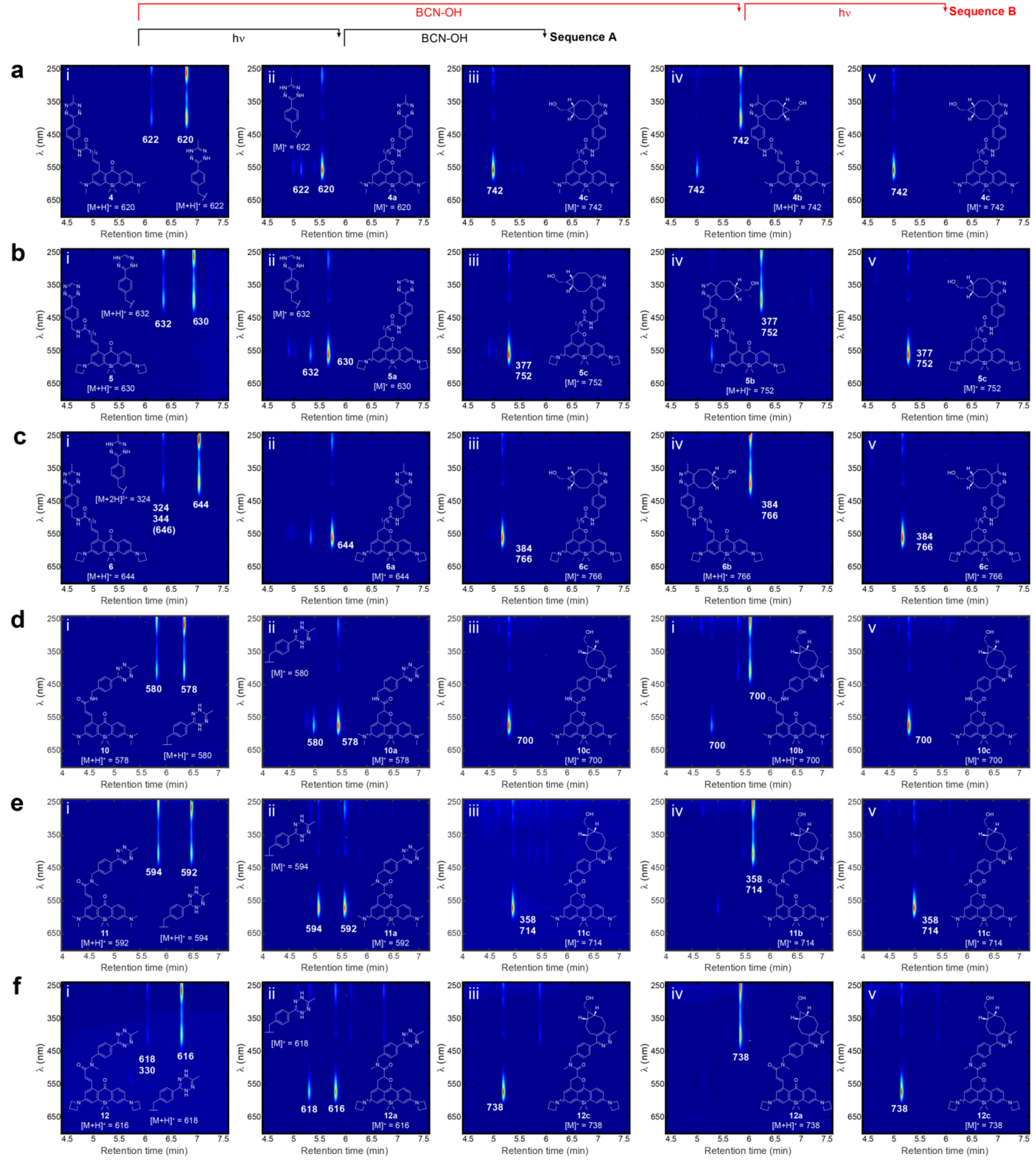

**Figure S6.** LC-MS chromatograms of the reaction mixtures of PaX-Tz 4 (**a**), 5 (**b**), 6 (**c**), 10 (**d**), 11 (**e**), and 12 (**f**) before (i) and after the two stages of Sequence A (ii and iii) and B (iv and v), and the chemical structures of the major products detected.

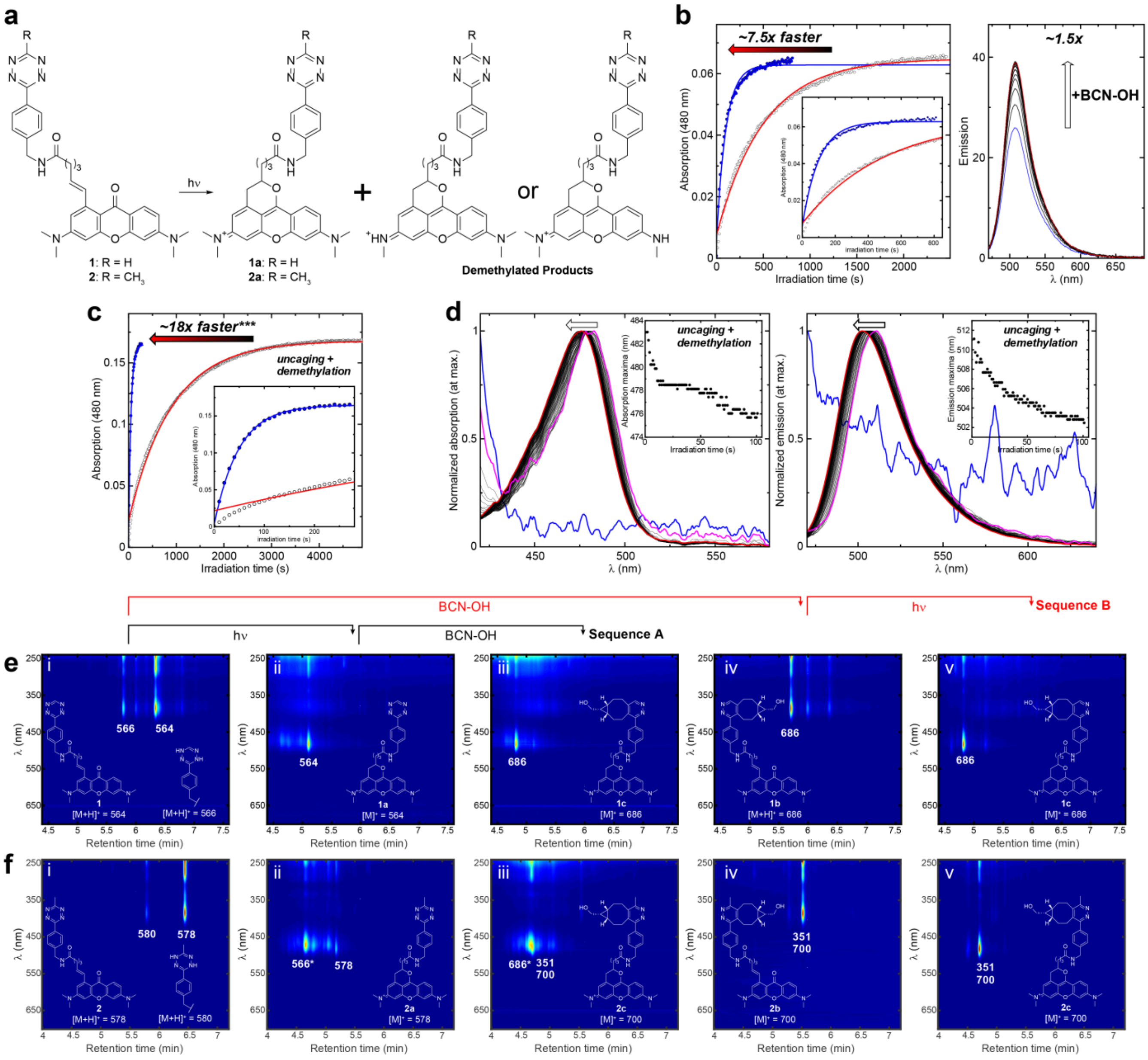

**Figure S7.** Photochemical behavior of PaX-Tz dyads derived from PaX$_{480}$. **a**, Proposed structures of photoblueing products arising from demethylation of the dye core. **b**, Temporal evolution of the absorption of the closed-form of **1** in methanol during irradiation in Sequence A (forming **1a**, white circles, red line) and B (BCN-OH first, then irradiation; forming **1c,** blue circles, blue line). The right plot shows the fluorogenicity of **1a** upon addition of BCN-OH. **c**, Temporal evolution of the absorption of **2** in methanol during irradiation in Sequence A (forming **2a**, white circles, red line) and B (BCN-OH first, then irradiation to form **2c**, blue circles, blue line). **d**, Normalized emission spectra of **2** during irradiation (Sequence A), showing a blue-shift in the emission maxima upon prolonged irradiation. **e**,**f**, LC-MS chromatograms of the reaction mixtures of PaX-Tz **1** (**e**), **2** (**f**), before (i) and after the two stages of Sequence A (ii and iii) and B (iv and v), and the chemical structures of the major products detected.

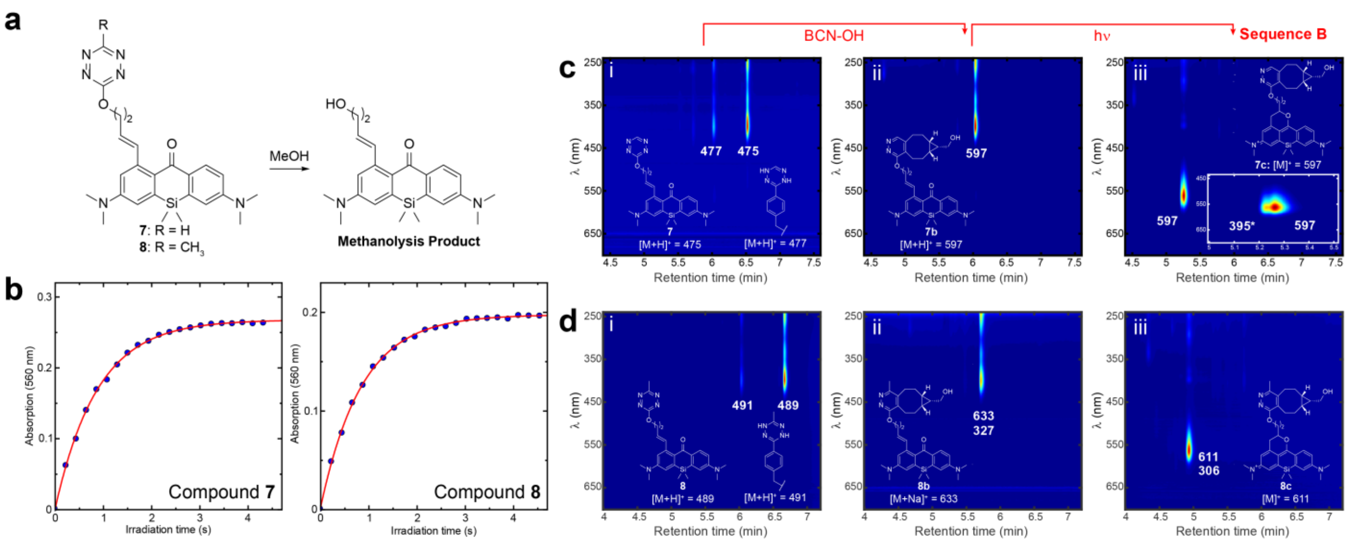

**Figure S8.** Photochemical behavior of PaX-Tz dyads featuring an ether linkage. **a**, Proposed structure of the methanolysis product. **b**, Temporal evolution of the absorption of closed-form structures (**7c** and **8c**) in methanol during irradiation of **7b** and **8b** in Sequence B. **c,d,** LC-MS chromatograms of the reaction mixtures of PaX-Tz **7** (**c**) and **8** (**d**), before (i) and after the two stages of Sequence B (ii and iii), and the chemical structures of the major products detected.

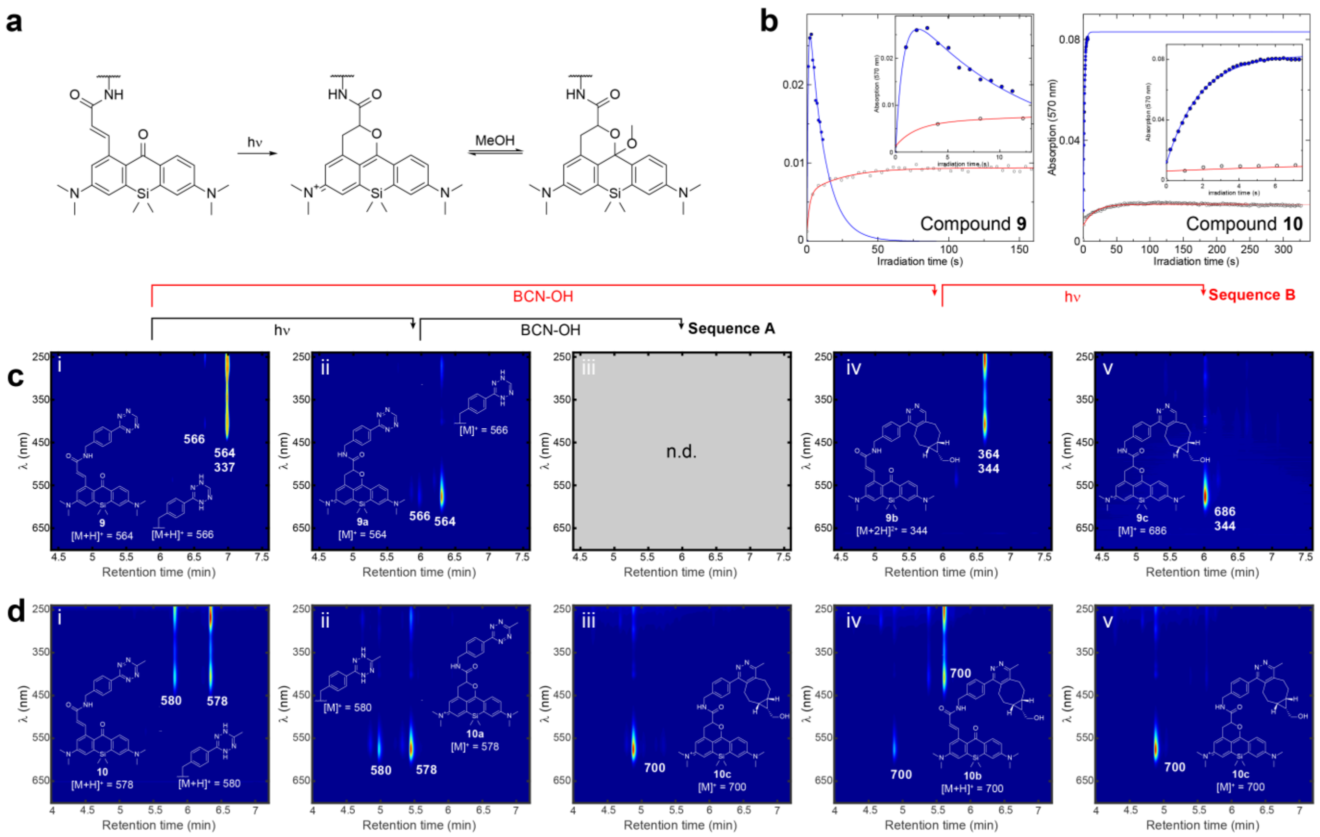

**Figure S9.** Photochemical behavior of PaX-Tz dyads featuring a secondary acrylamide linkage. **a**, Proposed structure of the methanol adduct formed after photolysis. **b**, Temporal evolution of the absorption of the closed-form products of **9** and **10** in methanol, during the irradiation in Sequence A (forming **9a** or **10a**, white circles, red line) and B (BCN-OH first, then irradiation; forming **9c** or **10c**, blue circles, blue line). **c**,**d**, LC-MS chromatograms of the reaction mixtures of PaX-Tz **9** (**e**), **10** (**f**), before (i) and after the two stages of Sequence A (ii and iii) and B (iv and v), and the chemical structures of the major products detected. iii for compound **9** was not determined.

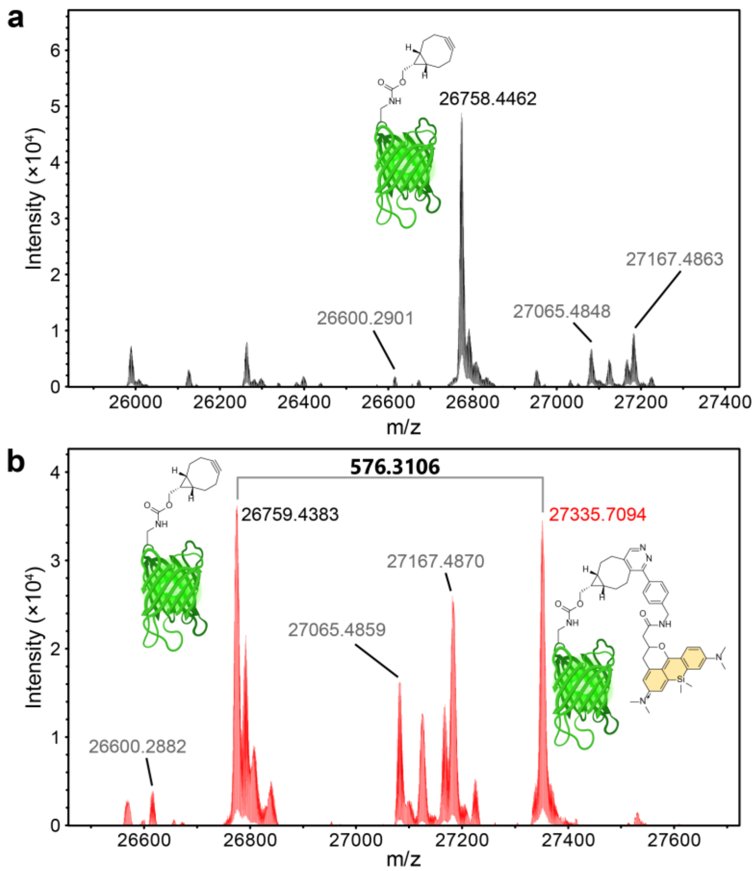

**Figure S10.** Electrospray ionization mass spectrometry of Y35TAG-GFP containing an *endo* BCN-*L*-lysine residue. The spectra of the unlabeled GFP (**a**) and the labeled one (**b**) are presented. Chemical Structure the dye-adduct (**3a**) is shown. Created with BioRender.com.

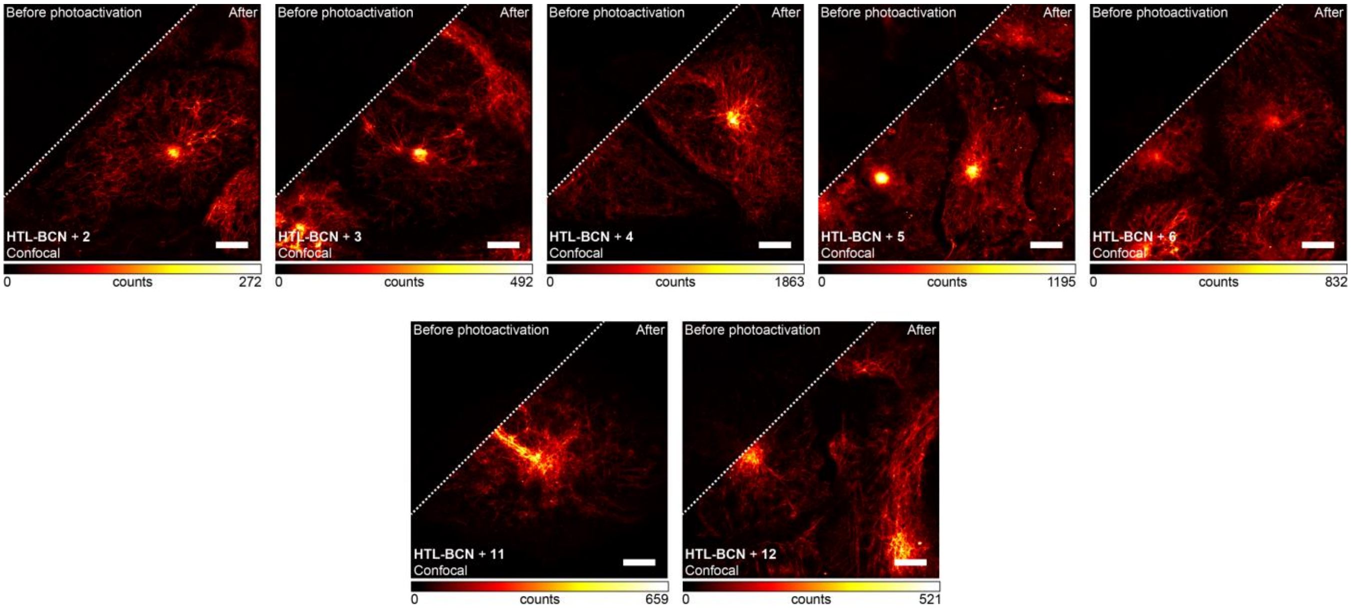

**Figure S11.** Live cell permeability and BCN-specificity were assessed in U2OS cells stably expressing a vimentin-HaloTag construct in a two-step labeling strategy utilizing the HaloTag specific ligand HTL-BCN (10 µM, 30 min) and a dyad (200 nM, overnight), fixed with PFA and imaged by confocal. Conversion to PaXCF was achieved by 405 nm illumination.

**a**

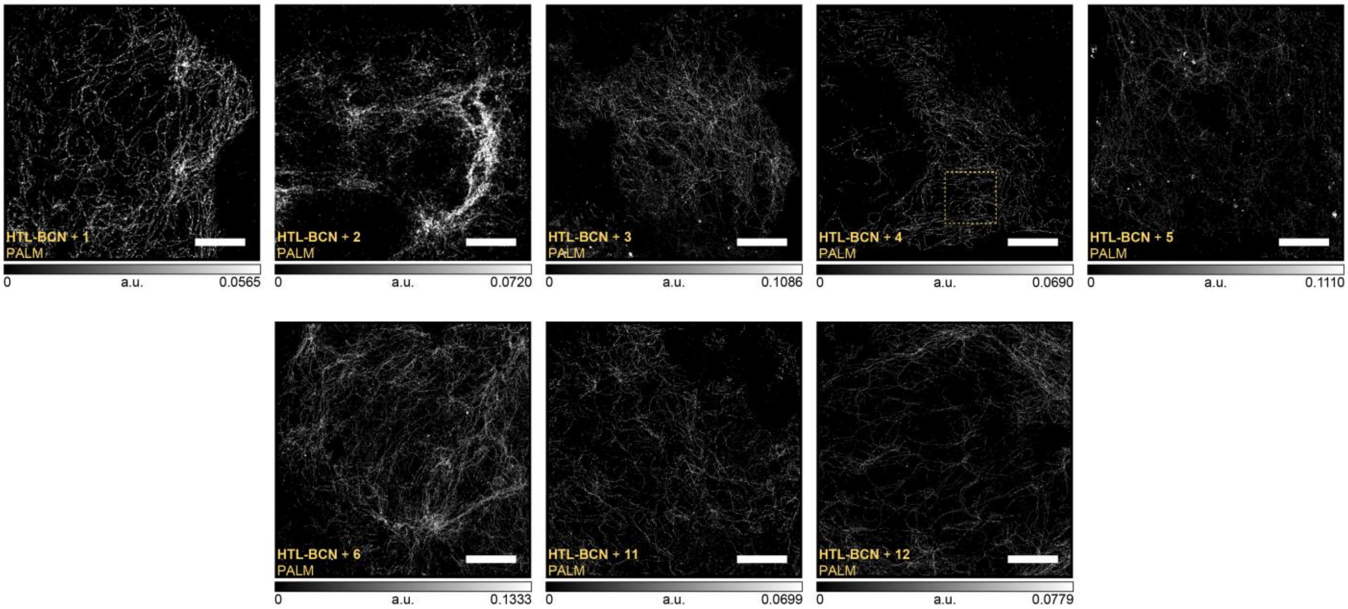

**b**

| | 1 | 2 | 3 | 4 | 5 | 6 | 11 | 12 |
|---|---|---|---|---|---|---|---|---|
| **Mean Photon Count** | $8.2\times10^2$ | $7.2\times10^2$ | $1.8\times10^3$ | $1.5\times10^3$ | $2.6\times10^3$ | $2.1\times10^3$ | $1.6\times10^3$ | $2.2\times10^3$ |
| **Mean Localization Precision (nm)** | 25 | 24 | 14 | 16 | 11 | 13 | 15 | 14 |
| **Median Localization Precision (nm)** | 25 | 24 | 13 | 15 | 10 | 11 | 14 | 12 |

**Figure S12. a**, PALM images of cells stably expressing a vimentin-HaloTag construct, labelled in a two-step strategy utilizing the HaloTag specific ligand HTL-BCN (10 µM, 30 min) and a dyad (200 nM, overnight), fixed with PFA and imaged. The marked region denotes the area displayed in Figure 4c. **b**, Mean photon counts and mean and median localization precisions for the images shown in a. Scale bars: 5 µm.

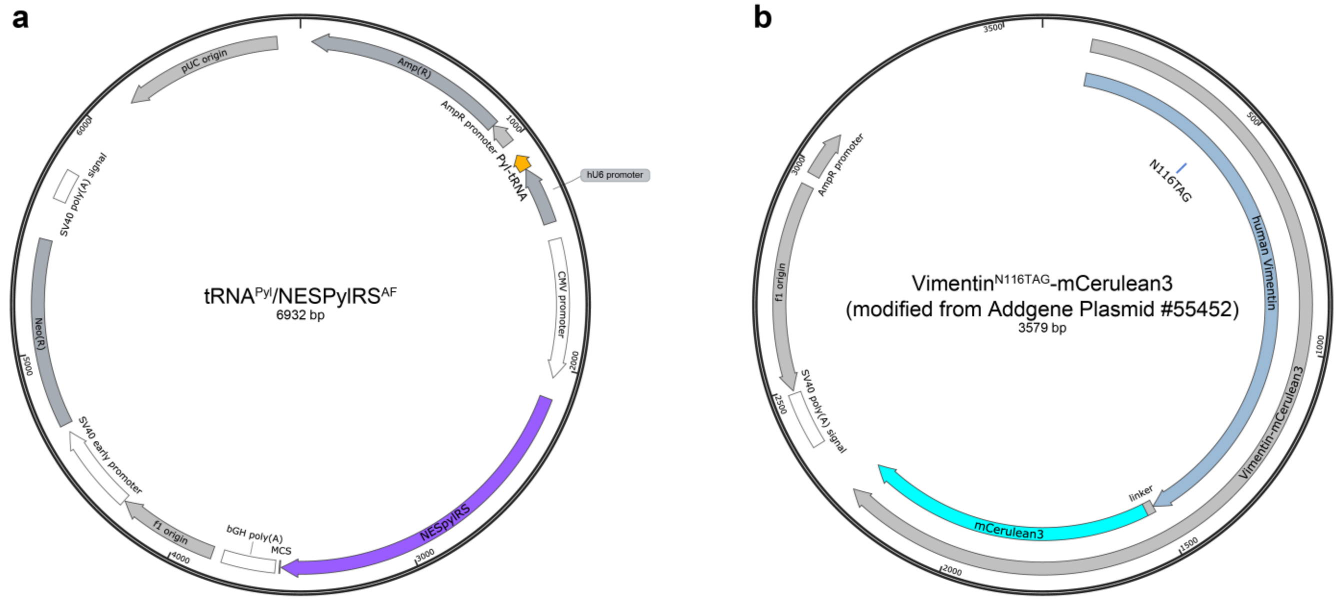

**Figure S13.** Plasmids maps of CMV_NES-PylRS(AF)_hU6tRNAPyl[1] and Vimentin(N116TAG)-mCerulean3 (pVim-Cer) (modified by the Lemke Group from Addgene, #55452[2]) for incorporating unnatural amino acids. Both plasmids were provided by Lemke Lab (EMBL, Heidelberg).

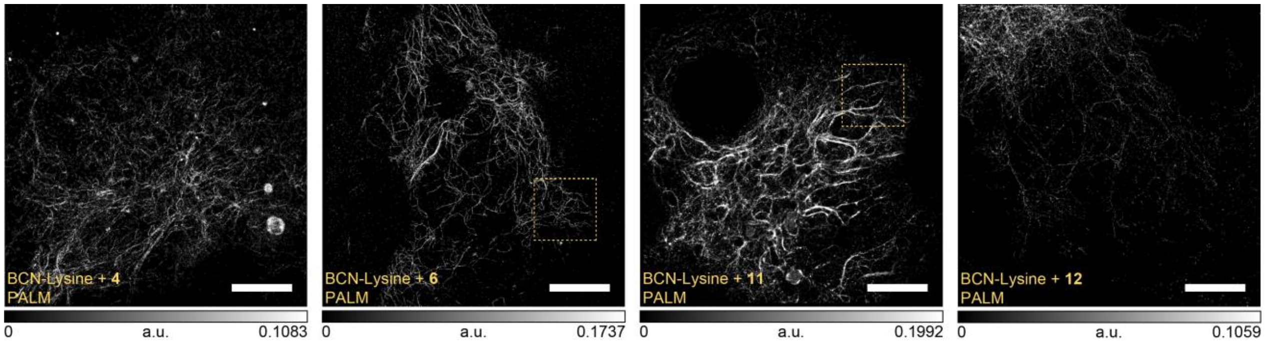

**Figure S14.** PALM images of COS-7 cells transiently expressing a vimentin-mCerulean3 fusion construct[2] carrying a N116TAG mutation (for incorporating the UAA endo BCN-L-lysine) in vimentin;[1, 3] plasmids were provided by the Lemke lab (see Figure S13 for detailed maps of tRNAPyl/NESPylRSAF, and vimentinN116TAG-mCerulean3 plasmids). Transfected cells were labeled with a dyad (500 nM, 4 h), fixed with PFA and imaged. The marked regions denote the areas displayed in Figure 4e. Scale bars: 5 µm.

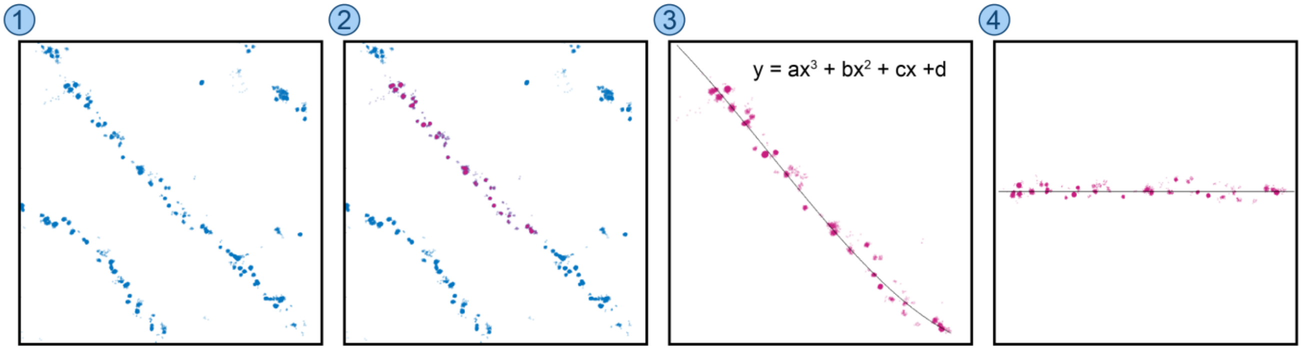

**Figure S15.** Representative diagram of the filament linearization workflow. From a rendered image of MINFLUX localizations (**1**), a filament segment of interest was manually brushed (**2**). The selected localizations were fit to a third-degree polynomial (**3**) to obtain the straightened filament segment (**4**) which was used for subsequent analysis.

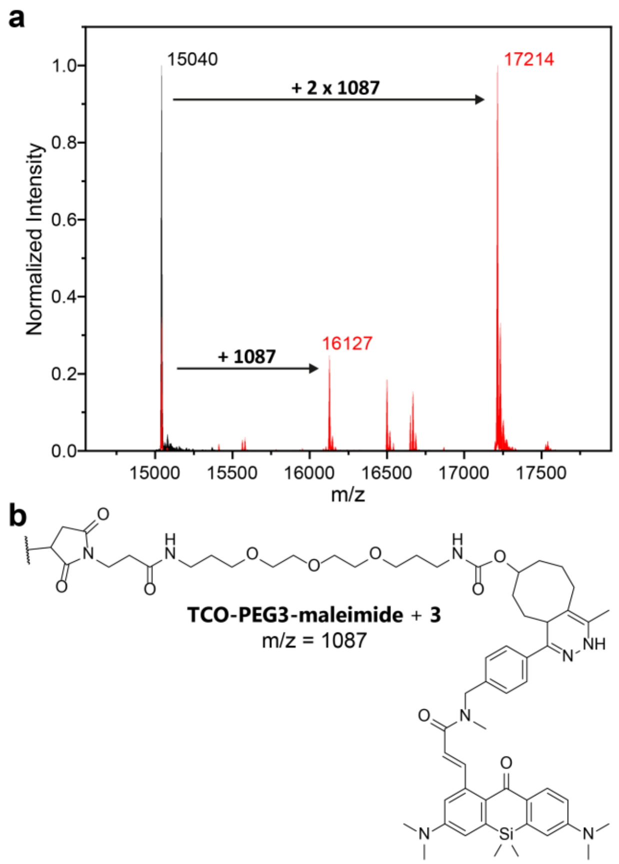

**Figure S16. a**, Electrospray ionization mass spectrometry of FluoTag-X2 anti-GFP nanobody clone 1H1 containing two cysteine residues. The spectra of the unlabeled nanobody (black) and the labeled one (red) are presented. **b**, Chemical Structure and the molecular mass of the dye-adduct (composed of **3** + **TCO-PEG3-Maleimide**).

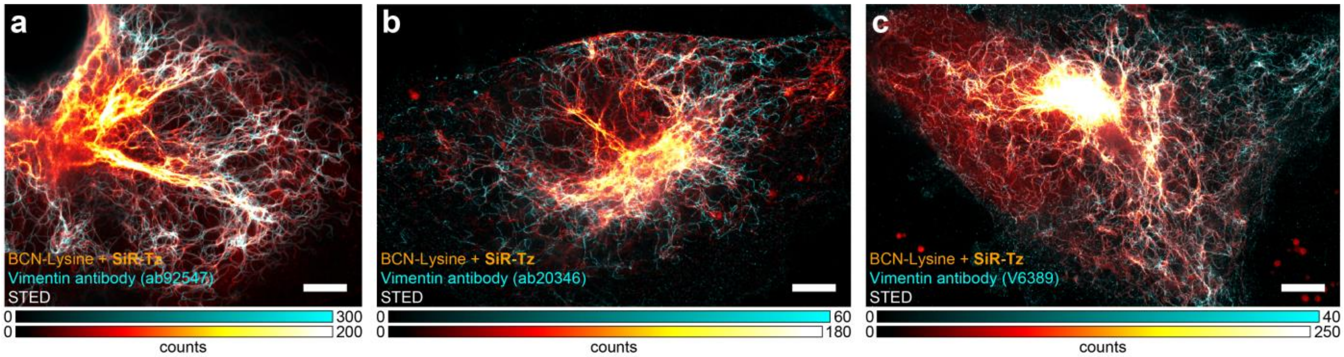

**Figure S17.** STED image of COS-7 cells transiently expressing a vimentin-mCerulean3 fusion construct carrying a N116TAG mutation (for incorporating the UAA endo BCN-L-lysine) in vimentin. Cells were labeled with SiR-tetrazine (1µM, 30 min), fixed with PFA and stained with the indicated anti-vimentin primary antibodies and secondary antibodies labeled with Alexa594. Two color images in SiR-tetrazine (red hot) and Alexa594 (cyan) channels show two separate populations of vimentin filaments. Scale bars: 5 µm.

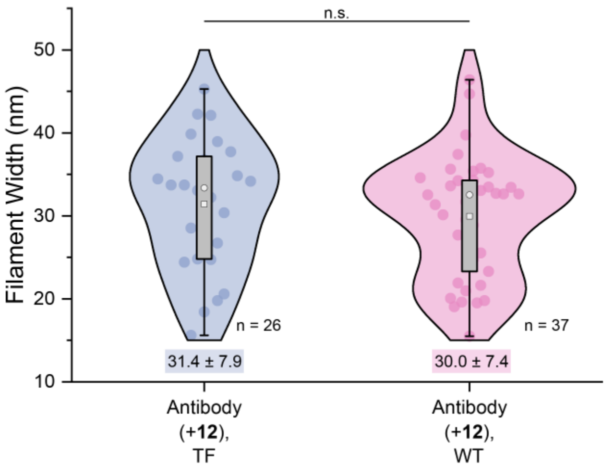

**Figure S18.** Distribution of filament thicknesses of vimentin segments labeled with anti-vimentin primary antibodies and secondary antibodies labeled with **12** in transfected COS-7 cells (TF) transiently expressing a vimentin-mCerulean3 fusion construct carrying a N116TAG mutation (for incorporating the UAA endo BCN-L-lysine) in vimentin), and in wildtype COS-7 cells (WT). n.s. not significant using the Kruskal-Wallis and the Dunn method for post-hoc correction for multiple comparisons.

### SUPPLEMENTARY METHODS

#### General Experimental Information and Synthesis

All chemical reagents (TCI, Sigma-Aldrich, Alfa Aesar) and dry solvents for synthesis (over molecular sieves, AcroSeal package, Acros Organics) were purchased from reputable suppliers and were used as received without further purification. The products were lyophilized from a suitable solvent system using Alpha 2-4 LDplus freeze-dryer (Martin Christ Gefriertrocknungsanlagen GmbH).

#### Thin Layer Chromatography

Normal phase TLC was performed on silica gel 60 $F_{254}$ (Merck Millipore, Germany). For TLC on reversed phase silica gel 60 RP-18 $F_{254}$s (Merck Millipore) was used. Compounds were detected by exposing TLC plates to UV-light (254 or 366 nm) or heating with vanillin stain (6 g vanillin and 1.5 mL conc. $H_2SO_4$ in 100 mL ethanol), unless indicated otherwise.

#### Flash Chromatography

Preparative flash chromatography was performed with an automated Isolera One system with Spektra package (Biotage AG) using commercially available cartridges of suitable size as indicated (RediSep Rf series from Teledyne ISCO, Puriflash Silica HP 30µm series from Interchim).

#### Nuclear Magnetic Resonance (NMR)

NMR spectra were recorded on a Bruker DPX 400 spectrometer utilizing the Bruker Topspin 3.5 software. All spectra are referenced to tetramethylsilane as an internal standard (δ = 0.00 ppm). Multiplicities of the signals are described as follows: s = singlet, d = doublet, t = triplet, q = quartet, m = multiplet or overlap of non-equivalent resonances; br = broad signal. Coupling constants $^{n}J_{X\text{-}Y}$ are given in Hz, where n is the number of bonds between the coupled nuclei X and Y ($J_{H\text{-}H}$ are always listed as $J$ without indices).

#### Mass-Spectrometry (MS)

Low resolution mass spectra (100 - 1500 *m/z*) with electro-spray ionization (ESI) were obtained on a Shimadzu LC-MS system described below. High resolution mass spectra (HRMS) were obtained on a maXis II ETD (Bruker) with electrospray ionization (ESI) at the Mass Spectrometry Core facility of the Max-Planck Institute for Medical Research (Heidelberg, Germany).

#### High-Performance Liquid Chromatography (HPLC)

Analytical liquid chromatography-mass spectrometry was performed on an LC-MS system (Shimadzu, controlled with LabSolutions 5.89 software): 2x LC-20AD HPLC pumps with DGU-20A3R solvent degassing unit, SIL-20ACHT autosampler, CTO-20AC column oven, SPD-M30A diode array detector and CBM-20A communication bus module, integrated with CAMAG TLC-MS interface 2 and LCMS-2020 spectrometer with electrospray ionization (ESI, 100 – 1500 *m/z*). Analytical column: Hypersil GOLD 50×2.1 mm 1.9µm, standard conditions: sample volume 1-2 µL, solvent flow rate 0.5 mL/min, column temperature 30 °C. General method: isocratic 95:5 A:B over 2 min, then gradient 95:5 – 0:100 A:B over 5 min, then isocratic 0:100 A:B over 2 min; solvent A = water + 0.1% v/v $HCO_2H$, solvent B = acetonitrile + 0.1% v/v $HCO_2H$.

Preparative high-performance liquid chromatography was performed on a Büchi Reveleris Prep system using the suitable preparative columns and conditions as indicated for individual preparations. Method scouting was performed on a HPLC system (Shimadzu, controlled with LabSolutions 5.89 software): 2x LC-20AD HPLC pumps with DGU-20A3R solvent degassing unit, CTO-20AC column oven equipped with a manual injector with a 20 µL sample loop, SPD-M20A diode array detector, RF-20A fluorescence detector and CBM-20A communication bus module; or on a Dionex Ultimate 3000 UPLC system: LPG-3400SD pump, WPS-3000SL autosampler, TCC-3000SD column compartment with 2× 7-port 6-position valves and DAD-3000RS diode array detector. The test runs were performed on analytical columns with matching phases (HPLC: Interchim 250×4.6 mm 10 µm C18HQ, Interchim 250×4.6 mm 5 µm PhC4, solvent flow rate 1.2 mL/min; UPLC: Interchim C18HQ or PhC4 75×2.1 mm 2.2 µm, ThermoFisher Hypersil GOLD 100×2.1 mm 1.9 µm, solvent flow rate 0.5 mL/min).

#### Optical spectroscopy

Fluorescence lifetimes were measured with a FluoTime 300 fluorescence lifetime spectrometer (PicoQuant, controlled with the EasyTau1.4 software). All measurements were performed in air-saturated solvents at ambient temperature.

#### Antibodies, Nanobodies and Other Fluorescent Conjugates

anti-vimentin EPR3776 rabbit, Abcam, ab92547; unconjugated AffiniPure goat antirabbit, Jackson ImmunoResearch, 111-005-003; unconjugated FluoTag-X2 anti-GFP clone 1H1, NanoTag Biotechnologies, N0302; FluoTag-X2 anti-GFP STAR635P, NanoTag

Biotechnologies, N0304-Ab635P-S; SiR-tetrazine, Spirochrome, SC008; Anti-Vimentin antibody [EPR3776] - Cytoskeleton Marker, abcam, ab92547; Anti-Vimentin antibody [VI-10], abcam, ab20346; Anti-Vimentin antibody clone V9, Sigma Aldrich, clone V9; Goat anti-Mouse IgG (H+L) Alexa Fluor™ 594, Invitrogen, A-11032; Donkey anti-Rabbit IgG (H+L) Alexa Fluor™ 594, Invitrogen, A-21207.

Antibodies and nanobodies were used without further validation as the obtained labelling was clearly compatible with the expected structures.

**Photolysis and model click reactions of PaX-Tz compounds**

Solutions in methanol (1.66 µg $mL^{-1}$) were irradiated in a previously described[4] home-built setup with a 405 nm LED source (M405L3, Thorlabs Inc.) in combination with a bandpass (10 nm) filter (FB405-10, Thorlabs Inc.). During the irradiation, samples were maintained at 20 °C and continuously stirred with a Peltier-based temperature-controlled cuvette holder (Luma 40, Quantum Northwest, Inc.). The absorption and emission of irradiated solutions was monitored at desired irradiation intervals with a fiber-based spectrometer (Flame-S-UV-Vis-ES, Ocean Insight). For absorption measurements, a deuterium and tungsten halogen source was used for illumination (DH-2000-BAL, Ocean Insight), and for fluorescence excitation was performed in a 90° configuration with a 505 nm LED source (M505L3, Thorlabs Inc.) in combination with a bandpass (10 nm) filter (FL508,5-10, Thorlabs). Click reaction with model compound BCN-OH ((1R,8S,9s)-Bicyclo[6.1.0]non-4-yn-9-ylmethanol, Sigma Aldrich) were studied in the same setup, with dark reactions intervals (i.e. no irradiation performed) between the recording of the absorption and emission. The reaction was started after the addition of 50 eq. (H-Tz's) or 100 eq. (Me-Tz's) of BCN-OH. Data collection and analysis was performed with custom-made routines in MATLAB. Samples for LCMS or ESI-MS analysis were taken before and after photolysis was performed.

To prepare the adduct of compound **3a** and Y35TAG-GFP, compound **3** was irradiated in the setup described above until full conversion to **3a**, and mixed with 1.1 equivalents of the protein in phosphate buffer (100 mM, pH = 7.0). The evolution of the emission of the FRET donor (GFP) and the acceptor (**3a**) was recorded in a Varian Cary Eclipse fluorescence spectrophotometer (Agilent Technologies controlled with the Cary Eclipse Scan Application 1.2(147)).). At the end of the experiment, the sample was submitted for mass analysis (ESI) along with an aliquot of unreacted protein. The modified GFP plasmid was purchased from Addgene (#92225) and the pEvolv_tRNAPyl/PylRSAF plasmid for bacterial expression of GFP mutant was provided by the Lemke lab (EMBL, Heidelberg).

**Confocal and STED (stimulated emission depletion) microscopy.**

Confocal and STED images were acquired using two Abberior Expert Line (Abberior Instruments GmbH, Göttingen, Germany) fluorescence microscopes built on a motorized inverted microscope IX83 (Olympus, Tokyo, Japan). Microscope 1 is equipped with pulsed STED lasers at 595 nm and 775 nm shaped by Spatial Light Modulators (SLMs), and with 355 nm, 405 nm, 485 nm, 561 nm, and 640 nm excitation lasers, and a 100x/1.40 oil immersion objective lenses (Olympus). Microscope 2 is equipped with pulsed STED lasers at 655 nm and 775 nm, and with 520 nm, 561 nm, 640 nm, and multiphoton (Chameleon Vision II, Coherent, Santa Clara, USA) excitation lasers, and a 60x/1.42 oil immersion objective lens (Olympus). Imaging and image processing were done with Imspector software, and all images are displayed as raw data unless otherwise noted.

Confocal images were acquired on Microscope 1. Confocal imaging was performed with a pixel size of 80×80 nm, and pixel by pixel activation/excitation. Imaging of $PaX_{560}$ was performed with a 561 nm excitation laser (13–15 µW, 60–150 µs), and a detection window of 571–691 nm, and a 405 nm activation laser (<13 µW, 20–50 µs). Confocal imaging of $PaX_{480}$ was performed with a 485 nm excitation laser (13 µW, 90 µs), a detection window of 515–600 nm, and a 405 nm activation laser (50 µW, 20–50 µs). Confocal imaging of mCerulean3 was performed with a detection window of 440–570 nm and a 405 nm excitation laser (<13 µW, 20 µs), which also acted as the activation for the $PaX_{560}$ channel.

STED images of samples labeled with Alexa594 and SiR-tetrazine were acquired on Microscope 1. Imaging was performed with 40×40 nm pixel size and 2 line accumulations. Imaging of Alexa594 was performed with a 561 nm excitation laser (5.5 µW, 10 µs), followed by the 775 nm STED laser (750 ps delay, 8 ns width, 150 mW), a detection window of 584–630 nm and 3 line accumulations. Imaging of SiR was performed with a 640 nm excitation laser (10 µW, 10 µs), followed by the 775 nm STED laser (750 ps delay, 8 ns width, 75 mW), a detection window of 650–763 nm and 3 line accumulations. STED images of samples labeled with $PaX_{560}$ compounds were acquired on Microscope 2 due to the availability of the 660 nm STED line.

Imaging was performed with 25×25 nm pixel size, 4 line accumulations, a 561 nm excitation laser (2 µW, 60 µs), followed by the 660 nm STED laser (750 ps delay, 8 ns width, 3 µW), and a detection window of 580–800 nm. The activation was performed via a fluorescence lamp due to the absence of a 405 nm activation laser. Corresponding confocal image was acquired with 25×25 nm pixel size, no line accumulations, a 561 nm excitation laser (1 µW, 60 µs), and a detection window of 580–800 nm.

**Superresolution single molecule localization microscopy (SMLM) / Photoactivated localization microscopy (PALM)**

Images were acquired on a custom-built setup[5], equipped with a 473 nm (500 mW), a 532 nm (1 W) and a 560 nm (1 W) laser for excitation, a 405 nm (300 mW) laser for activation, a back illuminated EMCCD camera (Andor iXon 897 / 512×512 sensor), and a Leica HCX PL APO CS 100x/1.46 oil lens. Emission light was separated from the excitation and activation light with proper combination of a dichroic mirror and a suppression filter for each excitation laser in filter cubes green (505 nm, Chroma T 505 lpxr / 508–598 nm, Semrock 550/88 BL HC) and orange (580 nm, Semrock HC R561 / 589–739 nm, Semrock 665/150 BL HC). A movable mirror was used to switch between wide field, highly inclined and laminated optical sheet (HILO) and total internal reflection fluorescence (TIRF) illumination modes. Images were acquired with the orange (for $PaX_{560}$ derivatives) and green filter cubes (for $PaX_{480}$ derivatives), 20 ms exposure time, and 20–30% of maximum available powers of 560 (for $PaX_{560}$ derivatives) and 473 nm (for $PaX_{480}$ derivatives) excitation lasers, depending on the dye and sample properties. The 405 nm activation laser was incorporated as 200 µs pulses, in between frames, gradually increased up to the maximum available power a power up to 1 mW in the back focal plane. The measurement was stopped when the events became sparse. The microscope components were controlled with custom LabView (2019 32bit) software, and the camera was controlled with the Andor Solis 4.31.30022 software package.

All images were analyzed and processed using the ThunderSTORM plugin[2] on ImageJ (version 1.52p). In brief, images were filtered with a wavelet filer (B-spline order 3, scale 2.0), approximate localization of the molecules was performed with a local maximum method (peak intensity threshold of ca. 1.4–1.6 standard deviations; connectivity 8-neighbourhood), and sub-pixel localization of the molecules was performed with a maximum likelihood fitting method (PSF integrated method, fitting radius of 3 pixels, initial sigma 1.6). Post-processing was performed with the same plug-in. Data was drift-corrected based on the cross-correlation method, merged within the size of 0.5 pixel with 0 off-frames allowed. Sigma values were filtered to converge to a normal Gaussian distribution function. Photon numbers, number of detections per molecule and uncertainty values were restricted (to 200 < intensity < 20000, detections < 20 and 4 < uncertainty < 40) to filter any outliers. Lastly, a density filter was applied to remove noise caused by isolated localizations without at least 3 neighboring localizations in 100 nm proximity. Final images were produced using normalized Gaussian rendered with variable sigma localization uncertainties, and a pixel-size of 5 nm.

**MINFLUX microscopy**

MINFLUX microscopy was performed on an Abberior Instruments 3D MINFLUX microscope equipped with 560 nm and 640 nm (cw) excitation laser lines for MINFLUX, a 488 nm (cw) excitation laser for confocal imaging, and a 405 nm (cw) activation laser, SLM-based beam shaping modules, and an EOD-based MINFLUX scanner[6]. Fluorescence photons emitted from the sample were counted using an avalanche photo diode with filter-based detection in the Cy3 channel (580–630 nm). To ensure measurements with molecular precision, the Abberior MINFLUX microscope is equipped with a 975 nm illumination laser for stability control. The position of the sample was locked on the gold beads. Imaging was achieved by using 22–26 µW (before the scanner) of 561 nm laser in the first iteration. The activation 405 nm laser was attenuated with an ND2 filter. Activation was switched on and the power was gradually increased up to ~0.5 µW, sustaining the frequency of detected events, until the events became sparse in time and the imaging was stopped. Imaging was achieved using 30 photons per localization and a final TCP diameter L of 40 nm. Images were processed in the microscopy software Imspector 16.3 to aggregate the localizations to 100 photons per localization prior to analysis.

Images were post-processed and analyzed with a custom-built MATLAB (2022a, MathWorks) routine. Prior to analysis, the data was drift corrected using the redundant cross-correlation algorithm. In a short, Drift correction was performed by dividing the localizations into time windows, rendering the images for each window and calculating the 3D correlation function between images from different time points. To obtain the estimated 3D drift path, the cross correlation between time windows was used. The final drift trajectory was then subtracted from the localization coordinates. Only trace IDs (TIDs) containing more than 3 localizations were considered. A bivariate normal distribution was fitted to the localizations belonging to the same trace using fitgmdist, a MATLAB built-in function with default settings. Mahalanobis distance was used to compute the distance of each localization to the fitted distribution, using MATLAB's mahal function. Localizations farther than the third quartile of the Mahalanobis distances from all traces were defined as outliers and therefore excluded from its respective trace. The filament's regions of interest (ROIs) were manually selected by brushing directly from the localizations, with a minimum ROI length of 100 nm. Filaments were thereafter linearized to a third order polynomial fit and the Root Mean Square Error was used as metric for the goodness of fit. Filament lengths were calculated as the length of the fitted curve. Filament width was estimated as the full-width half maximum (FWHM) of the Gaussian fit of the localization projection into the filament secondary axis. Statistical analysis of FWHM of different labeling strategies was performed in Origin 2020b using the Kruskal-Wallis ANOVA method to assess significance. The distances between localizations clusters along the filament trace axis were calculated from the histogram peaks with 4 nanometers bin width.

**Two-step Click Conjugation of Antibodies**

Bioconjugation of antibodies was performed according to a procedure adapted from the literature.[7] The pH of the secondary unconjugated AffiniPure goat anti-rabbit solution (2.4 mg/l, Jackson ImmunoResearch, 111-005-003) was adjusted to pH≈8 by addition of aqueous NaHCO3 (1 M) to a final concentration of 100 mM, and 20 equivalents of endo-bicyclo[6.1.0]non-4-in-9-ylmethyl]-N-succinimidylester (SiChem GmbH, SC-8074) in DMF (34 mM) was added. After incubation for 1 h at room temperature the antibody was purified by size exclusion chromatography with a 7K MWCO Zeba Spin Desalting Column (Thermo Scientific). Then, 6 equivalents of **3**, **11** or **12** in DMF (2 mM) was added and incubated for 1.5 h at room temperature. The labelled antibody was purified with a desalting column (7K MWCO Zeba Spin Desalting Columns, Thermo Scientific).

**Cell culture**

COS-7 (Cell Lines Service, 665470) cells were cultivated in DMEM:Ham's F12 medium (Gibco, 11320074) supplemented with 5% (v/v) fetal bovine serum (ThermoFisher, 10500064) and 1% penicillin-streptomycin (Gibco, 15140122) and U2OS-Vim-Halo (AG Stefan Jakobs, Göttingen) cells were cultivated in Dulbecco's modified Eagle medium (DMEM) (Gibco, 31966021) supplemented with 10% (v/v) fetal bovine serum and 1% penicillin-streptomycin in an incubator (37°C, 5% $CO_2$, 95% relative humidity). For dissociation of cells, TrypLE Express (Gibco, 12604013) was used. Cells were split every 4 days or at 80–90% confluency and regularly tested for mycoplasma contamination. For live imaging experiments, FluoroBrite DMEM (Gibco, A1896701) supplemented with 10% (v/v) fetal bovine serum, 1% GlutaMAX and 1% penicillin-streptomycin was used as the mounting medium.

##### Immunostaining with HaloTag Ligand-Bicyclononyne Linker

The staining was performed according to a procedure adapted from the literature.[8] Stock solutions of the HaloTag ligand-bicyclononyne linker (HTL-BCN) were prepared in DMSO (10 mM) and stored at -20°C for maximum 3 months until use. U2OS cells expressing HaloTag on vimentin were incubated in FluoroBrite DMEM (serum-free without additives) containing HTL-BCN (10 µM) for 30 min at 37 °C. The cells were washed (3×30 min) with the serum-free medium. Next, the cells were incubated overnight (37°C) in FluoroBrite DMEM supplemented with 10% (v/v) fetal bovine serum, 1% GlutaMAX and 1% penicillin-streptomycin containing the indicated PaX-Tz derivative (200 nM). Coverslips were washed (3×10 min) with the same medium and the fixation was performed with formaldehyde as described below.

##### Transient Transfection for Genetic Code Expansion for Incorporation of Bicyclononyne-*L*-Lysine in Vimentin

Plasmids —Vimentin(N116TAG)-mCerulean3 (modified by Lemke Group from Addgene, #55452 [2]) and CMV_NES-PylRS(AF)_hU6tRNAPyl[9]— for incorporating unnatural amino acids were provided by Lemke Lab (EMBL, Heidelberg) with plasmid maps shown in Figure S13. Transfection was performed according to a procedure adapted from the literature.[3] A Countess II FL Automated Cell Counter (Thermo Fisher Scientific) and Trypan Blue staining (0.4%) was used to seed preferred cell count prior to transfection. COS-7 cells ($0.5*10^5$–$1*10^5$ cells) were seeded on glass coverslips in 12-well plates in 1 ml of the corresponding cell culture medium per coverslip. After 16–24 h, half of the cell culture medium was exchanged with fresh medium containing *endo*-bicyclo[6.1.0]nonyne-*L*-lysine (BCN-*L*-lysine) (SiChem GmbH, SC-8014) to yield a final concentration of 200 µM and placed in the cell incubator for 30 min. Per well, 2 µl jetPRIME transfection reagent (Polyplus-transfection, 101000046) was mixed with 1 µg of each plasmid in 75 µl jetPRIME buffer, and incubated at room temperature for 15min. The mixture was added to the cell medium dropwise (75 µl per well), mixed and incubated overnight at 37°C.

Cells were washed (3×2 h) with fresh cell culture medium to remove excess BCN-*L*-lysine and the transfection efficiency was checked on a cell imaging multimode reader (Cytation 5, Biotek) prior to staining and imaging. Next, the cells were incubated for 4 h with the indicated PaX tetrazine derivative (500 nM) in FluoroBrite DMEM supplemented with 10% (v/v) fetal bovine serum, 1% GlutaMAX and 1% penicillin-streptomycin. Coverslips were washed (3×10 min) with the same medium and the fixation was performed with formaldehyde as described below. A counter staining with FluoTag-X2 anti-GFP STAR635P was performed in order to identify transfected cells and select regions of interests for imaging.

**Fixation with Paraformaldehyde**

After transfection and/or click labeling, cells were washed twice with PBS, and then treated with a 4% formaldehyde solution in PBS (PFA) at room temperature for 25 min. Fixed cells were washed twice with PBS, and treated with a quenching solution (0.1 M NH4Cl and 0.1 M Glycine in PBS) for 7 min at room temperature. Coverslips were incubated in blocking buffer (2% BSA + 0.1% Triton X-100 (AppliChem GmbH) in PBS) for 15–30 min at room temperature. Fixed cells were further handled as described in Immunostaining.

**Immunostaining with Affinity Probes**

The coverslips were overlaid with the primary antibody or anti-GFP nanobody solution prepared in a 1:1 dilution of blocking buffer in PBS and incubated in a humid chamber for 60–90 min at room temperature. Next, the coverslips were washed with PBS (3×5 min). If indicated, coverslips were overlaid with the secondary antibody solution prepared in a 1:1 dilution of (blocking buffer in PBS, and incubated in a humid chamber for 60–90 min at room temperature. For samples labeled with primary/secondary antibodies or anti-GFP nanobody, a counter staining with SiR-tetrazine was performed in order to identify transfected cells and select regions of interests for imaging. Finally, coverslips were washed with PBS (3×5 min). Prior to imaging on the MINFLUX setup, samples were treated with 150 nm gold beads (BBI Solutions, EM.GC150) for camera-based stabilization. Samples were mounted in Mowiol or PBS (for longer storage) and stored at 4°C until use.

**Computational Methods**

All quantum mechanical calculations were performed using Gaussian 16 Rev. B. 01.[10] All DFT and TD-DFT calculations were conducted with the ωB97X-D[11] functional in combination with the

Def2-SVP basis set[12] with an applied SMD model solvation of water.[13-14] Molecular structures and orbitals were visualized using GaussView 6. Frequency checks were carried out after each geometry optimization to ensure that the minima on the potential energy surfaces were found. Franck-Condon calculations were performed using the *External Iteration approach* for solvation.

### SYNTHESIS AND CHARACTERIZATION

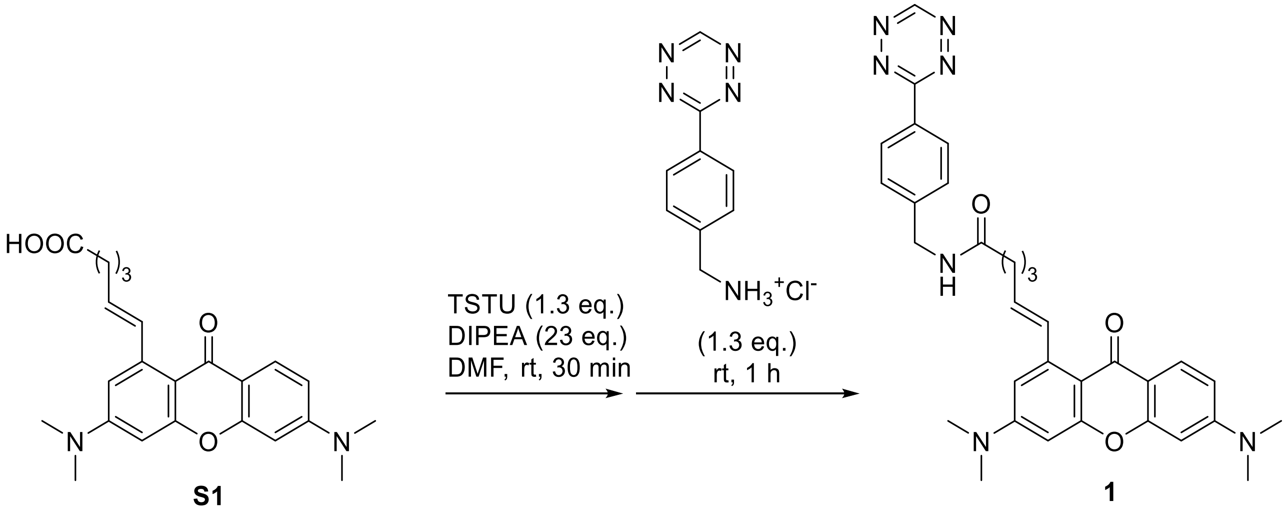

**1**

PaX$_{480}$-Tz

**Compound 1.** In an amber vial, compound **S1** (8.0 mg, 0.020 mmol) and TSTU (7.8 mg, 0.026 mmol) were dissolved in DMF (200 µL). DIPEA (59 mg, 0.45 mmol) was added, and the mixture was stirred for 30 min at rt. 3-(*p*-Benzylamino)-1,2,4,5-tetrazine hydrochloride (5.6 mg, 0.025 mmol) was added, and the reaction was stirred at rt for 1 h. The volatiles were removed *in vacuo*. The product was isolated isolated by preparative HPLC (Interchim Uptisphere Strategy PhC4 250×21.2 mm 5 µm, solvent flow rate 18 mL/min, gradient 50% to 80% A:B, A – acetonitrile + 0.1% (v/v) trifluoroacetic acid, B – water + 0.1% (v/v) trifluoroacetic acid) and freeze-dried from dioxane to yield 6.0 mg (52%) of **1** as a red solid.

$^{1}$H NMR (400 MHz, Chloroform-*d*) δ 10.18 (s, 1H), 9.05 (s, 1H), 8.44 (d, *J* = 8.4 Hz, 2H), 7.66 (d, *J* = 9.0 Hz, 1H), 7.50 (d, *J* = 8.3 Hz, 2H), 7.41 (d, *J* = 15.5 Hz, 1H), 6.55 – 6.47 (m, 2H), 6.39 (d, *J* = 2.6 Hz, 1H), 6.33 (d, *J* = 2.4 Hz, 1H), 5.78 (dt, *J* = 15.0, 7.2 Hz, 1H), 4.59 (d, *J* = 5.2 Hz, 2H), 3.09 (s, 6H), 3.02 (s, 5H), 2.57 (t, *J* = 6.3 Hz, 2H), 2.37 – 2.25 (m, 2H), 1.98 (s, 2H).

$^{13}$C NMR (101 MHz, Chloroform-*d*) δ 176.6, 176.2, 166.5, 159.3, 157.8, 157.5, 154.3, 153.5, 144.0, 142.3, 135.0, 130.4, 130.2, 129.2, 128.5, 127.6, 112.4, 109.3, 109.2, 108.9, 96.9, 96.6, 44.1, 40.3, 40.2, 34.2, 31.8, 24.4.

HRMS (ESI) m/z: [M+H]$^{+}$ Calcd for $C_{32}H_{33}N_{7}O_{3}$: 564.2718, found: 564.2719.

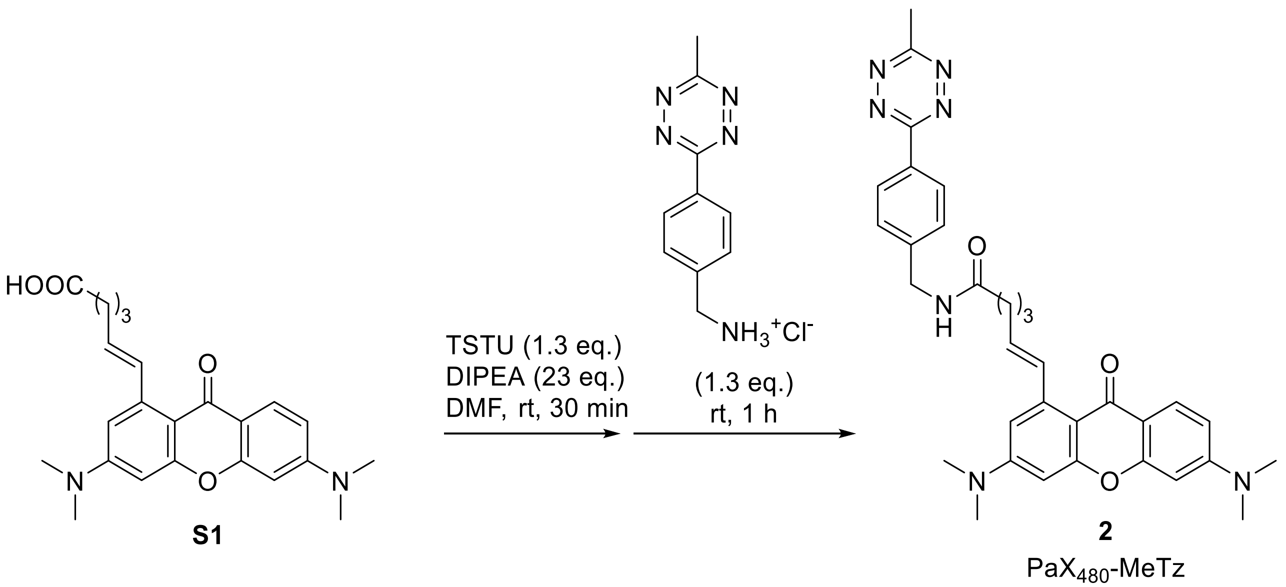

**Compound 2.** In an amber vial, compound **S1** (6.4 mg, 0.016 mmol) and TSTU (6.3 mg, 0.030 mmol) were dissolved in DMF (200 µL) and DIPEA (59 mg, 0.45 mmol) was added and stirred for 1 h. at rt. (4-(6-Methyl-1,2,4,5-tetrazin-3-yl)phenyl)methanamine hydrochloride (5.2 mg, 0.022 mmol) was added and the reaction was stirred at rt for 1 h. The volatiles were removed *in vacuo*. The product was isolated by flash chromatography on a Biotage Isolera system (12g Interchim SiHP 30 µm cartridge, gradient 0% to 100% acetonitrile/dichloromethane) and freeze-dried from dioxane to yield 8.4 mg (88%) of **4** as an orange solid.

$^{1}$H NMR (400 MHz, Chloroform-*d*) δ 8.50 (s, 1H), 8.43 (d, *J* = 8.4 Hz, 2H), 7.71 (d, *J* = 9.0 Hz, 1H), 7.53 (d, *J* = 8.5 Hz, 2H), 7.47 (d, *J* = 15.6 Hz, 1H), 6.56 (d, *J* = 2.6 Hz, 1H), 6.53 (dd, *J* = 9.0, 2.4 Hz, 1H), 6.43 (d, *J* = 2.6 Hz, 1H), 6.40 (d, *J* = 2.4 Hz, 1H), 5.80 (dt, *J* = 15.5, 7.2 Hz, 1H), 4.59 (d, *J* = 4.3 Hz, 2H), 3.09 (s, 7H), 3.07 (s, 3H), 3.02 (s, 6H), 2.53 – 2.44 (m, 2H), 2.42 – 2.26 (m, 3H), 2.03 – 1.91 (m, 2H).

$^{13}$C NMR (101 MHz, Chloroform-*d*) δ 176.3, 174.4, 167.1, 164.2, 159.2, 157.3, 154.0, 153.2, 144.5, 142.5, 134.7, 130.6, 130.3, 129.0, 128.1, 127.8, 112.9, 109.6, 109.4, 108.9, 97.2, 97.1, 43.6, 40.5, 40.4, 34.6, 31.8, 24.6, 21.3.

HRMS (ESI) m/z: $[M+H]^+$ Calcd for $C_{33}H_{35}N_7O_3$: 578.2874 , found: 578.2866.

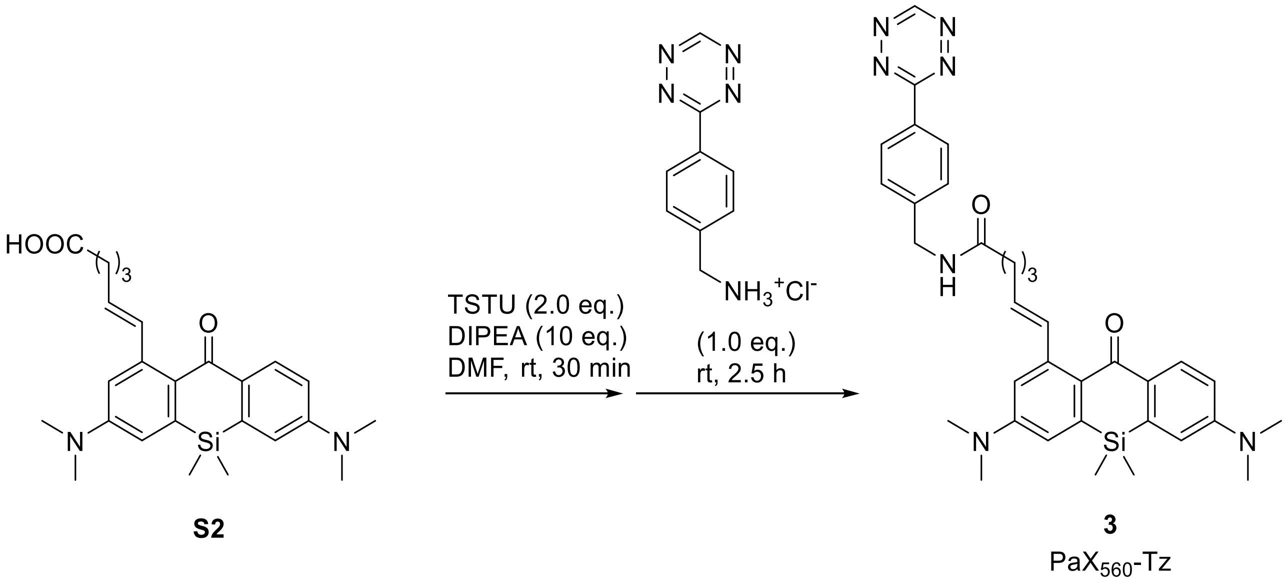

**Compound 3.** In an amber vial, compound **S2** (13.1 mg, 0.030 mmol) and TSTU (11.8 mg, 0.039 mmol) were dissolved in DMF (150 µL). DIPEA (39 mg, 0.3 mmol) was added, and the mixture was stirred for 30 min at rt. 3-(*p*-Benzylamino)-1,2,4,5-tetrazine hydrochloride (7.2 mg, 0.032 mmol) was added, and the reaction was stirred at rt for 2.5 h. The volatiles were removed *in vacuo*. The product was isolated by flash chromatography on a Biotage Isolera system (12g Interchim SiHP 30 µm cartridge, gradient 50% to 90% ethyl acetate/hexane) and freeze-dried from dioxane to yield 5.5 mg (30%) of **3** as an orange solid.

$^{1}$H NMR (400 MHz, DMSO-$d_6$): δ 10.57 (s, 1H), 8.52 (t, $J$ = 6.0 Hz, 1H), 8.45 (d, $J$ = 8.4 Hz, 2H), 7.93 (d, $J$ = 8.9 Hz, 1H), 7.56 (d, $J$ = 8.4 Hz, 2H), 7.15 (d, $J$ = 15.6 Hz, 1H), 6.84 (t, $J$ = 3.0 Hz, 2H), 6.79 (dd, $J$ = 9.0, 2.8 Hz, 1H), 6.73 (d, $J$ = 2.8 Hz, 1H), 5.88 (dt, $J$ = 15.5, 6.9 Hz, 1H), 4.45 (d, $J$ = 5.8 Hz, 2H), 3.30 (s, 6H), 3.05 (s, 6H), 3.02 (s, 7H), 2.32 (t, $J$ = 7.2 Hz, 2H), 2.22 (q, $J$ = 6.8 Hz, 2H), 1.78 (p, $J$ = 7.3 Hz, 2H), 0.42 (s, 6H).

$^{13}$C NMR (101 MHz, DMSO-$d_6$): δ 186.7, 172.4, 165.4, 158.1, 151.0, 150.6, 145.1, 142.6, 140.9, 138.5, 134.3, 131.0, 130.6, 130.3, 128.6, 128.1, 127.8, 126.9, 114.5, 113.9, 113.0, 112.9, 41.9, 39.3, 39.4, 34.7, 32.0, 25.1, -1.1.

HRMS (ESI) m/z: $[M+H]^+$ Calcd for $C_{34}H_{39}N_7O_2Si$: 606.3007, found: 606.3007.

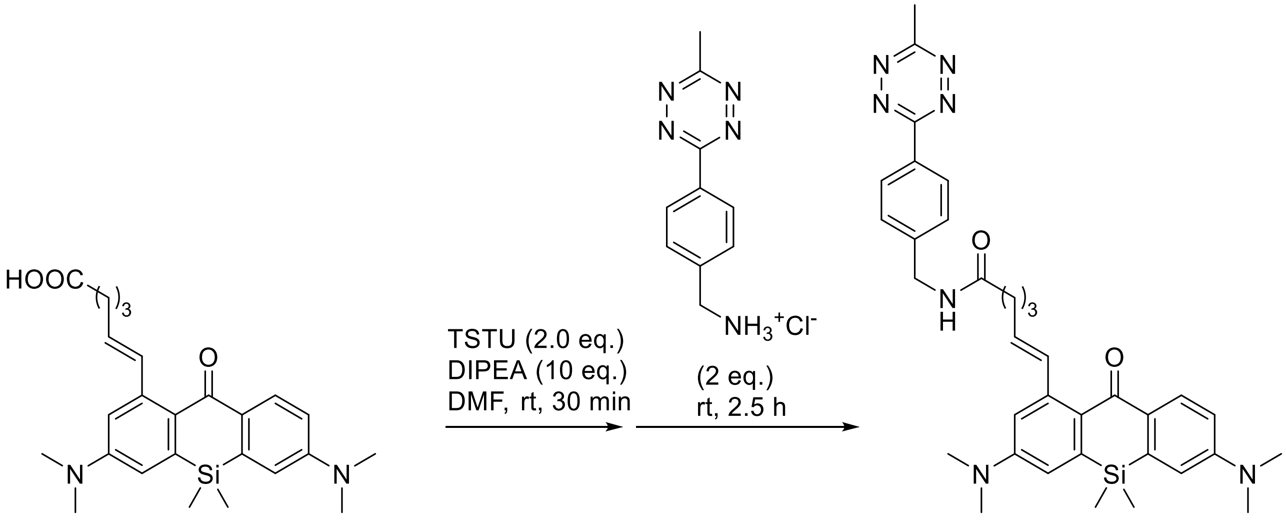

**S2**

**4**

$PaX_{560}$-MeTz

**Compound 4.** In an amber vial, compound **S2** (8.0 mg, 0.018 mmol) and TSTU (11 mg, 0.036 mmol) were dissolved in DMF (100 µL) and DIPEA (23 mg, 0.18 mmol) was added and stirred for 30 min. at rt. (4-(6-Methyl-1,2,4,5-tetrazin-3-yl)phenyl)methanamine hydrochloride (5.4 mg, 0.027 mmol) was added and the reaction was stirred at rt for 2.5 h. The volatiles were removed *in vacuo*. The product was isolated by flash chromatography on a Biotage Isolera system (12g Interchim SiHP 30 µm cartridge, gradient 20% to 80% ethyl acetate/hexane) and freeze-dried from dioxane to yield 7.6 mg (68%) of **4** as an orange solid.

$^1$H NMR (400 MHz, Chloroform-*d*) δ 8.65 (t, *J* = 6.0 Hz, 1H), 8.52 – 8.47 (m, 2H), 7.76 (d, *J* = 8.9 Hz, 1H), 7.62 (d, *J* = 8.4 Hz, 2H), 7.04 (d, *J* = 15.4 Hz, 1H), 6.72 (d, *J* = 2.8 Hz, 1H), 6.69 (d, *J* = 2.8 Hz, 1H), 6.67 (d, *J* = 2.8 Hz, 1H), 6.58 (dd, *J* = 9.0, 2.8 Hz, 1H), 5.67 (dt, *J* = 15.0, 7.3 Hz, 1H), 4.72 (d, *J* = 6.0 Hz, 2H), 3.08 (s, 6H), 3.07 (s, 3H), 3.01 (s, 6H), 2.55 – 2.49 (m, 2H), 2.31 – 2.24 (m, 2H), 2.00 – 1.90 (m, 2H), 0.43 (s, 6H).

$^{13}$C NMR (101 MHz, Chloroform-*d*) δ 188.0, 174.4, 167.2, 164.2, 151.3, 151.0, 144.8, 144.2, 141.9, 139.5, 137.3, 131.4, 131.2, 130.5, 128.9, 128.3, 128.2, 127.7, 114.3, 114.2, 113.9, 113.2, 43.4, 40.1, 34.5, 31.6, 25.1, 21.3, -0.8.

HRMS (ESI) m/z: $[M+H]^+$ Calcd for $C_{35}H_{41}N_7O_2Si$: 620.3164, found: 620.3163.

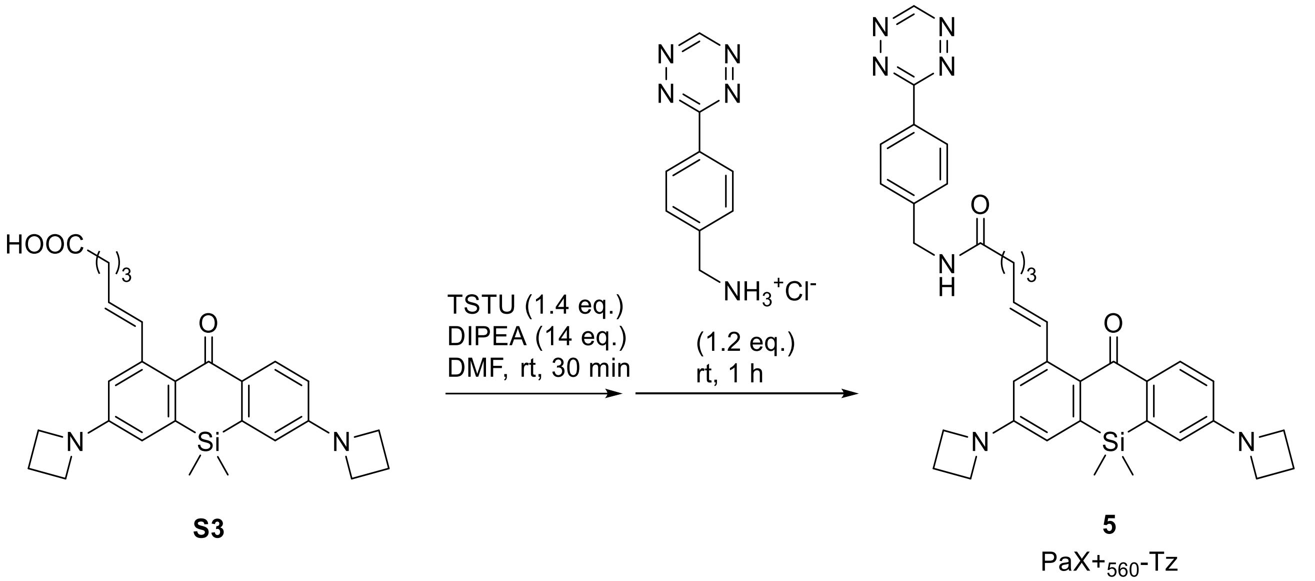

**Compound 5.** In an amber vial, compound **S3** (7.2 mg, 0.016 mmol) and TSTU (6.7 mg, 0.022 mmol) were dissolved in DMF (100 µL) and DIPEA (30 mg, 0.23 mmol) was added and stirred for 30 min. at rt. 3-(*p*-Benzylamino)-1,2,4,5-tetrazine hydrochloride (4.1 mg, 0.018 mmol) was added and the reaction was stirred at rt for 1 h. The volatiles were removed *in vacuo*. The product was isolated by flash chromatography on a Biotage Isolera system (12g Interchim SiHP 30 µm cartridge, gradient 50% to 100% ethyl acetate/hexane) and freeze-dried from dioxane to yield 2.1 mg (30%) of **6** as an orange solid.

$^{1}$H NMR (400 MHz, DMSO-d6) δ 10.57 (s, 1H), 8.50 (t, *J* = 6.0 Hz, 1H), 8.45 (d, *J* = 8.4 Hz, 2H), 7.89 (d, *J* = 8.7 Hz, 1H), 7.56 (d, *J* = 8.4 Hz, 2H), 7.09 (d, *J* = 15.7 Hz, 1H), 6.54 (d, *J* = 2.5 Hz, 1H), 6.53 (d, *J* = 2.5 Hz, 1H), 6.45 (dd, *J* = 8.7, 2.5 Hz, 1H), 6.42 (d, *J* = 2.5 Hz, 1H), 5.85 (dt, *J* = 15.5, 6.8 Hz, 1H), 4.44 (d, *J* = 5.9 Hz, 2H), 4.03 – 3.77 (m, 8H), 2.41 – 2.26 (m, 6H), 2.20 (q, *J* = 6.9 Hz, 2H), 1.77 (p, *J* = 7.3 Hz, 2H), 0.39 (s, 6H).

$^{13}$C NMR (101 MHz, DMSO-d6) δ 186.96, 172.34, 165.42, 158.09, 152.20, 151.75, 145.08, 142.38, 140.60, 138.27, 133.75, 131.86, 130.33, 128.77, 128.08, 127.81, 127.70, 113.25, 112.75, 111.94, 111.58, 51.46, 51.43, 41.89, 34.73, 32.01, 25.03, 16.16, -1.27.

HRMS (ESI) m/z: $[M+H]^+$ Calcd for $C_{36}H_{39}N_7O_2Si$: 630.3007, found: 630.3007.

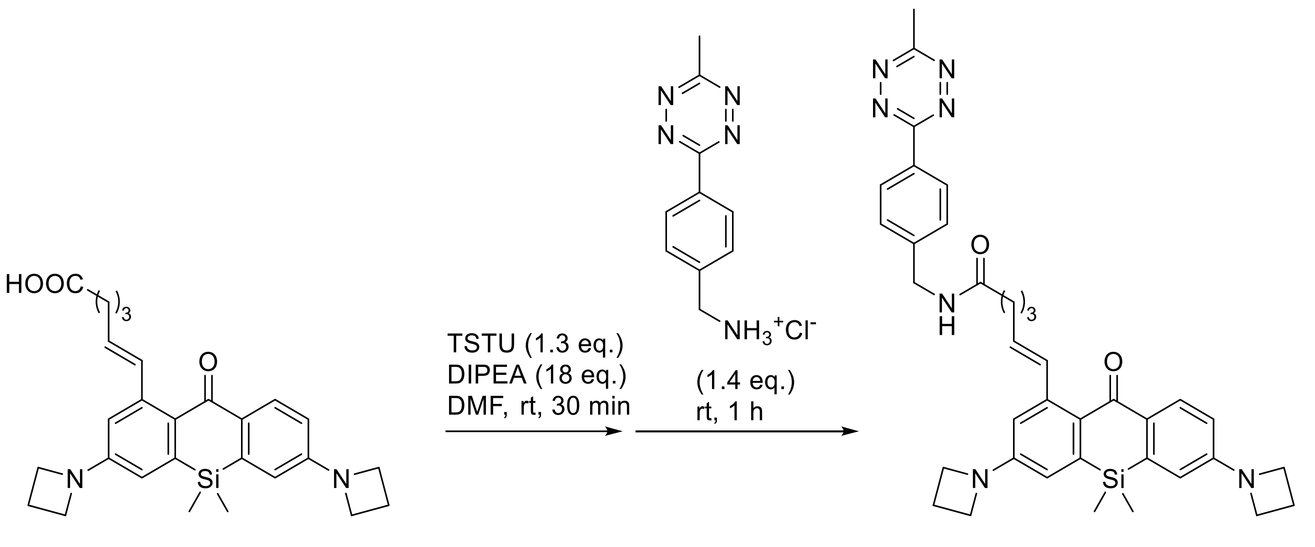

**S3**

**6**
PaX+$_{560}$-MeTz

**Compound 6.** In an amber vial, compound **S3** (6.1 mg, 0.013 mmol) and TSTU (5.2 mg, 0.017 mmol) were dissolved in DMF (100 µL) and DIPEA (30 mg, 0.23 mmol) was added and stirred for 30 min. at rt. (4-(6-Methyl-1,2,4,5-tetrazin-3-yl)phenyl)methanamine hydrochloride (4.1 mg, 0.018 mmol) was added and the reaction was stirred at rt for 1 h. The volatiles were removed *in vacuo*. The product was isolated by flash chromatography on a Biotage Isolera system (12g Interchim SiHP 30 µm cartridge, gradient 0% to 60% ethyl acetate/hexane) and freeze-dried from dioxane to yield 8.5 mg (97%, remainder dioxane) of **6** as a red solid.

$^{1}$H NMR (400 MHz, $CDCl_3$) δ 8.49 (d, *J* = 8.4 Hz, 1H), 7.72 (d, *J* = 8.7 Hz, 1H), 7.60 (d, *J* = 8.5 Hz, 2H), 6.97 (d, *J* = 15.4 Hz, 1H), 6.45 (d, *J* = 2.5 Hz, 2H), 6.39 (d, *J* = 2.5 Hz, 1H), 6.33 (dd, *J* = 8.7, 2.5 Hz, 1H), 5.66 (dt, *J* = 15.1, 7.3 Hz, 1H), 4.70 (d, *J* = 5.8 Hz, 2H), 4.03 (t, *J* = 7.3 Hz, 4H), 3.97 (t, *J* = 7.3 Hz, 4H), 2.56 – 2.47 (m, 2H), 2.48 – 2.37 (m, 4H), 2.32 – 2.22 (m, 2H), 2.03 – 1.90 (m, 2H), 0.40 (s, 6H).

$^{13}$C NMR (101 MHz, Chloroform-*d*) δ 188.2, 174.4, 167.2, 164.2, 152.4, 152.1, 144.7, 144.0, 141.7, 139.3, 136.8, 132.2, 131.0, 130.5, 128.9, 128.5, 128.4, 128.2, 113.2, 112.9, 112.8, 112.0, 51.8, 43.4, 34.5, 31.6, 25.1, 21.3, 16.8, -1.0.

HRMS (ESI) m/z: $[M+H]^+$ Calcd for $C_{37}H_{41}N_7O_2Si$: 644.3164, found: 644.3153.

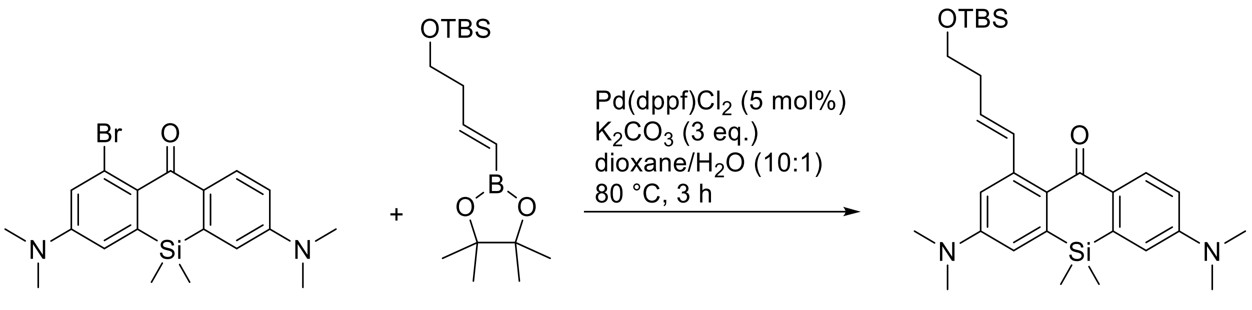

**Compound S6.** In a 10 mL reaction tube, compound **S4** (81 mg, 0.20 mmol; prepared according to literature procedure[15]), compound **S5** (94 mg, 0.30 mmol, 1.5 equiv; prepared according to literature procedure [16]), $K_2CO_3$ (82 mg, 0.60 mmol, 3.0 equiv) and Pd(dppf)$Cl_2 \cdot CH_2Cl_2$ (8.2 mg, 10 µmol, 5 mol%) were loaded. Dioxane (2 mL) and water (200 µL) were added. The mixture was sparged with argon for 30 min and then stirred at 80°C for 20 h. Upon cooling, the reaction mixture was diluted with ethyl acetate (20 mL) and washed with brine (20 mL). The organics were dried over $Na_2SO_4$, filtered, evaporated. The product was isolated by flash chromatography on Biotage Isolera system (25 g Interchim SiHP 30 µm cartridge, gradient 0% to 30% ethyl acetate/hexane) and freeze-dried from dioxane to yield 87 mg (85%) of **S6** as a yellow solid.

$^1$H NMR (400 MHz, Chloroform-*d*) δ 8.24 (d, *J* = 8.9 Hz, 1H), 7.34 (d, *J* = 15.5 Hz, 1H), 6.82 (dd, *J* = 8.9, 2.8 Hz, 1H), 6.80 – 6.74 (m, 3H), 5.95 (dt, *J* = 15.5, 7.0 Hz, 1H), 3.81 (t, *J* = 6.8 Hz, 2H), 3.09 (s, 6H), 3.07 (s, 6H), 2.55 (m, 2H), 0.92 (s, 9H), 0.46 (s, 6H), 0.09 (s, 6H).

$^{13}$C NMR (101 MHz, Chloroform-*d*) δ 188.1, 151.2, 150.8, 143.7, 141.3, 139.0, 135.9, 132.4, 131.4, 128.4, 126.3, 114.3, 113.9, 113.8, 113.4, 63.6, 40.2, 40.1, 36.8, 26.1, 18.5, -0.9, -5.0.

HRMS (ESI) m/z: $[M+H]^+$ Calcd for $C_{29}H_{44}N_2O_2Si_2$: 509.3014, found: 509.3013.

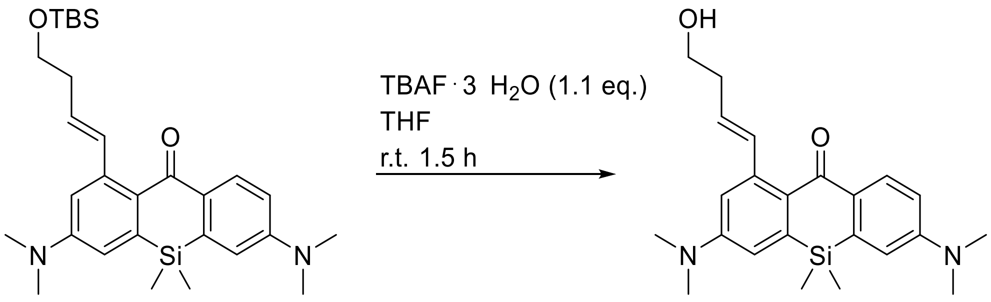

**S6** **S7**

**Compound S7.** A solution of compound **S6** (51 mg, 0.1 mmol) in THF (2 mL) was cooled in ice-water bath and tetrabutylammonium fluoride trihydrate (TBAF; 35 mg, 0.11 mmol) was added. After stirring for 45 min, the solution was warmed to rt and stirred for 1.5 hr. Sat. aq. $NH_4Cl$ (20 mL) was added and the reaction mixture was extracted with ethyl acetate (2 × 10 mL). The combined extracts were washed with brine (20 mL), dried over $Na_2SO_4$, filtered, evaporated, the product was isolated by flash chromatography on a Biotage Isolera system (12g Interchim SiHP 30 µm cartridge, gradient 20% to 80% ethyl acetate/hexane) and freeze-dried from 1,4-dioxane to yield 39 mg (99%) of **S7** as a yellow solid.

$^{1}H$ NMR (400 MHz, Chloroform-*d*) δ 8.26 (d, *J* = 8.9 Hz, 1H), 7.24 (d, *J* = 15.7 Hz, 1H), 6.80 (dd, *J* = 9.0, 2.8 Hz, 1H), 6.75 (d, *J* = 2.8 Hz, 2H), 6.71 – 6.66 (m, 1H), 5.76 (dt, *J* = 15.7, 7.2 Hz, 1H), 3.80 (t, *J* = 5.5 Hz, 2H), 3.35 (s, 1H), 3.10 (s, 6H), 3.08 (s, 6H), 2.50 (qd, *J* = 6.1, 1.1 Hz, 2H), 0.45 (s, 6H).

$^{13}C$ NMR (101 MHz, Chloroform-*d*) δ 187.7, 151.3, 150.9, 144.0, 141.6, 139.2, 138.9, 131.8, 131.8, 128.1, 124.7, 114.3, 113.9, 113.8, 113.4, 62.0, 40.2, 40.1, 35.9, -0.8.

HRMS (ESI) m/z: $[M+H]^+$ Calcd for $C_{23}H_{30}N_2O_2Si$: 395.2149, found: 395.2147.

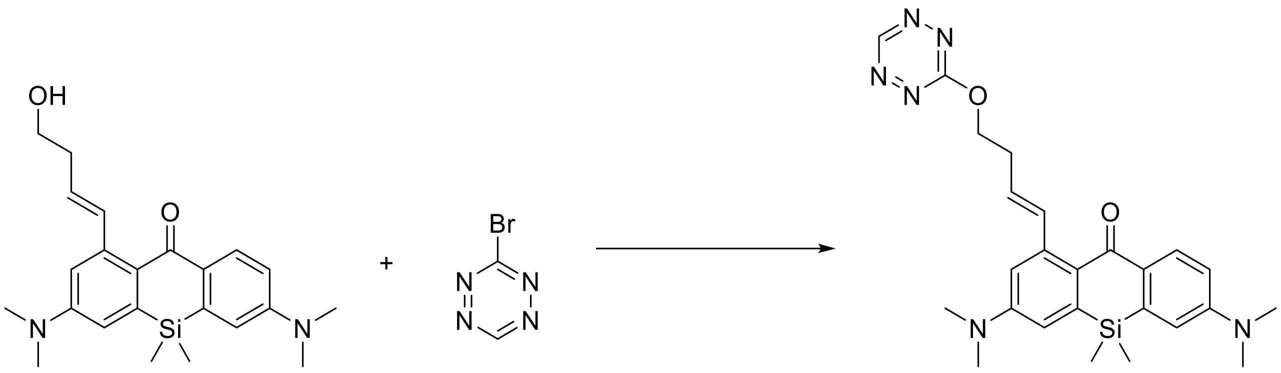

**S7** **7**

**Compound 7.** In an amber vial, compound **S7** (7.9 mg, 0.02 mmol), 3-bromo-1,2,4,5-tetrazine (16 mg, 0.1 mmol) were dissolved in DMF (200 µL) and 2,4,6-collidine (34 mg, 0.20 mmol) was added and stirred for 1 h at rt. The volatiles were removed *in vacuo*. The product was isolated by flash chromatography on a Biotage Isolera system (12g Interchim SiHP 30 µm cartridge, gradient 20% to 80% ethyl acetate/hexane) and freeze-dried from dioxane to yield 1.7 mg (18%) of **7** as an orange solid.

HRMS (ESI) m/z: $[M+H]^+$ Calcd for $C_{25}H_{30}N_6O_2Si$: 475.2272, found: 475.2270.

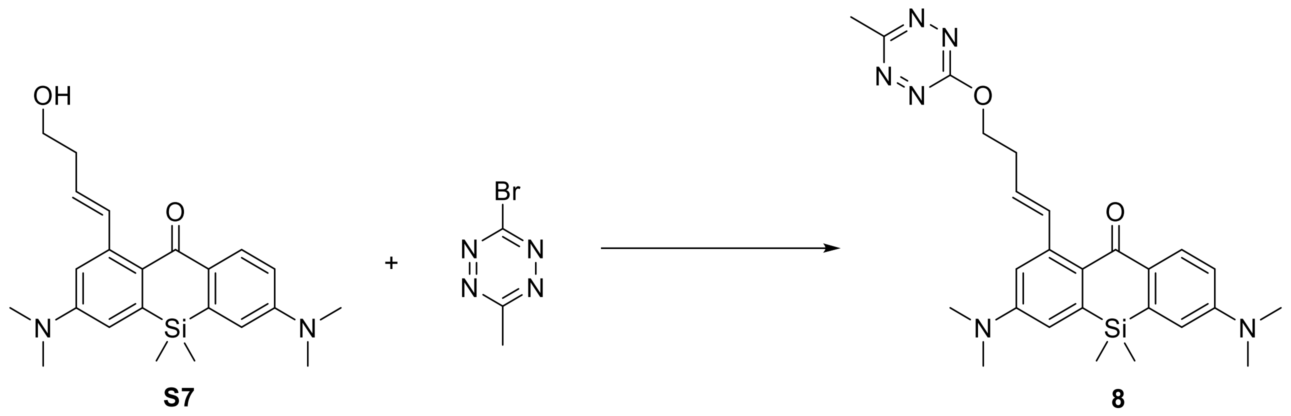

**Compound 8.** In an amber vial, compound **S7** (6.4 mg, 0.016 mmol), 3-bromo-1,2,4,5-tetrazine (7.6 mg, 0.043 mmol) were dissolved in MeCN (800 µL) and 2,4,6-collidine (6.6 mg, 0.054 mmol) was added and stirred for 16 h at rt. The volatiles were removed *in vacuo*. The product was isolated by flash chromatography on a Biotage Isolera system (12g Interchim SiHP 30 µm cartridge, gradient 0% to 100% ethyl acetate/hexane) and freeze-dried from dioxane to yield 5.1 mg (65%) of **7** as a red solid.

HRMS (ESI) m/z: $[M+H]^+$ Calcd for $C_{26}H_{32}N_6O_2Si$: 489.2429, found: 489.2430.

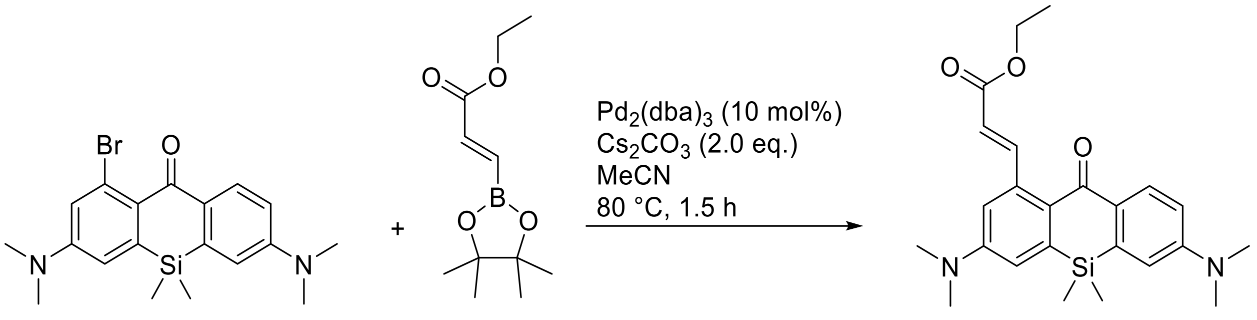

S4 S8

**Compound S8.** In a 10 mL reaction tube, compound **S4** (81 mg, 0.20 mmol; prepared according to literature procedure[15]), (*E*)-Ethyl 3-(4,4,5,5-tetramethyl-1,3,2-dioxaborolan-2-yl)acrylate (68 mg, 0.30 mmol, 1.5 equiv), $Cs_2CO_3$ (98 mg, 0.30 mmol, 1.5 equiv), $Pd_2(dba)_3$ (19 mg, 20 µmol, 10 mol%) and Xphos (19 mg, 40 µmol, 20 mol%) were loaded. MeCN was added (2 mL). The mixture was stirred at 80°C for 1.5 h. Upon cooling, the reaction mixture was poured into sat. aq. $NH_4Cl$ (20 mL) and extracted with ethyl acetate (3 × 20 mL) and washed with brine (20 mL). The combined organics were washed with brine, dried over $Na_2SO_4$, filtered, and evaporated. The product was isolated by flash chromatography on Biotage Isolera system (12 g Interchim SiHP 30 µm cartridge, gradient 0% to 50% ethyl acetate/hexane) and freeze-dried from dioxane to yield 62 mg (73%) of **S8** as a yellow solid.

$^1$H NMR (400 MHz, Chloroform-*d*) δ 8.51 (d, *J* = 15.5 Hz, 1H), 8.30 (d, *J* = 9.0 Hz, 1H), 6.85 – 6.81 (m, 2H), 6.77 (d, *J* = 2.8 Hz, 1H), 6.71 (d, *J* = 2.8 Hz, 1H), 6.08 (d, *J* = 15.6 Hz, 1H), 4.29 (q, *J* = 7.1 Hz, 2H), 3.10 (s, 6H), 3.09 (s, 6H), 1.36 (t, *J* = 7.1 Hz, 3H), 0.47 (s, 6H).

$^{13}$C NMR (101 MHz, Chloroform-*d*) δ 186.8, 167.6, 151.5, 151.0, 150.8, 141.9, 141.3, 139.2, 131.8, 131.0, 129.1, 117.4, 115.6, 114.2, 113.9, 113.5, 60.4, 40.2, 40.2, 14.6, -0.8.

HRMS (ESI) m/z: $[M+H]^+$ Calcd for $C_{24}H_{30}N_2O_3Si$: 423.2098, found: 423.2094.

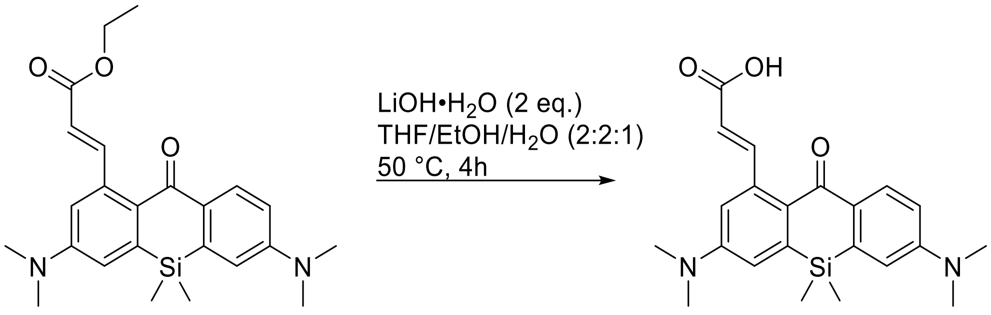

**S8** **S9**

**Compound S9.** To a solution of **S8** (54 mg, 0.13 mmol) in THF (1 mL) and ethanol (1 mL) was added dropwise a solution of lithium hydroxide monohydrate (11 mg. 0.26 mmol) in $H_2O$ (500 µL). The solution was was placed in a 50 °C oil bath and stirred for 4 h. The mixture was then cooled down to rt and poured into 20 mL of water. The pH was adjusted to 2 with HCl (1 M) and extracted with DCM (4 × 10 mL). The combined extracts were washed with brine (20 mL), dried over $Na_2SO_4$, filtered, and evaporated. The product was isolated by preparative HPLC (Interchim Uptisphere Strategy PhC4 250×21.2 mm 5 µm, solvent flow rate 18 mL/min, gradient 40% to 70% A/B, A: acetonitrile + 0.1% (v/v) formic acid, B: water + 0.1% (v/v) formic acid) and freeze-dried from dioxane to give 45 mg (89%) of **S9** as a pink solid.

$^1$H NMR (400 MHz, DMSO-$d_6$) δ 12.24 (s, 1H), 8.25 (d, $J$ = 15.6 Hz, 1H), 8.02 (d, $J$ = 8.8 Hz, 1H), 6.96 (d, $J$ = 2.8 Hz, 1H), 6.89 (d, $J$ = 2.7 Hz, 1H), 6.86 (dd, $J$ = 8.9, 2.8 Hz, 1H), 6.75 (d, $J$ = 2.7 Hz, 1H), 6.06 (d, $J$ = 15.5 Hz, 1H), 3.08 (s, 6H), 3.05 (s, 6H), 0.46 (s, 6H).

$^{13}$C NMR (101 MHz, DMSO-$d_6$) δ 185.8, 168.0, 151.3, 150.6, 149.6, 141.4, 139.9, 138.8, 130.7, 129.9, 127.6, 117.8, 116.0, 114.1, 113.5, 113.2, -1.1.

HRMS (ESI) m/z: $[M+H]^+$ Calcd for $C_{22}H_{26}N_2O_3Si$: 395.1785, found: 395.1784.

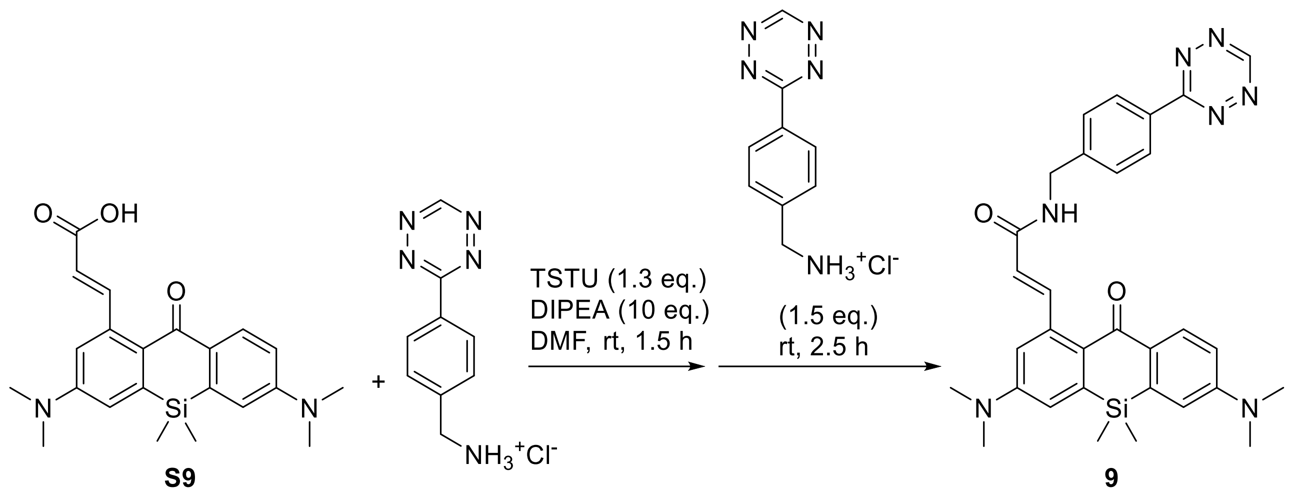

**Compound 9.** In a pear-shaped flask, compound **S9** (9.3 mg, 0.024 mmol) and TSTU (9.4 mg, 0.031 mmol) were dissolved in DMF (100 µL) and DIPEA (31 mg, 0.24 mmol) was added and stirred for 1.5 h at rt. 3-(*p*-Benzylamino)-1,2,4,5-tetrazine hydrochloride (8.2 mg, 0.037 mmol) was added and the reaction was stirred at rt for 2.5 h. The volatiles were removed *in vacuo*. The product was isolated by flash chromatography on a Biotage Isolera system (12g Interchim SiHP 30 µm cartridge, gradient 50% to 80% ethyl acetate/hexane) and freeze-dried from dioxane to yield 3.7 mg (27%) of **9** as an orange solid.

HRMS (ESI) m/z: $[M+H]^+$ Calcd for $C_{31}H_{33}N_7O_2Si$: 564.2538, found: 564.2530.

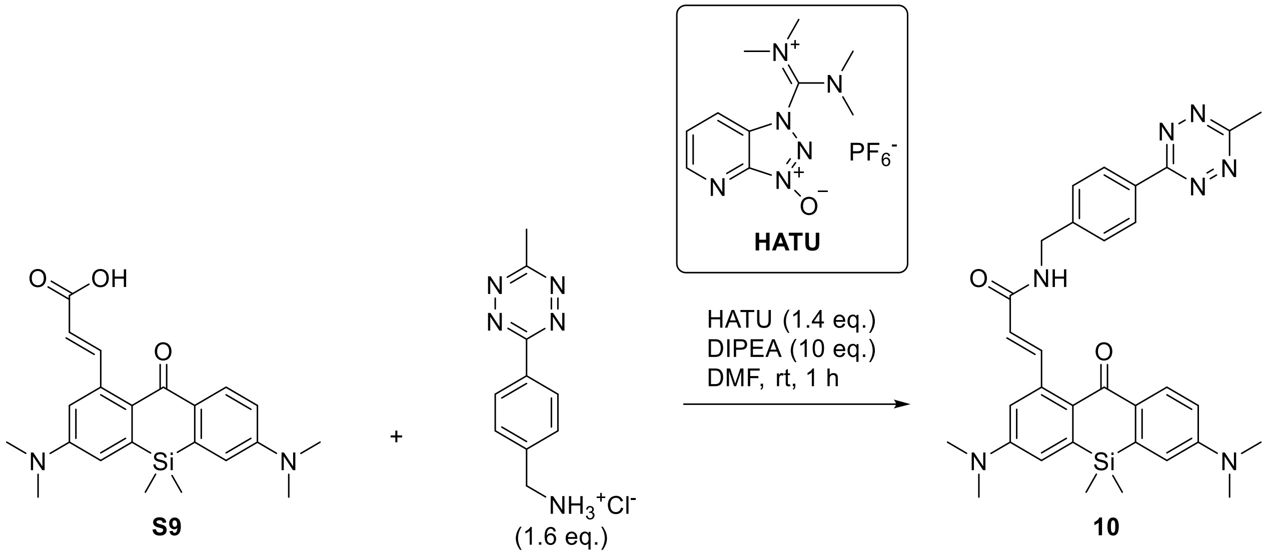

**Compound 10.** In an amber vial, compound **S9** (10 mg, 0.025 mmol) 3-(*p*-Benzylamino)-1,2,4,5-tetrazine hydrochloride (7.9 mg, 0.040 mmol) and HATU (13 mg, 0.034 mmol) were dissolved in DMF (200 µL) and DIPEA (32 mg, 0.25 mmol) was added and stirred for 1 h at rt. The volatiles were removed *in vacuo*. The product was isolated by flash chromatography on a Biotage Isolera system (12g Interchim SiHP 30 µm cartridge, gradient 40% to 90% ethyl acetate/hexane) and freeze-dried from dioxane to yield 11.1 mg (65%, remainder dioxane) of **10** as an orange solid.

$^1$H NMR (400 MHz, DMSO-$d_6$) δ 8.67 (t, *J* = 6.0 Hz, 1H), 8.46 (d, *J* = 8.4 Hz, 2H), 8.16 (d, *J* = 15.4 Hz, 1H), 8.02 (d, *J* = 8.8 Hz, 1H), 7.61 (d, *J* = 8.3 Hz, 2H), 6.96 (d, *J* = 2.8 Hz, 1H), 6.91 – 6.83 (m, 2H), 6.75 (d, *J* = 2.8 Hz, 1H), 6.30 (d, *J* = 15.4 Hz, 1H), 4.55 (d, *J* = 5.9 Hz, 2H), 3.08 (s, 6H), 3.04 (s, 6H), 2.99 (s, 3H), 0.46 (s, 6H).

$^{13}$C NMR (101 MHz, DMSO-$d_6$) δ 185.9, 167.1, 165.7, 163.2, 151.3, 150.6, 144.7, 144.4, 141.3, 140.7, 138.7, 130.6, 130.4, 130.1, 128.4, 127.8, 127.5, 120.8, 115.7, 114.1, 113.2, 42.1, 39.5, 39.4, 20.8, -1.1.

HRMS (ESI) m/z: $[M+H]^+$ Calcd for $C_{32}H_{35}N_7O_2Si$: 578.2694, found: 578.2691.

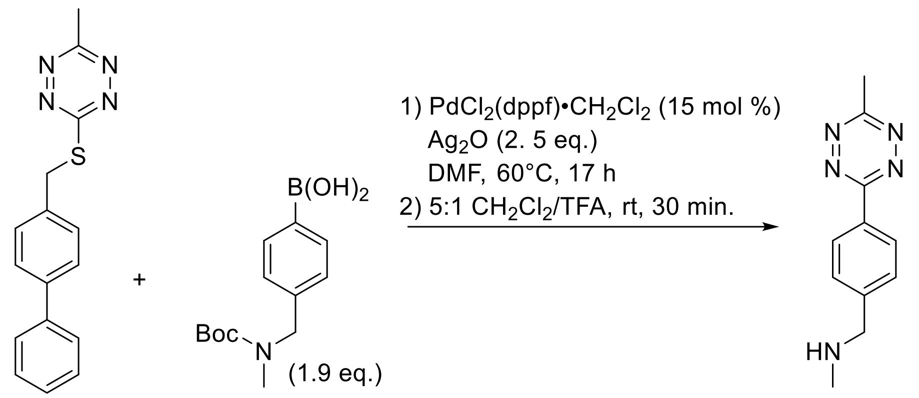

**S10**

**Compound S10.** In a 10 mL reaction tube, 3-(([1,1′-Biphenyl]-4-ylmethyl)thio)-6-methyl-1,2,4,5-tetrazine (29 mg, 0.1 mmol, known compound[17]), (4-{[(tert-Butoxycarbonyl)(methyl)amino]methyl}phenyl)boronic acid (50 mg, 0.19 mmol, abcr catalog no. AB266066), $PdCl_2(dppf)•CH_2Cl_2$ (12 mg, 15 µmol, 15 mol%), and $Ag_2O$ (58 mg, 0.25 mmol) were loaded. DMF was added, the headspace was flushed with argon, and the tube was sealed. The mixture was stirred at 60 °C for 17 h. Upon cooling, the volatiles were removed *in vacuo*. The residue was resuspended in ethyl acetate and filtered through silica, washing with ethyl acetate. The product was isolated by flash chromatography on a Biotage Isolera system (12g Interchim SiHP 30 µm cartridge, gradient 0% to 100% A/B; A: 10% ethyl acetate in dichloromethane, B: dichloromethane). The resulting red oil was used in the next step without further purification.

In a 10 mL pear-shaped flask, the red oil was dissolved in dichloromethane (1 mL) and trifluoroacetic acid (200 µL) was added dropwise. The resulting reaction mixture was stirred for 30 minutes at rt. The volatiles were removed *in vacuo* by coevaporation with toluene (3 × 10 ml). The product was freeze-dried from dioxane to give 27 mg (~100%, remainder dioxane/$H_2O$/DCM/ethyl acetate) of **S10** as a red solid.[18]

$^1$H NMR (400 MHz, DMSO-$d_6$) δ 8.47 (d, $J$ = 8.3 Hz, 2H), 7.68 (d, $J$ = 8.4 Hz, 2H), 4.03 (s, 2H), 3.00 (s, 3H), 2.47 (s, 3H).

$^{13}$C NMR (101 MHz, DMSO-$d_6$) δ 167.2, 163.1, 140.9, 131.3, 129.8, 127.5, 52.7, 33.9, 20.8.

HRMS (ESI) m/z: $[M+H]^+$ Calcd for $C_{11}H_{13}N_5$: 216.1244, found: 216.1244.

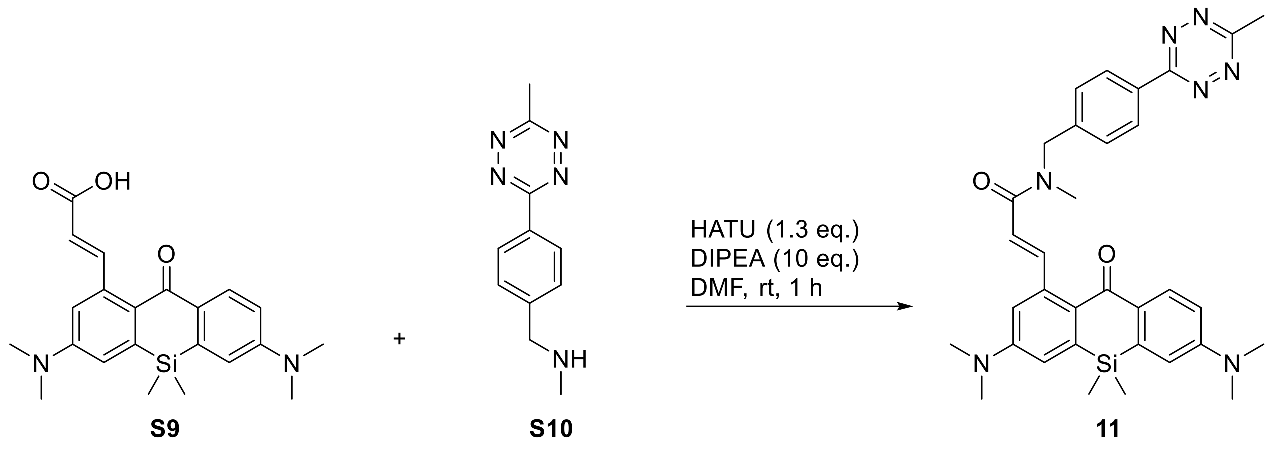

**Compound 11.** In an amber vial, compound **S9** (10 mg, 0.025 mmol) **S10** (8 mg, 0.038 mmol) and HATU (12 mg, 0.033 mmol) were dissolved in DMF (200 µL) and DIPEA (32 mg, 0.25 mmol) was added and stirred for 1 h at rt. The volatiles were removed *in vacuo*. The product was isolated by flash chromatography on a Biotage Isolera system (12g Interchim SiHP 30 µm cartridge, gradient 30% to 90% ethyl acetate/hexane) and freeze-dried from dioxane to yield 8.7 mg (59%) of **10** as an orange solid.

HRMS (ESI) m/z: $[M+H]^+$ Calcd for $C_{33}H_{37}N_7O_2Si$: 592.2851, found: 592.2853.

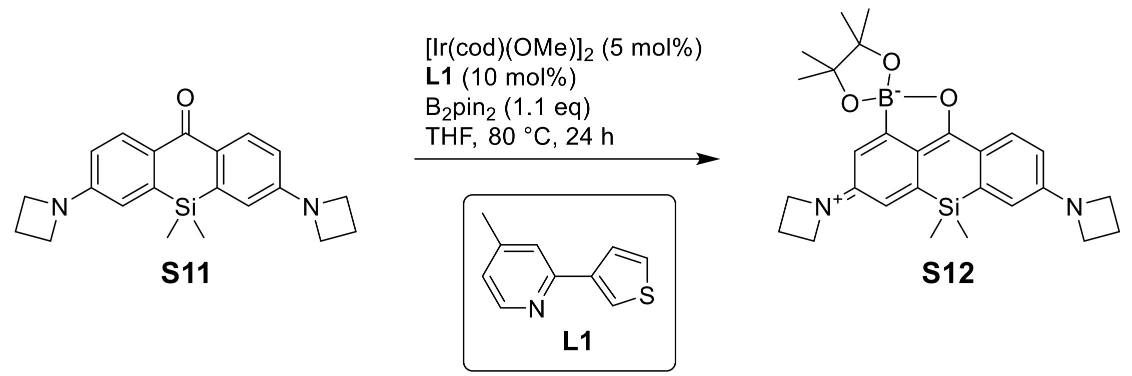

**Compound S12.** A mixture of **S11** (157 mg, 0.45 mmol), (1,5-cyclooctadiene)(methoxy)iridium(I) dimer (14.9 mg, 22.5 µmol, 5 mol%), ligand **L1** (7.9 mg, 45.0 µmol, 10 mol%; known compound: [19]) and bis(pinacolato)diboron (126 mg, 0.50 mmol, 1.1 equiv) in degassed THF (5 mL) was stirred at 80 °C for 24 h. The reaction mixture was evaporated on Celite, the product was isolated by flash column chromatography (12 g Interchim SiHP 30 µm cartridge, gradient 5% to 30% EtOAc/$CH_2Cl_2$) and freeze-dried from 1,4-dioxane to yield 142 mg (67%) of **S12** as bright orange solid.[20]

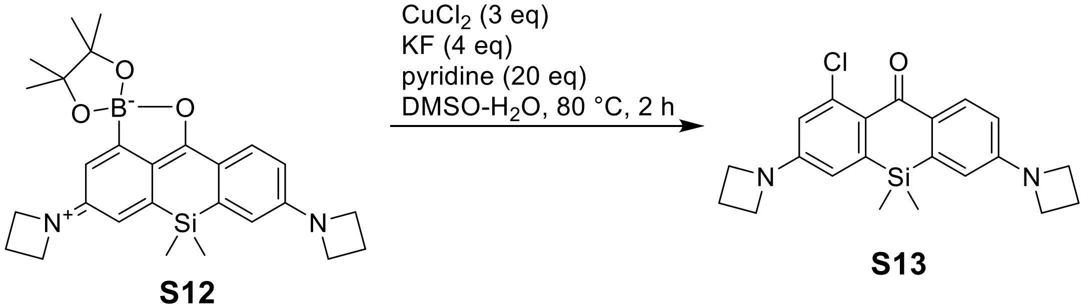

**Compound S13.** Anhydrous copper(II) chloride (121 mg, 0.9 mmol, 4.5 equiv) and potassium fluoride (70 mg, 1.2 mmol, 4 equiv) were added to **S12** (142 mg, 0.3 mmol), followed by DMSO (2 mL), pyridine (0.48 mL, 6 mmol, 20 equiv) and water (0.2 mL). The resulting mixture was stirred at 80 °C for 2 h, cooled to rt, poured into water (100 mL), and the product was extracted with $CHCl_3$ (3×50 mL). The combined extracts were washed with brine and dried over $Na_2SO_4$, the product was isolated by flash column chromatography (12 g Interchim SiHP 30 µm cartridge, gradient 0% to 30% EtOAc/hexane with 30% $CH_2Cl_2$ constant additive) and freeze-dried from 1,4-dioxane to yield 90 mg (78%) of **S13** as bright yellow solid.

$^1$H NMR (400 MHz, $CDCl_3$): δ 8.14 (d, *J* = 8.7 Hz, 1H), 6.51 (dd, *J* = 8.7, 2.5 Hz, 1H), 6.49 (d, *J* = 2.4 Hz, 1H), 6.44 (d, *J* = 2.5 Hz, 1H), 6.39 (d, *J* = 2.4 Hz, 1H), 4.01 (td, *J* = 7.3 Hz, 4H), 4.00 (td, *J* = 7.3 Hz, 4H), 2.48 – 2.37 (m, 4H), 0.43 (s, 6H).

$^{13}$C NMR (101 MHz, $CDCl_3$): δ 186.7, 152.5, 152.1, 142.4, 138.2, 136.8, 133.2, 131.2, 127.6, 115.3, 112.6, 112.5, 112.4, 51.9, 51.8, 16.84, 16.75, -1.2.

HRMS ($C_{21}H_{23}ClN_2OSi$): *m/z* (positive mode) = 383.1341 (found $[M+H]^+$), 383.1341 (calc.).

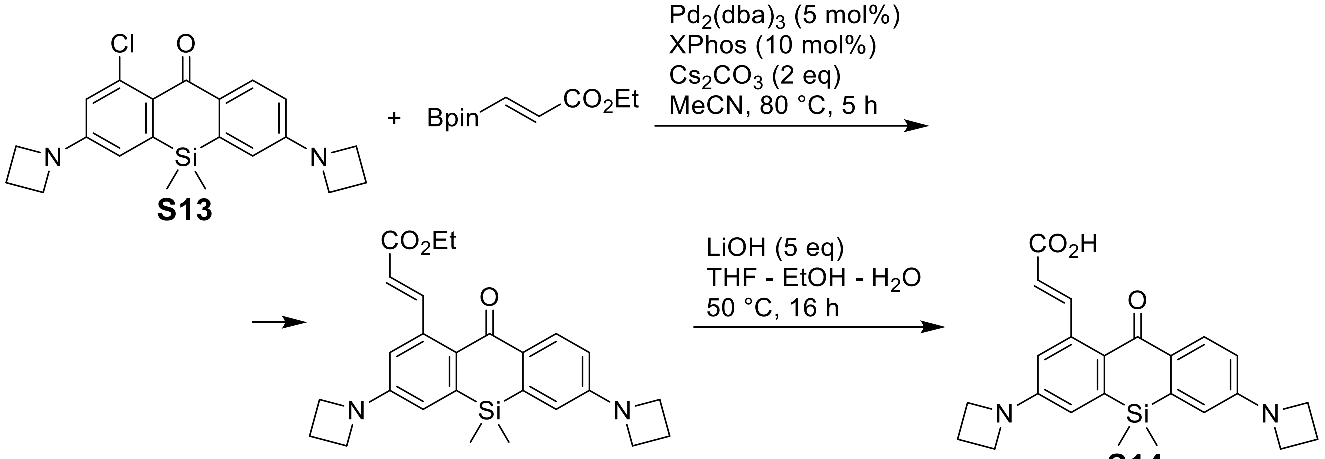

**S14**

**Compound S14.** A mixture of **S13** (20 mg, 52.2 µmol), (*E*)-2-(ethoxycarbonyl)vinylboronic acid pinacol ester (18 mg, 78.3 mmol, 1.5 equiv), tris(dibenzylideneacetone)dipalladium(0) (2.4 mg, 2.61 µmol, 5 mol%), XPhos (2.5 mg, 5.22 µmol, 10 mol%) and cesium carbonate (34 mg, 104 µmol, 2 equiv) in acetonitrile (1 mL) was stirred at 80 °C for 5 h. Upon cooling, the reaction mixture was diluted with brine (50 mL) and extracted with $CH_2Cl_2$ (3×20 mL). The combined extracts were dried over $Na_2SO_4$, filtered, the filtrate was evaporated and the crude ethyl ester was used directly in the next step.

Lithium hydroxide solution (11 mg of $LiOH{\cdot}H_2O$ in 0.4 mL water, 260 µmol, 5 equiv) was added to a stirred solution of the crude ethyl ester in THF (2 mL) and ethanol (0.4 mL), and the reaction mixture was stirred at 50 °C for 16 h. Acetic acid (30 µL) was then added, and the reaction mixture was evaporated to dryness. The product was isolated by flash column chromatography (12 g Interchim SiHP 30 µm cartridge, gradient 0% to 10% 2-propanol/$CH_2Cl_2$) and freeze-dried from aqueous 1,4-dioxane to yield 7 mg (32%) of **S14** as orange solid.

$^{1}$H NMR (400 MHz, DMSO-$d_6$): δ 12.18 (br.s, 1H), 8.17 (d, *J* = 15.5 Hz, 1H), 7.99 (d, *J* = 8.7 Hz, 1H), 6.65 (d, *J* = 2.5 Hz, 1H), 6.58 (d, *J* = 2.5 Hz, 1H), 6.52 (dd, *J* = 8.7, 2.5 Hz, 1H), 6.46 (d, *J* = 2.5 Hz, 1H), 6.02 (d, *J* = 15.5 Hz, 1H), 4.01 (t, *J* = 7.4 Hz, 4H), 3.96 (t, *J* = 7.3 Hz, 4H), 2.45 – 2.30 (m, 4H), 0.42 (s, 6H).

$^{13}$C NMR (101 MHz, DMSO-$d_6$): δ 186.0, 152.4, 151.7, 141.2, 139.8, 138.6, 130.7, 130.5, 128.4, 114.8, 113.0, 112.3, 112.1, 51.5, 29.04, 29.01, 16.1, -1.3.

HRMS ($C_{24}H_{26}N_2O_3Si$): *m/z* (positive mode) = 419.1786 (found $[M+H]^+$), 419.1785 (calc.).

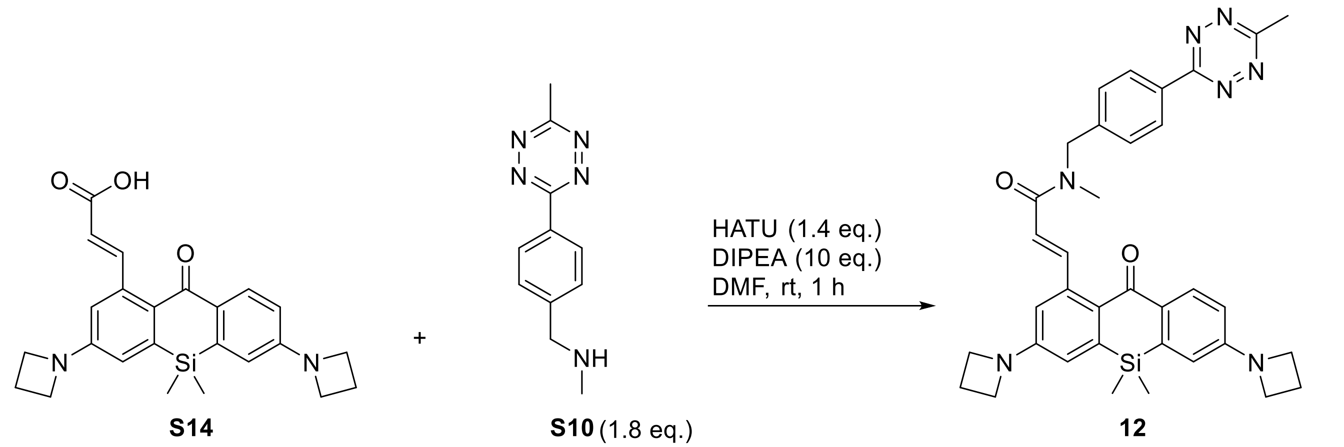

**Compound 12.** In an amber vial, compound **S14** (7.0 mg, 0.017 mmol) **S10** (6.5 mg, 0.030 mmol) and HATU (9.2 mg, 0.024 mmol) were dissolved in DMF (200 µL) and DIPEA (22 mg, 0.17 mmol) was added and stirred for 1 h at rt. The volatiles were removed *in vacuo*. The product was isolated by flash chromatography on a Biotage Isolera system (12g Interchim SiHP 30 µm cartridge, gradient 30% to 90% ethyl acetate/hexane) and freeze-dried from dioxane to yield 5.7 mg (54%) of **10** as an orange solid.

HRMS (ESI) m/z: $[M+H]^+$ Calcd for $C_{35}H_{37}N_7O_2Si$: 616.2851, found: 616.2854.

### SUPPLEMENTARY REFERENCES

1. Nikic, I.; Estrada Girona, G.; Kang, J. H.; Paci, G.; Mikhaleva, S.; Koehler, C.; Shymanska, N. V.; Ventura Santos, C.; Spitz, D.; Lemke, E. A., Debugging Eukaryotic Genetic Code Expansion for Site-Specific Click-PAINT Super-Resolution Microscopy. *Angew Chem Int Ed Engl* **2016,** *55* (52), 16172-16176.

2. Markwardt, M. L.; Kremers, G. J.; Kraft, C. A.; Ray, K.; Cranfill, P. J.; Wilson, K. A.; Day, R. N.; Wachter, R. M.; Davidson, M. W.; Rizzo, M. A., An improved cerulean fluorescent protein with enhanced brightness and reduced reversible photoswitching. *PLoS One* **2011,** *6* (3), e17896.

3. Gregor, C.; Grimm, F.; Rehman, J.; Wurm, C. A.; Egner, A., Two-color live-cell STED nanoscopy by click labeling with cell-permeable fluorophores. *bioRxiv* **2022**, 2022.09.11.507450.

4. Uno, K.; Bossi, M. L.; Konen, T.; Belov, V. N.; Irie, M.; Hell, S. W., Asymmetric Diarylethenes with Oxidized 2-Alkylbenzothiophen-3-yl Units: Chemistry, Fluorescence, and Photoswitching. *Advanced Optical Materials* **2019,** *7* (6), 1801746.

5. Uno, K.; Aktalay, A.; Bossi, M. L.; Irie, M.; Belov, V. N.; Hell, S. W., Turn-on mode diarylethenes for bioconjugation and fluorescence microscopy of cellular structures. *Proc. Natl. Acad. Sci. U. S. A.* **2021,** *118* (14), e2100165118.

6. Schmidt, R.; Weihs, T.; Wurm, C. A.; Jansen, I.; Rehman, J.; Sahl, S. J.; Hell, S. W., MINFLUX nanometer-scale 3D imaging and microsecond-range tracking on a common fluorescence microscope. *Nat Commun* **2021,** *12* (1), 1478.

7. Zhang, Y.; Fang, C.; Wang, R. E.; Wang, Y.; Guo, H.; Guo, C.; Zhao, L.; Li, S.; Li, X.; Schultz, P. G.; Cao, Y. J.; Wang, F., A tumor-targeted immune checkpoint blocker. *Proc. Natl. Acad. Sci. U. S. A.* **2019,** *116* (32), 15889-15894.

8. Werther, P.; Yserentant, K.; Braun, F.; Kaltwasser, N.; Popp, C.; Baalmann, M.; Herten, D. P.; Wombacher, R., Live-Cell Localization Microscopy with a Fluorogenic and Self-Blinking Tetrazine Probe. *Angew Chem Int Ed Engl* **2020,** *59* (2), 804-810.

9. Nikic, I.; Plass, T.; Schraidt, O.; Szymanski, J.; Briggs, J. A.; Schultz, C.; Lemke, E. A., Minimal tags for rapid dual-color live-cell labeling and super-resolution microscopy. *Angew Chem Int Ed Engl* **2014,** *53* (8), 2245-9.

10. Frisch, M.; Trucks, G.; Schlegel, H.; Scuseria, G.; Robb, M.; Cheeseman, J.; Scalmani, G.; Barone, V.; Petersson, G.; Nakatsuji, H., Gaussian 16, Revision B. 01, Gaussian. *Inc., Wallingford CT* **2016,** *3*.

11. Chai, J.-D.; Head-Gordon, M., Long-range corrected hybrid density functionals with damped atom–atom dispersion corrections. *Physical Chemistry Chemical Physics* **2008,** *10* (44), 6615-6620.

12. Weigend, F.; Ahlrichs, R., Balanced basis sets of split valence, triple zeta valence and quadruple zeta valence quality for H to Rn: Design and assessment of accuracy. *Physical Chemistry Chemical Physics* **2005,** *7* (18), 3297-3305.

13. Marenich, A. V.; Cramer, C. J.; Truhlar, D. G., Universal Solvation Model Based on Solute Electron Density and on a Continuum Model of the Solvent Defined by the Bulk Dielectric Constant and Atomic Surface Tensions. *The Journal of Physical Chemistry B* **2009,** *113* (18), 6378-6396.

14. Chi, W. J.; Huang, L.; Wang, C.; Tan, D.; Xu, Z. C.; Liu, X. G., A unified fluorescence quenching mechanism of tetrazine-based fluorogenic dyes: energy transfer to a dark state. *Materials Chemistry Frontiers* **2021,** *5* (18), 7012-7021.

15. Lincoln, R.; Bossi, M. L.; Remmel, M.; D'Este, E.; Butkevich, A. N.; Hell, S. W., A general design of caging-group-free photoactivatable fluorophores for live-cell nanoscopy. *Nat. Chem.* **2022,** *14* (9), 1013-1020.

16. Wang, Y. N. D.; Kimball, G.; Prashad, A. S.; Wang, Y., Zr-mediated hydroboration: stereoselective synthesis of vinyl boronic esters. *Tetrahedron Lett.* **2005,** *46* (50), 8777-8780.

17. Lambert, W. D.; Fang, Y.; Mahapatra, S.; Huang, Z.; am Ende, C. W.; Fox, J. M., Installation of Minimal Tetrazines through Silver-Mediated Liebeskind–Srogl Coupling with Arylboronic Acids. *Journal of the American Chemical Society* **2019,** *141* (43), 17068-17074.

18. Li, X.; Wang, Y. Preparation of tetrazine-based fluorescent probes for detection of superoxide anion. CN115583920, 2023.

19. Hoque, M. E.; Hassan, M. M. M.; Chattopadhyay, B., Remarkably Efficient Iridium Catalysts for Directed C(sp(2))-H and C(sp(3))-H Borylation of Diverse Classes of Substrates. *J. Am. Chem. Soc.* **2021,** *143* (13), 5022-5037.

20. Lincoln, R.; Bossi, M. L.; Remmel, M.; D'Este, E.; Butkevich, A. N.; Hell, S. W., A general design of caging-group free photoactivatable fluorophores for live-cell nanoscopy. *bioRxiv* **2021**, 2021.11.15.468659.

### NMR SPECTRA

#### Compound 1 $^{1}H$

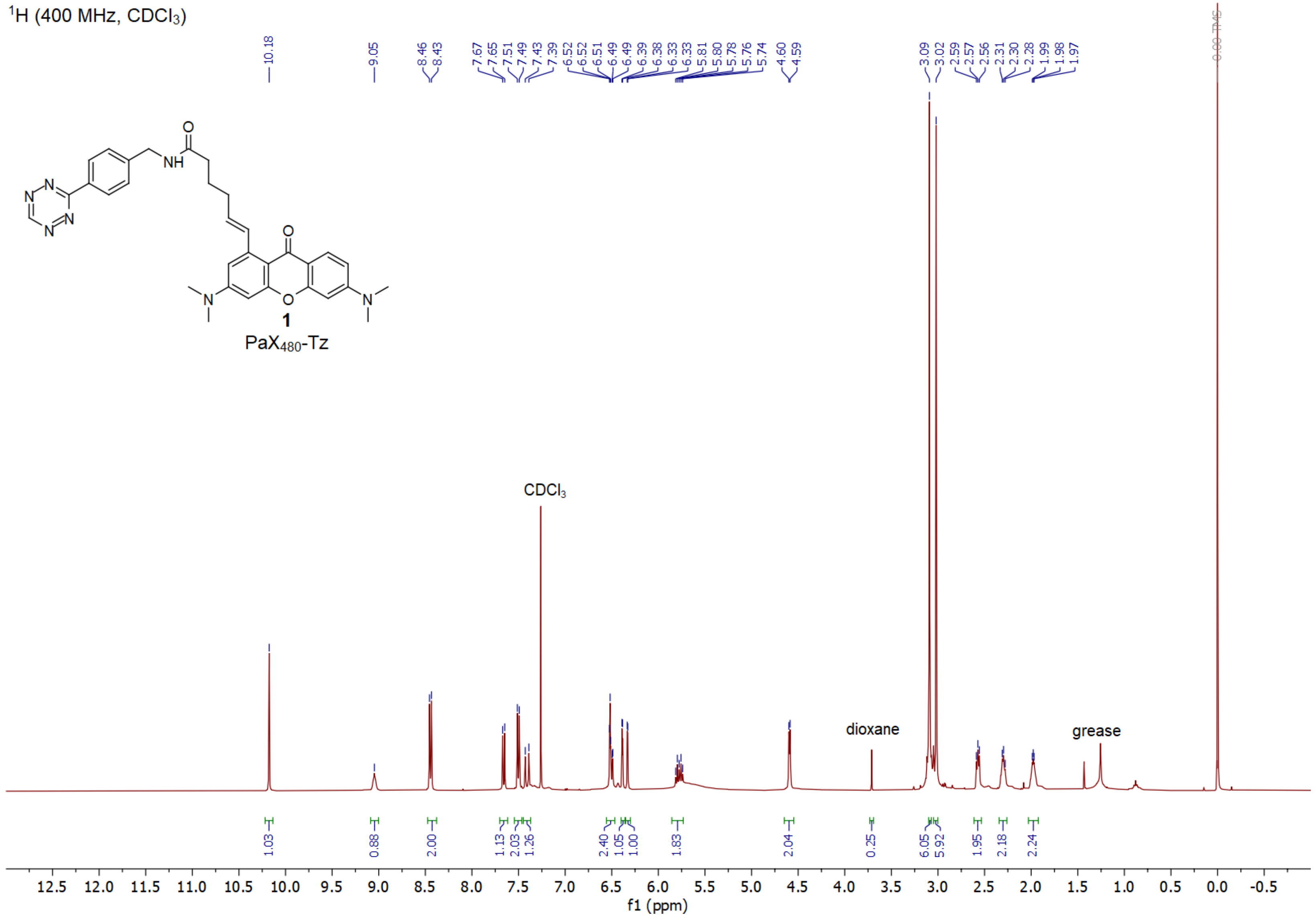

**Compound 1 $^{13}C$**

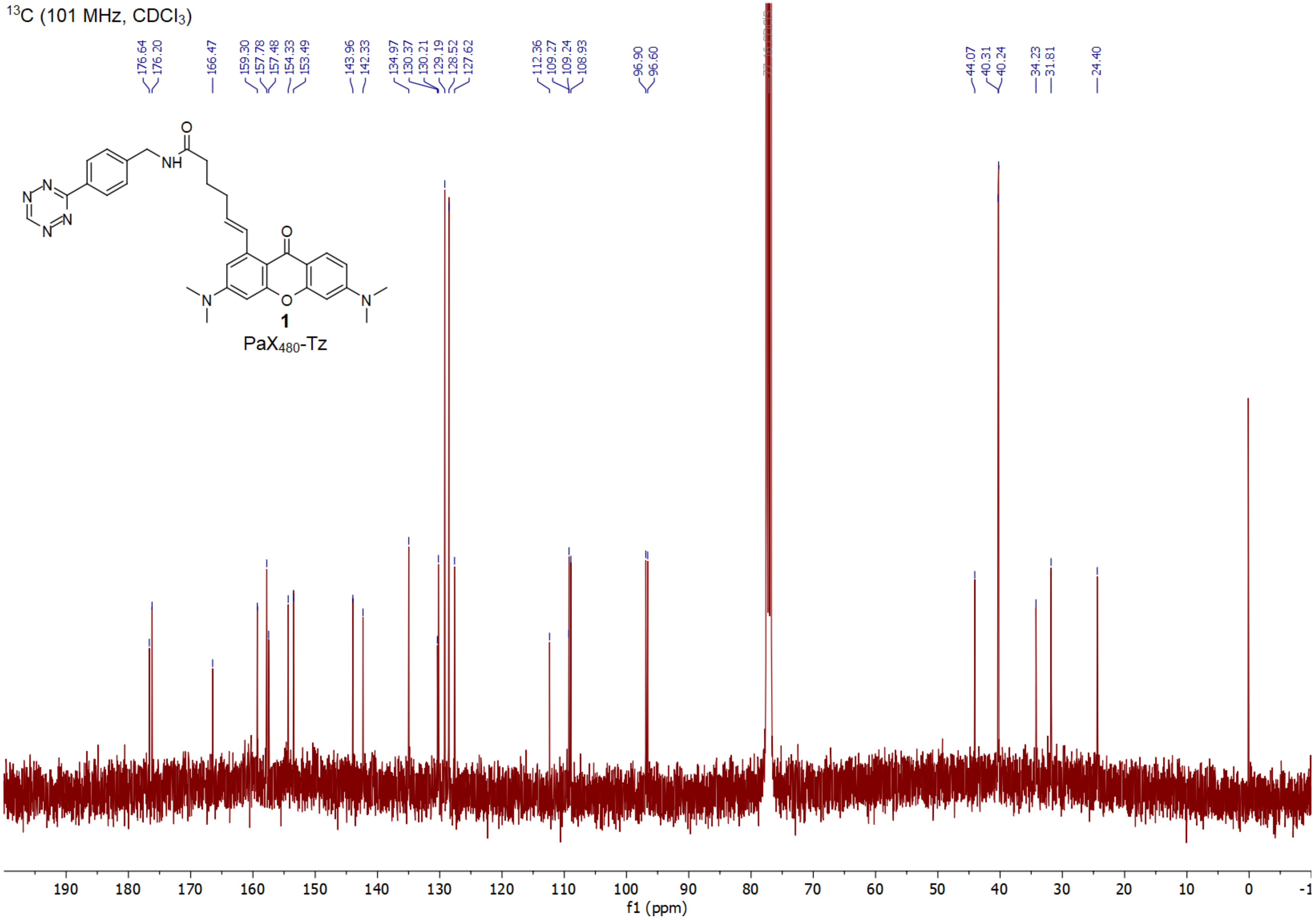

#### Compound 2 $^{1}H$

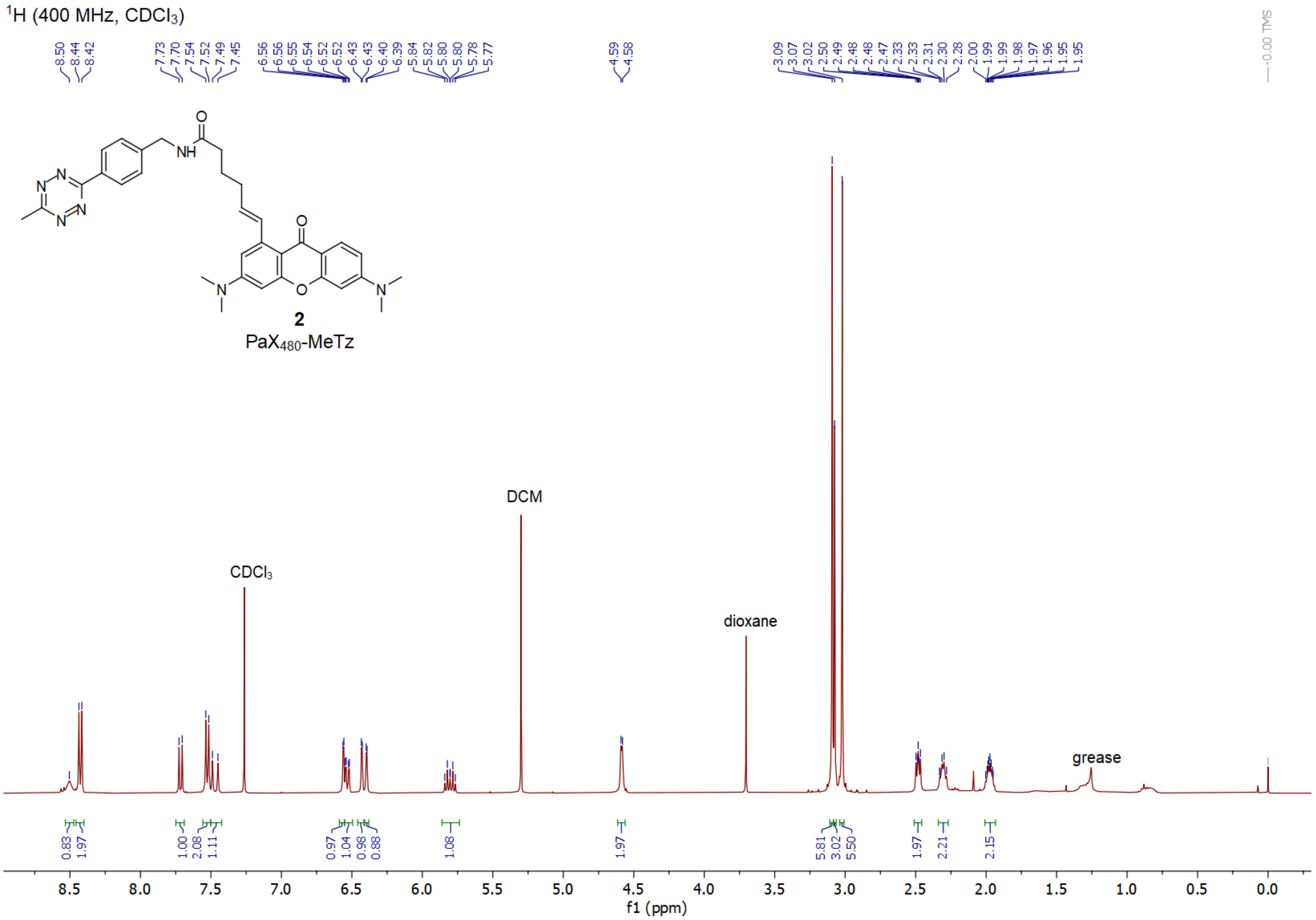

**Compound 2 $^{13}$C**

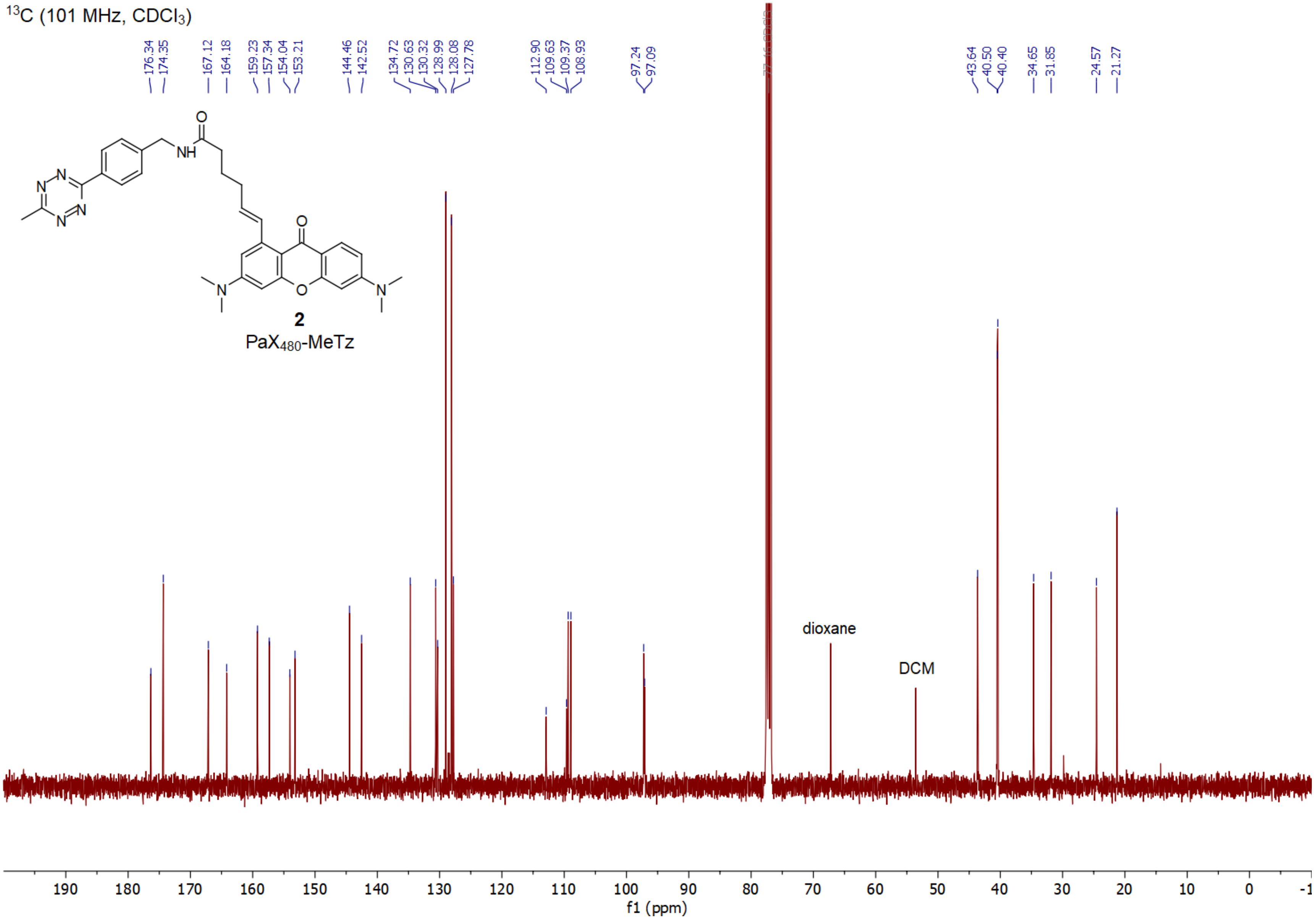

**Compound 3 $^{1}$H**

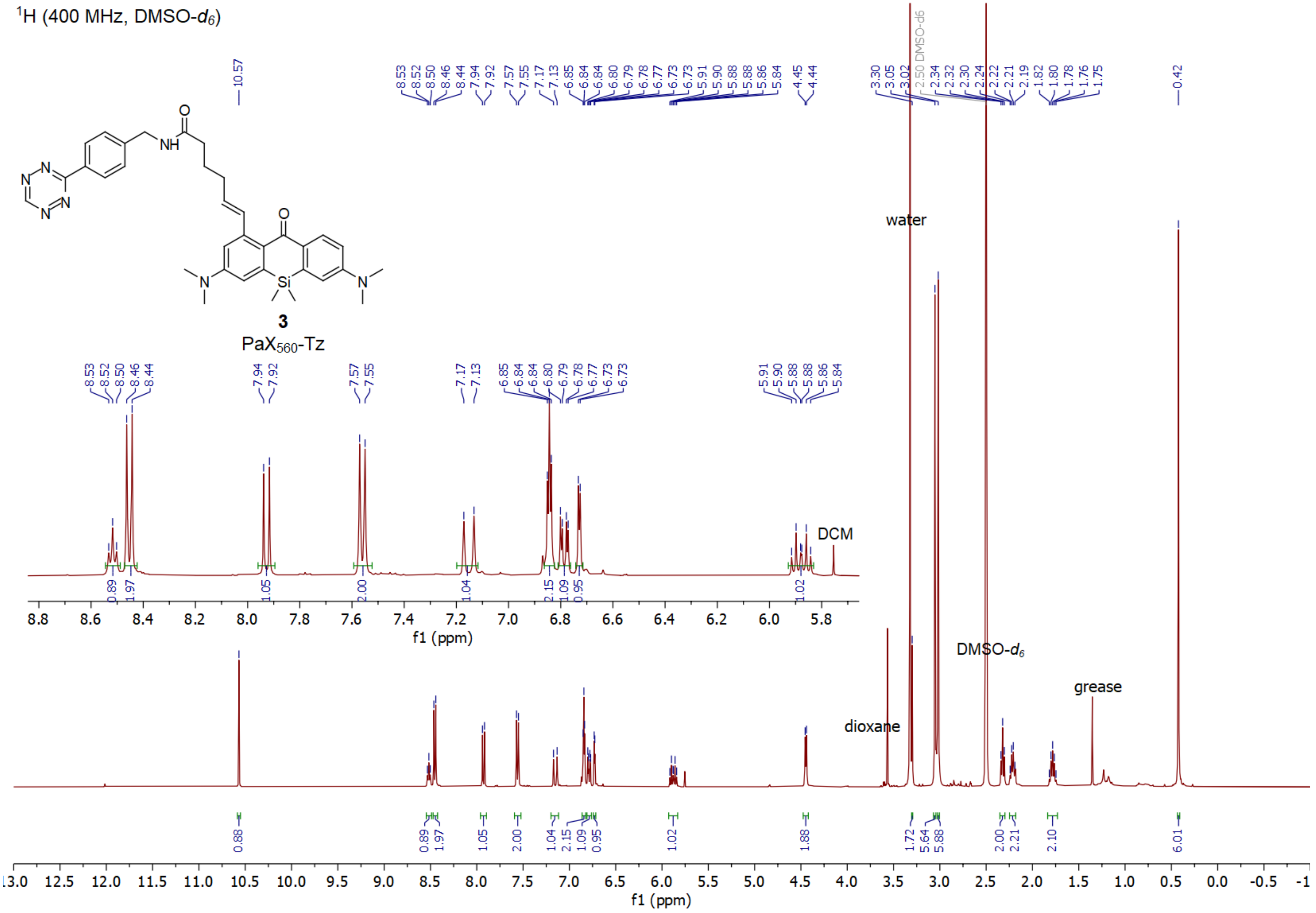

**Compound 3 $^{13}$C**

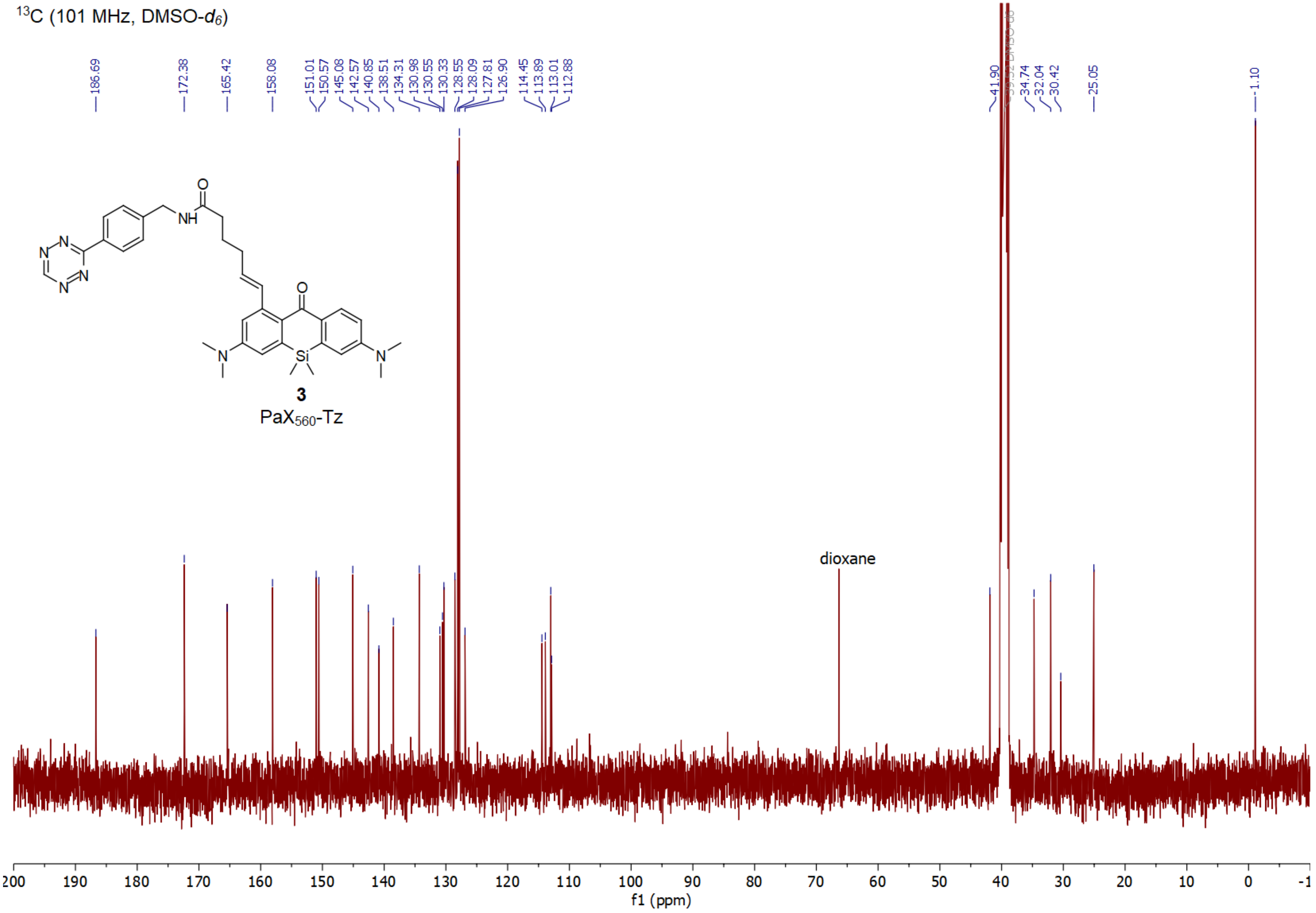

**Compound 4 $^{1}$H**

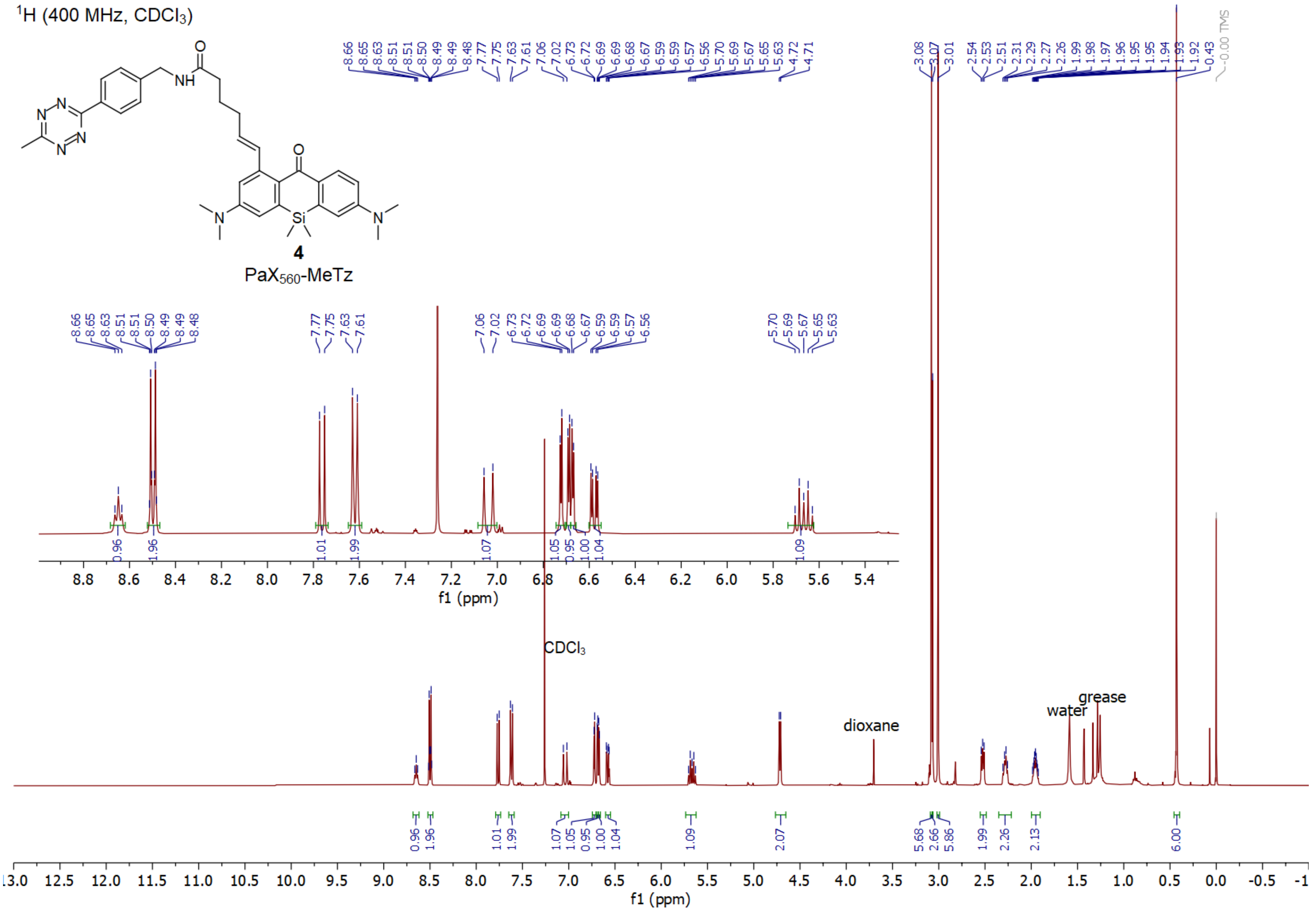

**Compound 4 $^{13}C$**

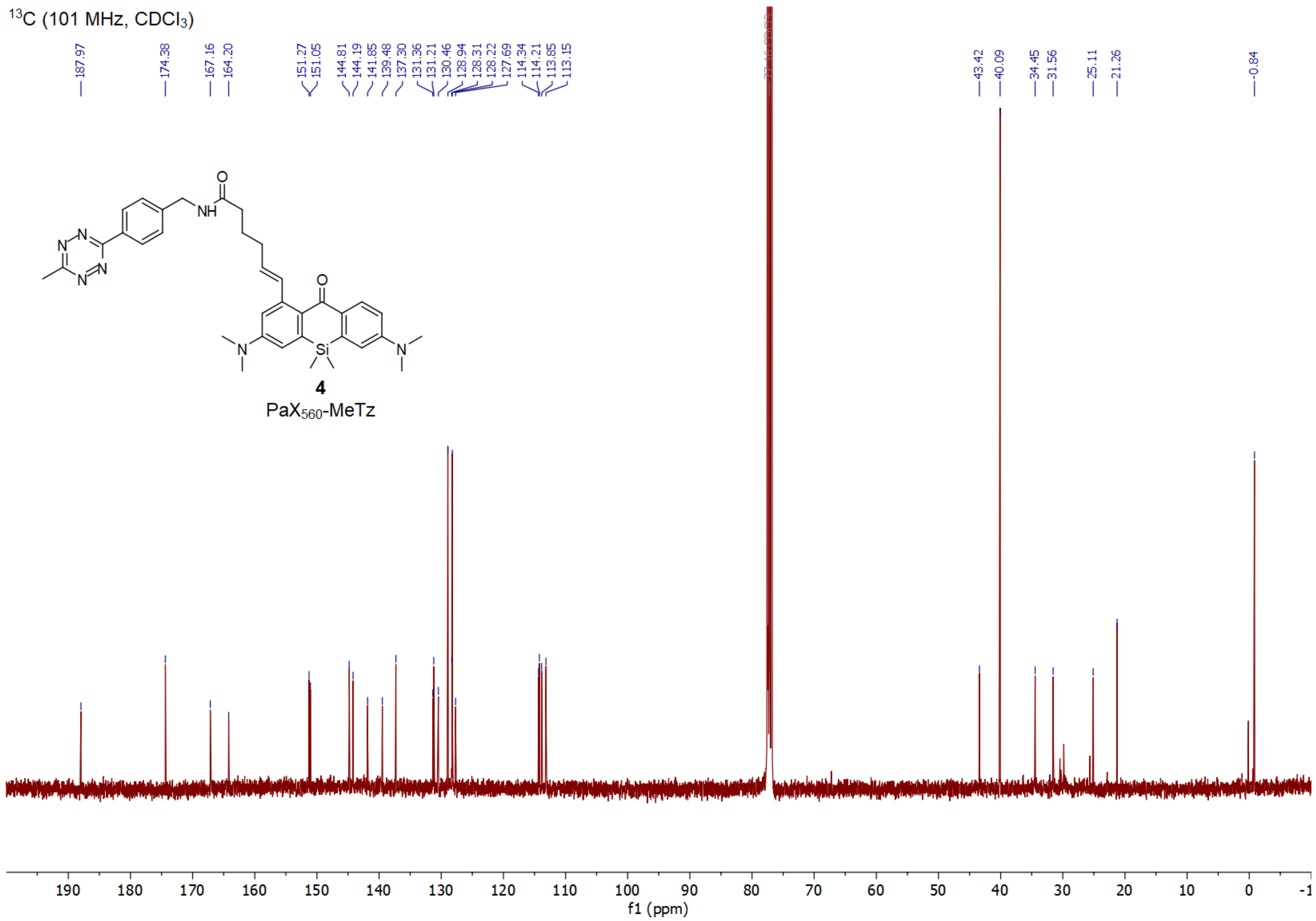

#### Compound 5 $^{1}H$

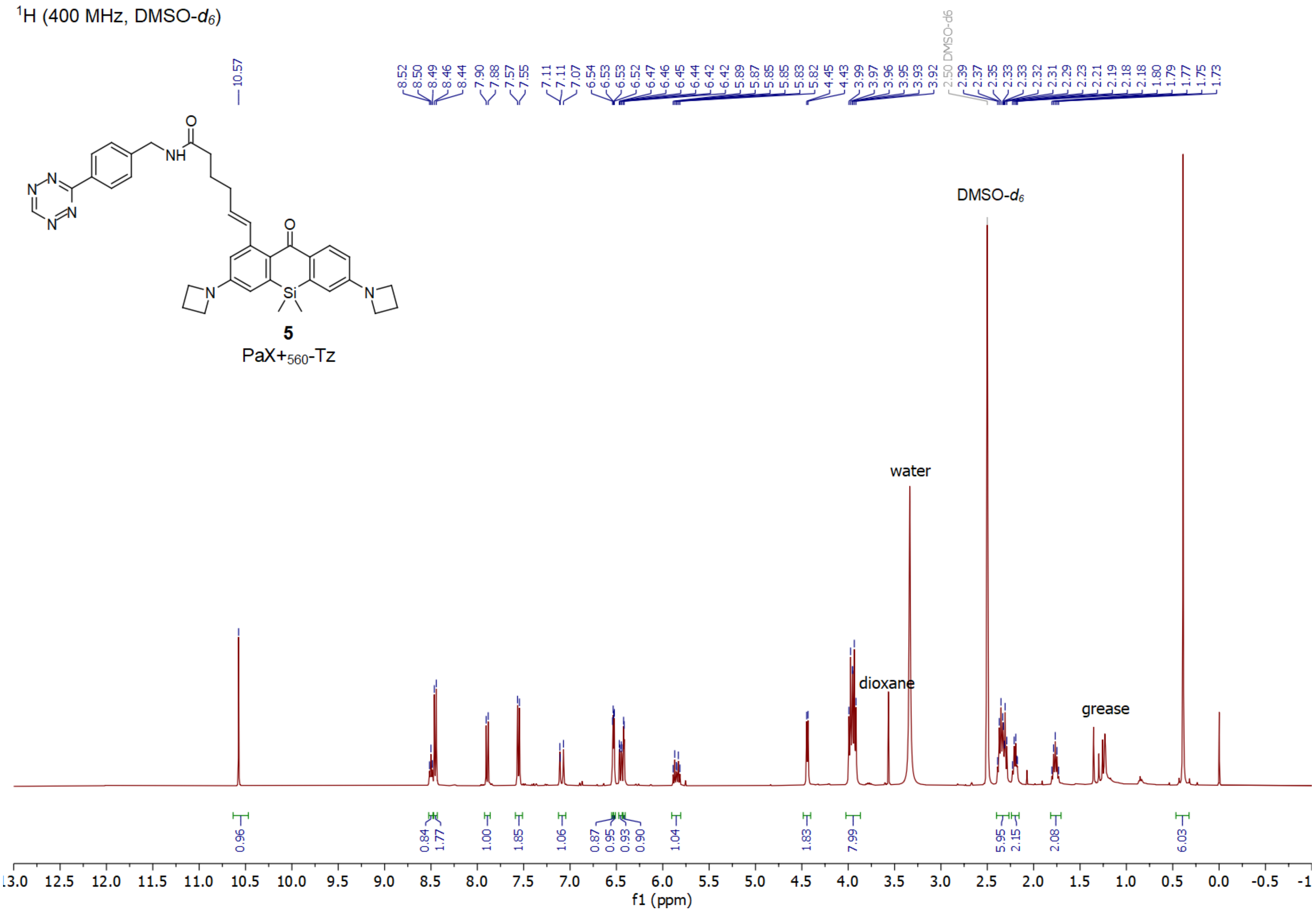

#### Compound 5 $^{13}C$

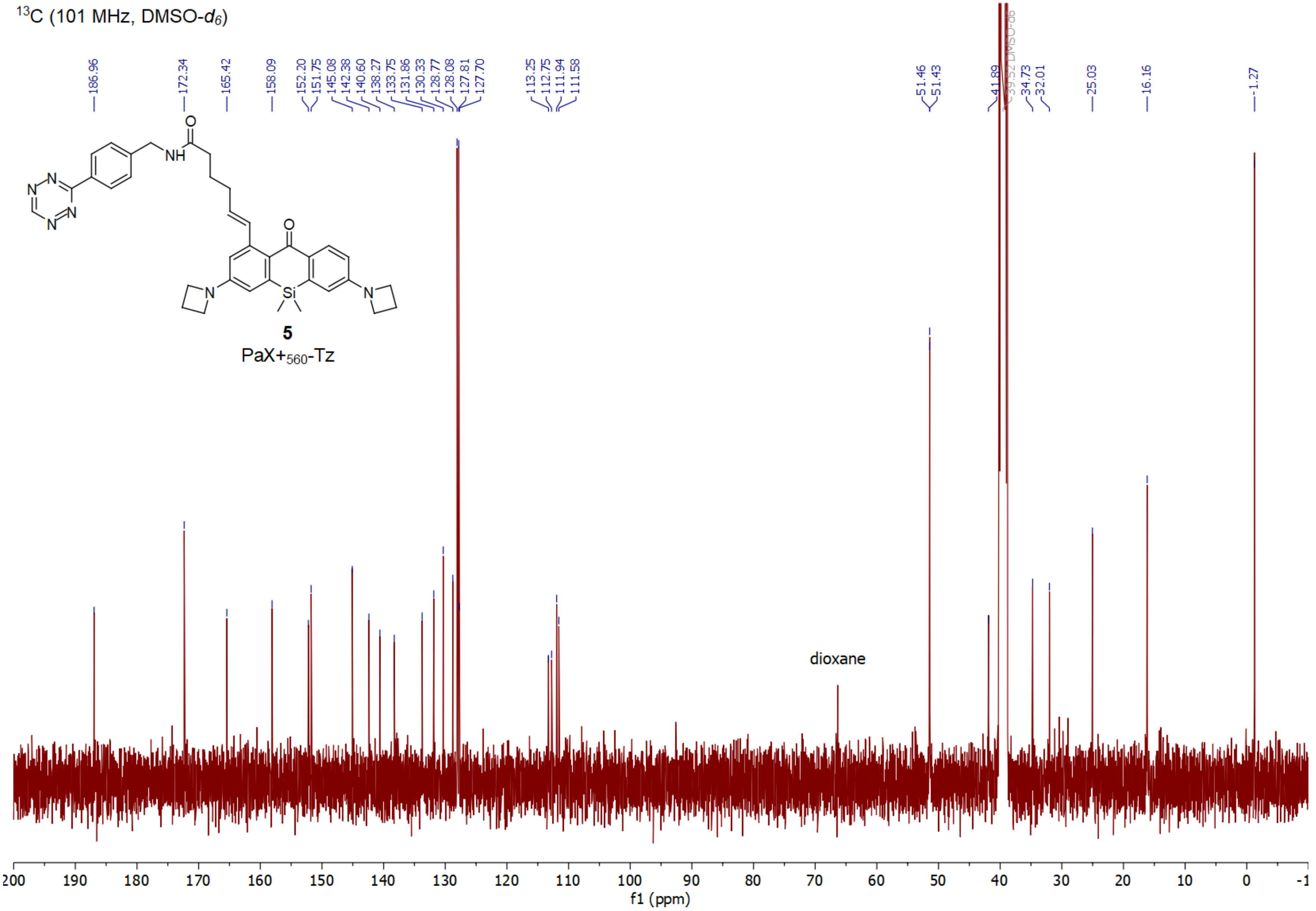

**Compound 6 $^{1}$H**

$^{1}$H (400 MHz, $CDCl_3$)

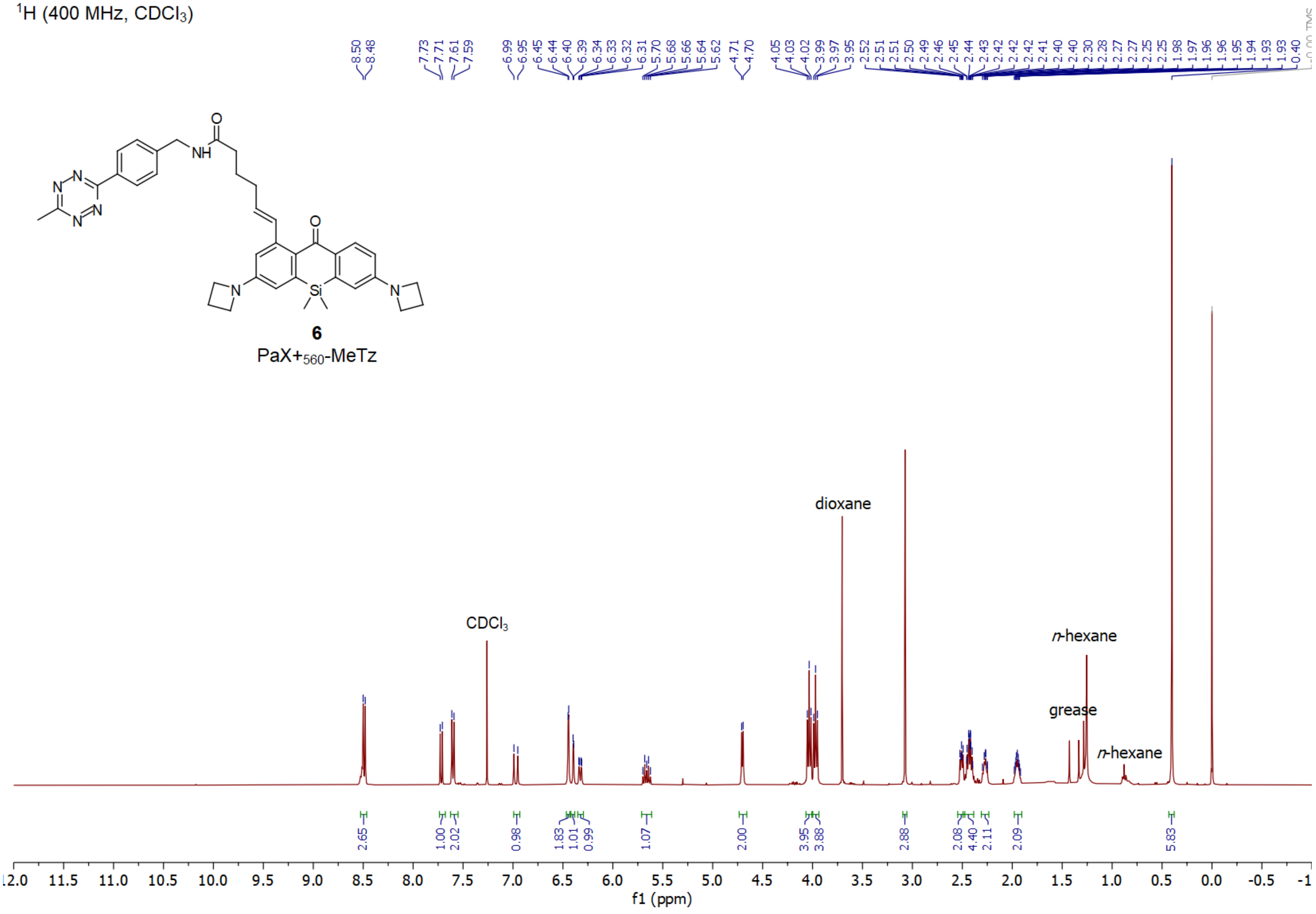

**Compound 6 $^{13}$C**

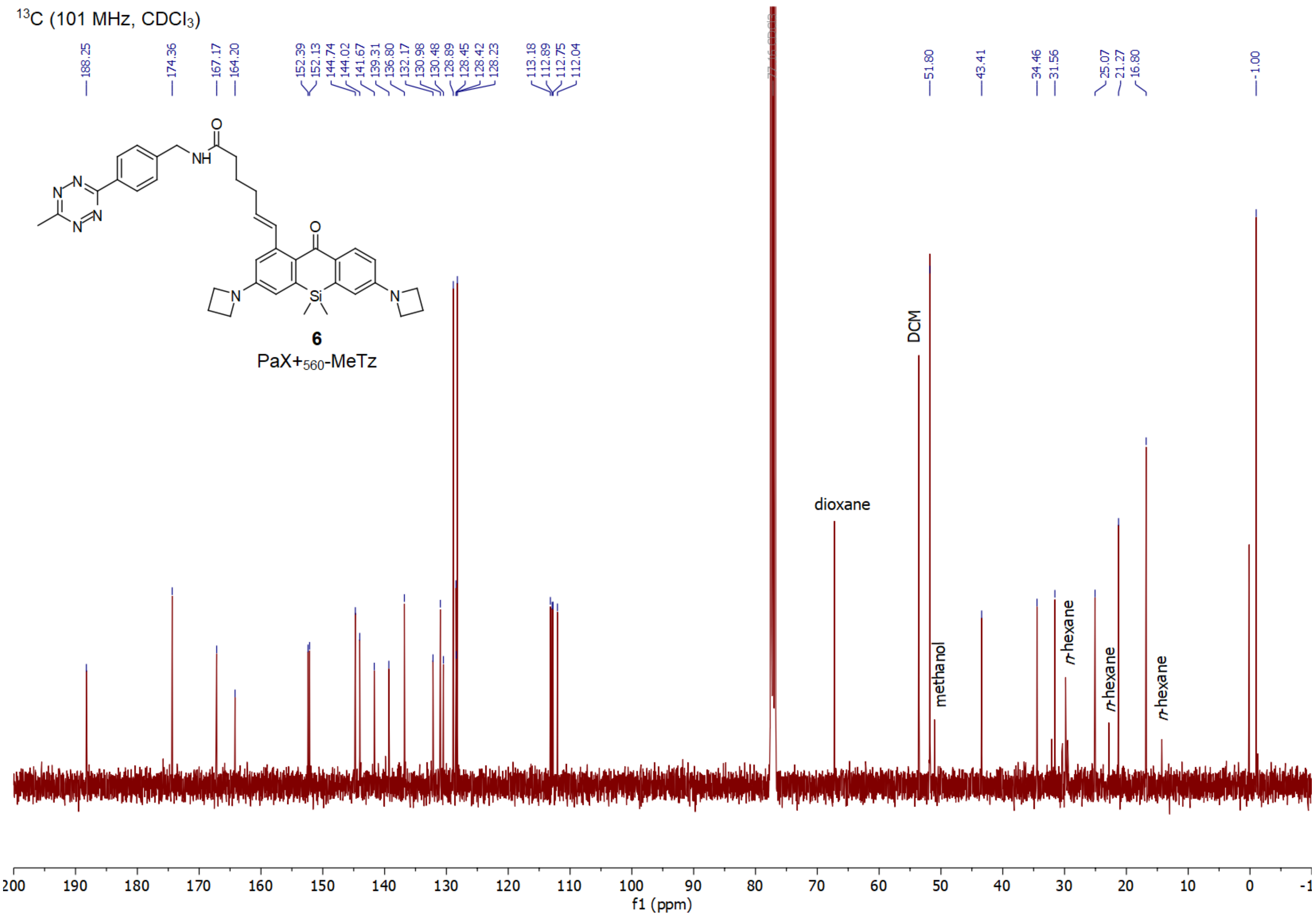

**Compound S6 $^{1}$H**

$^{1}$H (400 MHz, $CDCl_3$)

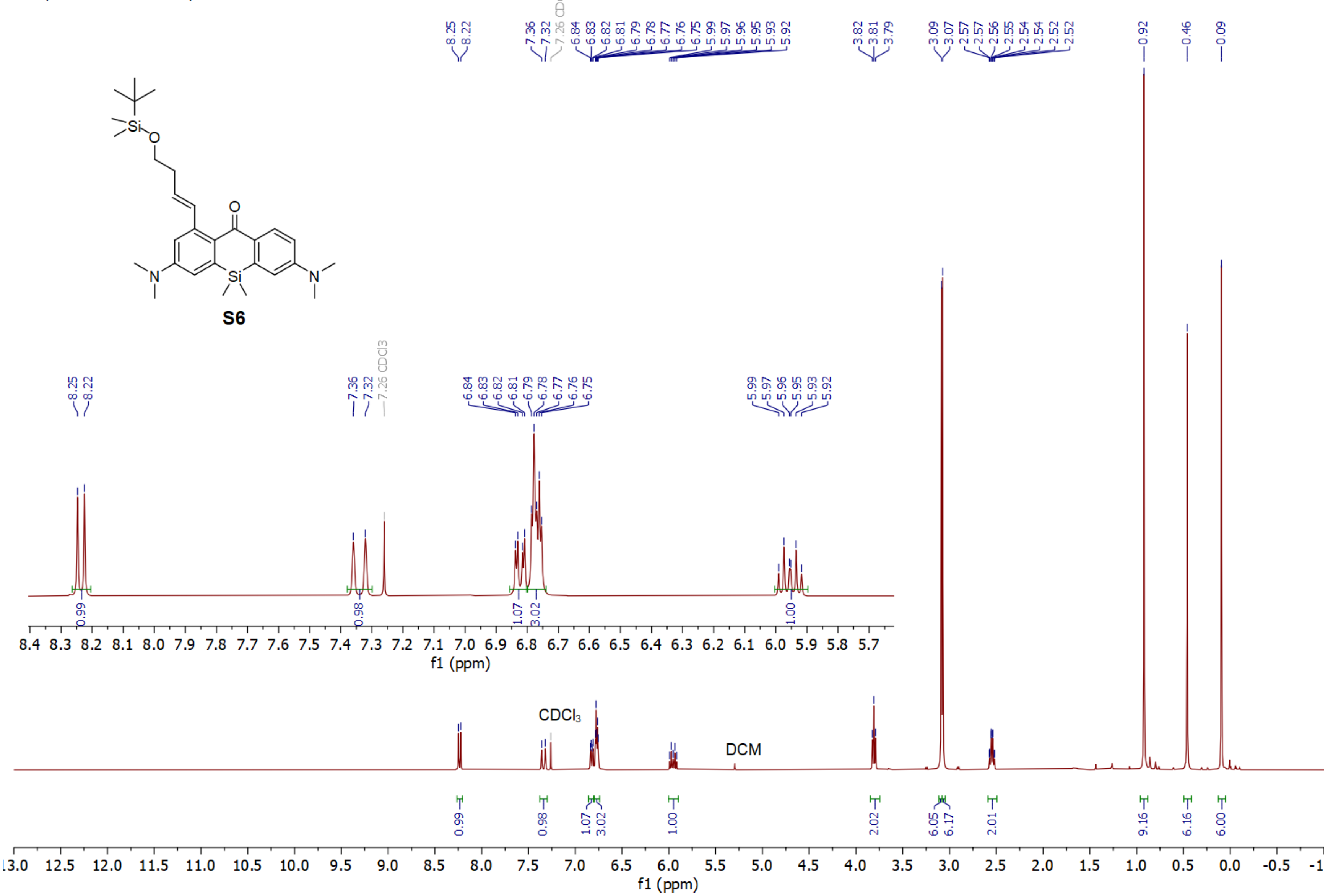

**Compound S6 $^{13}C$**

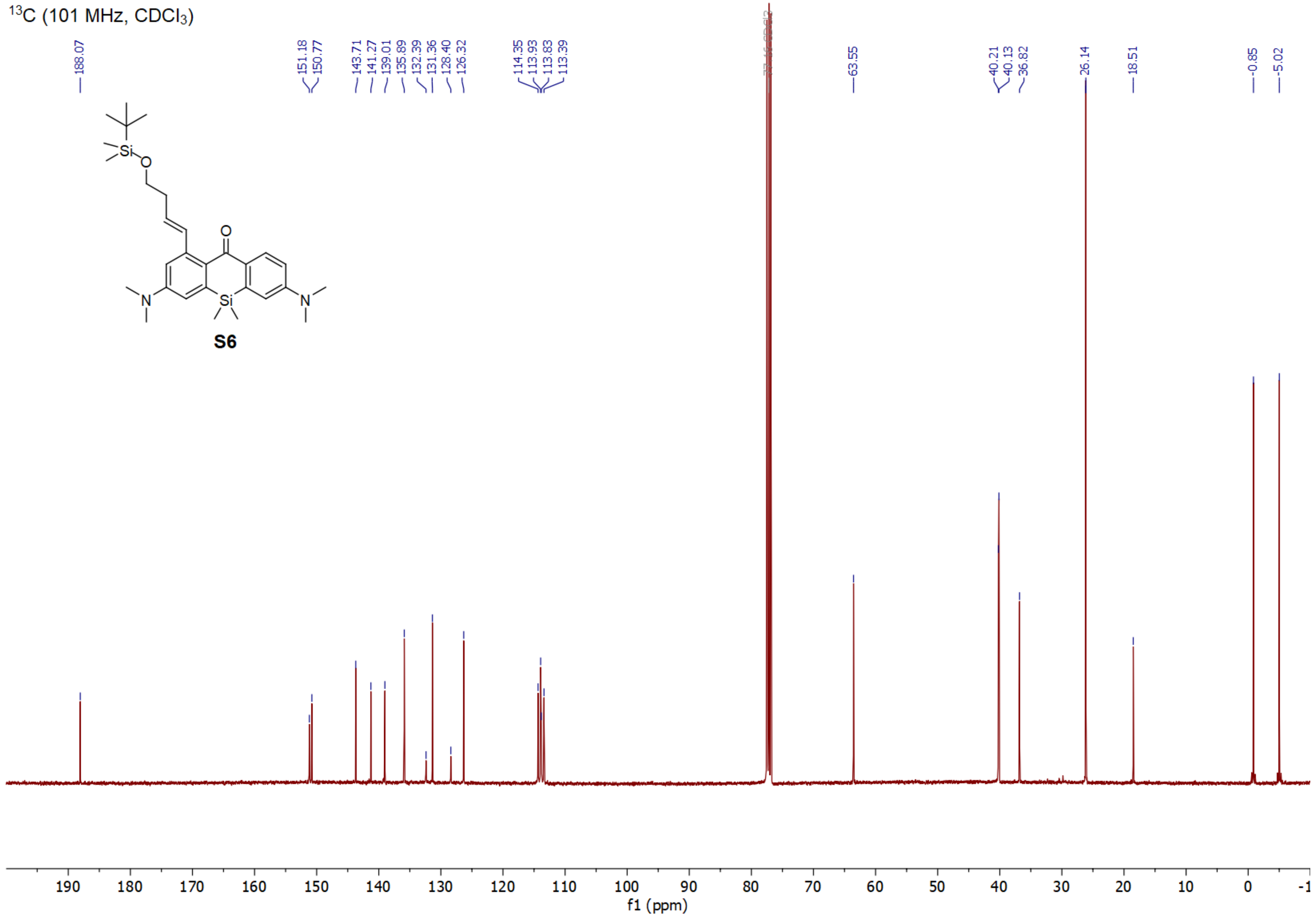

**Compound S7 $^{1}H$**

$^{1}H$ (400 MHz, $CDCl_3$)

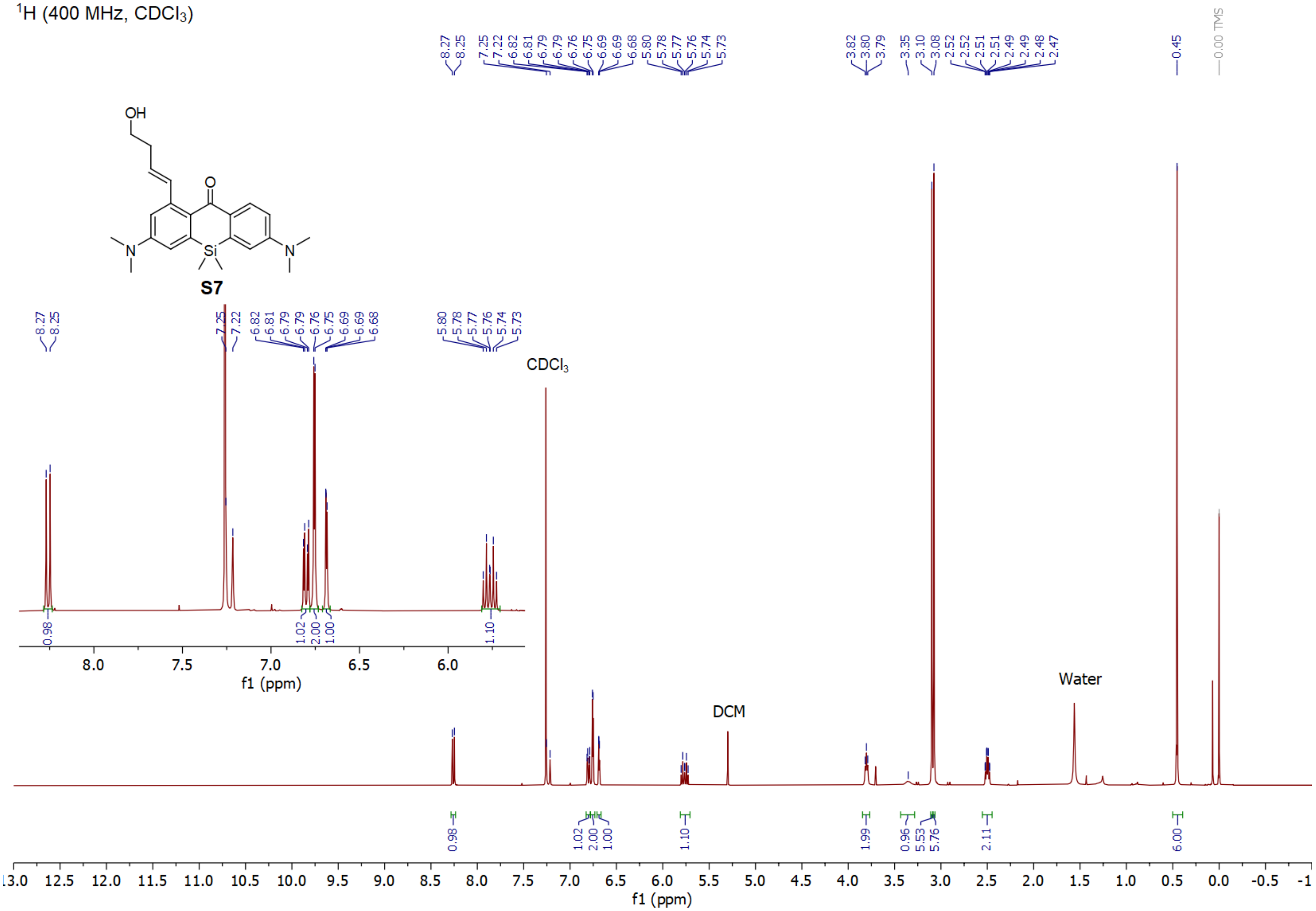

#### Compound S7 $^{13}C$

$^{13}C$ (101 MHz, $CDCl_3$)

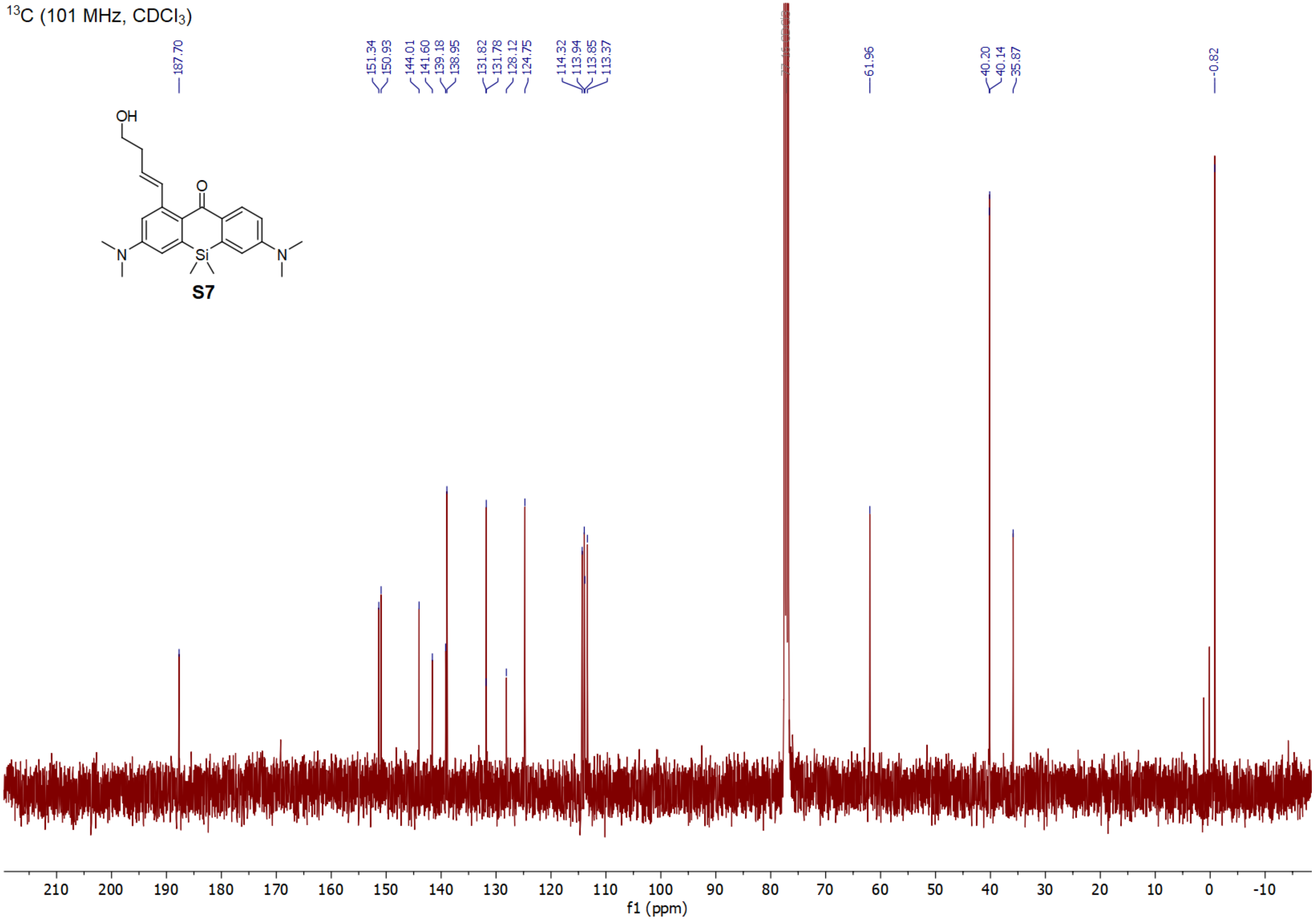

#### Compound S8 ¹H

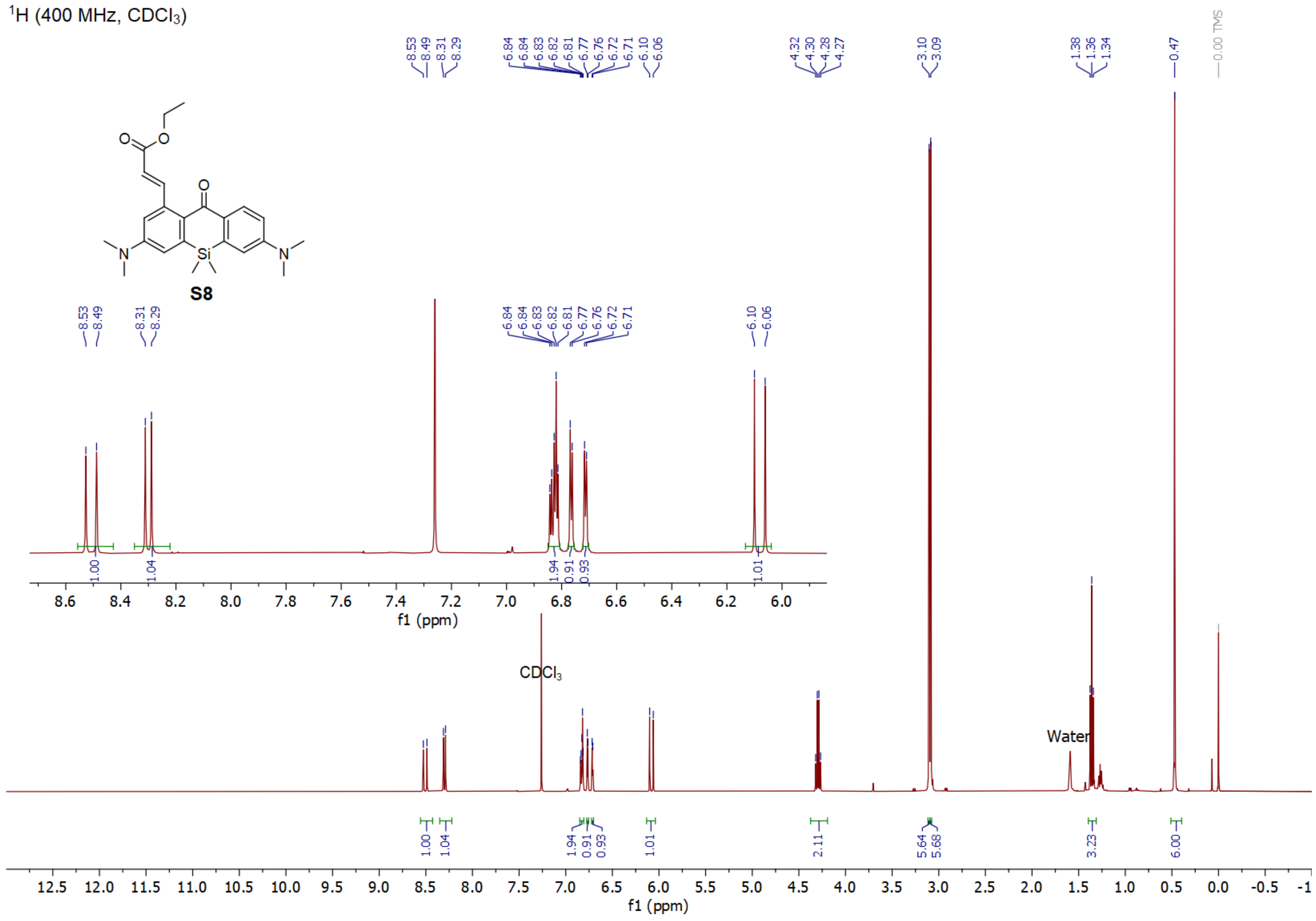

**Compound S8 $^{13}C$**

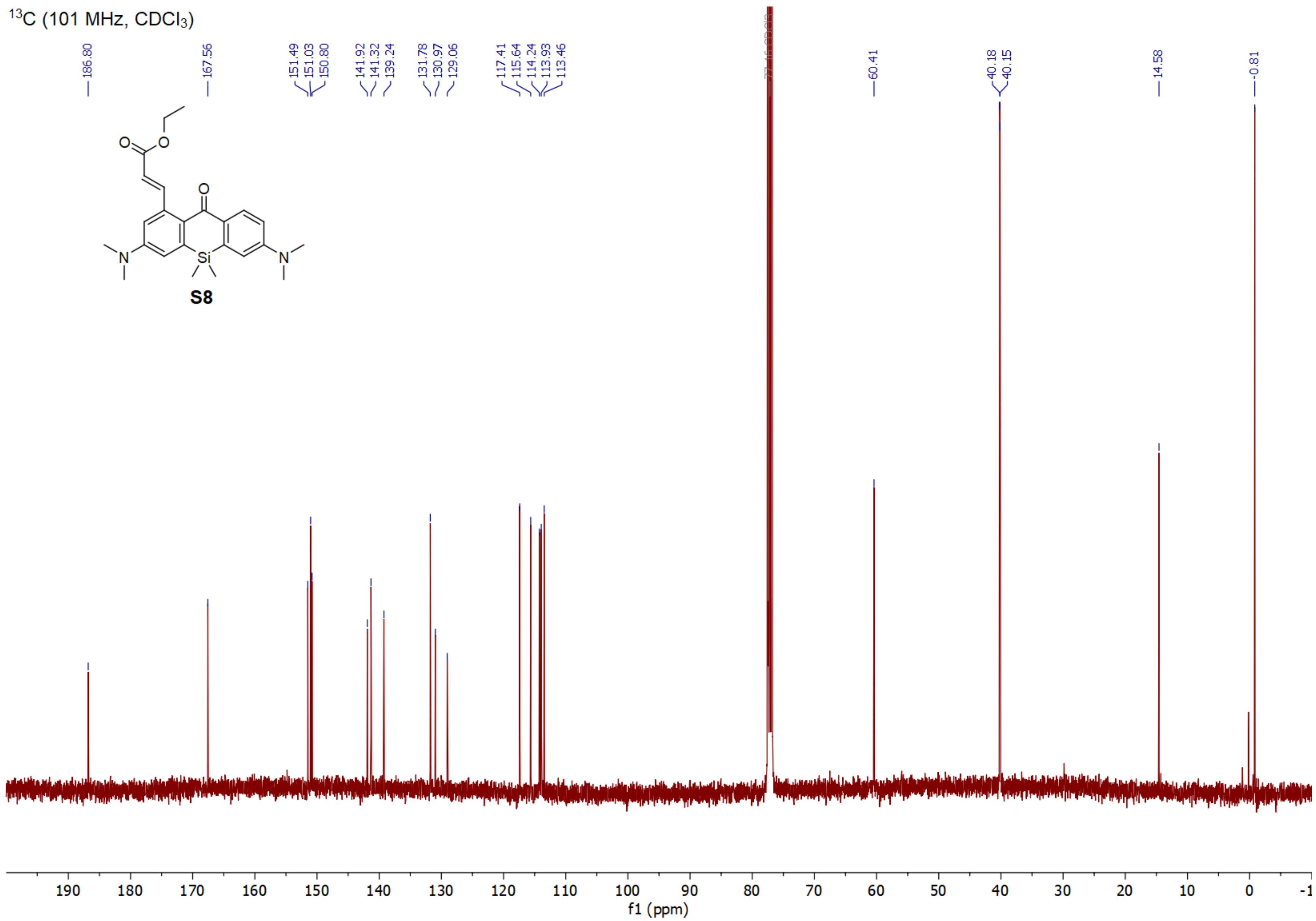

#### Compound S9 $^1$H

$^1$H (400 MHz, DMSO-$d_6$)

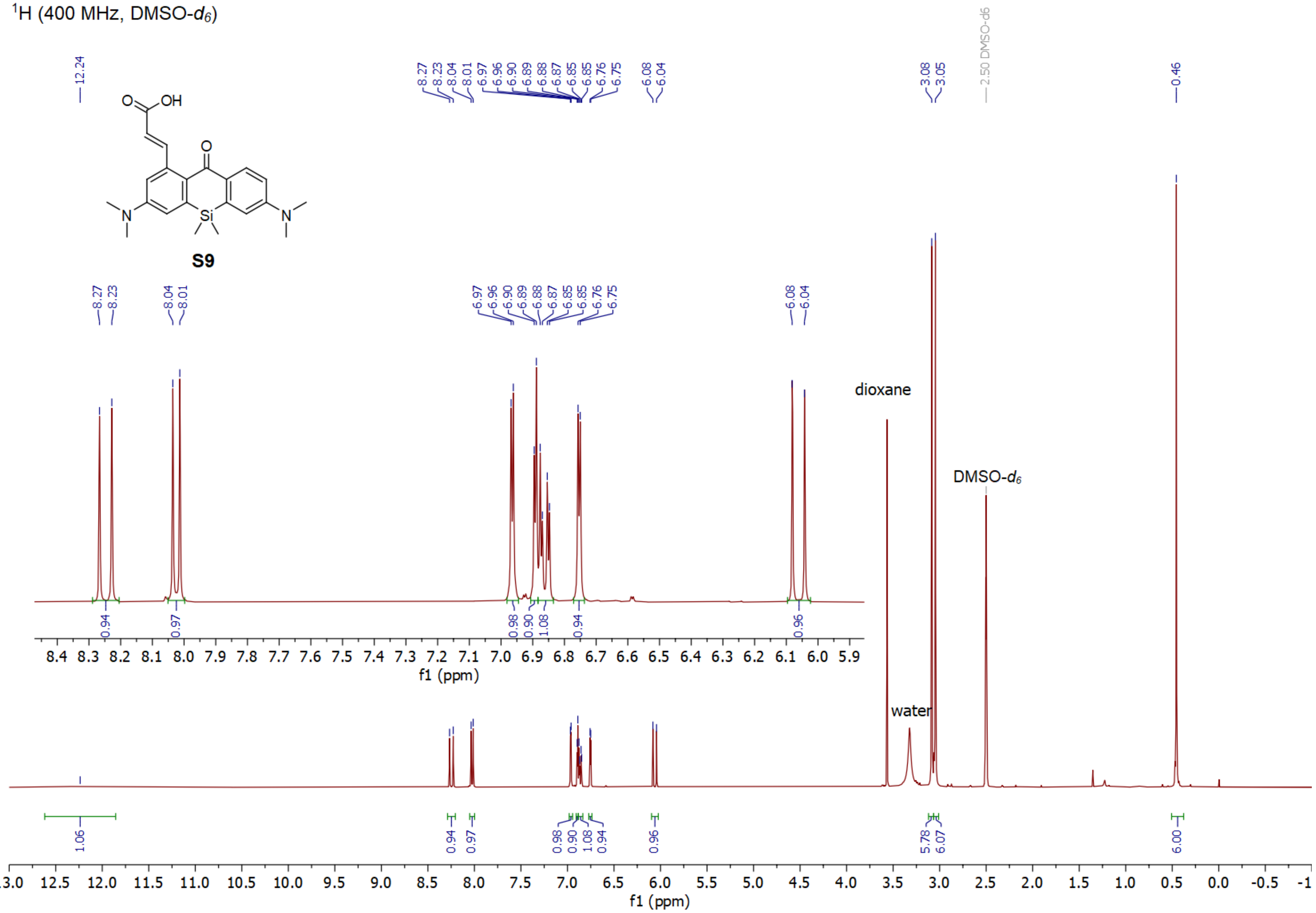

#### Compound S9 $^{13}C$

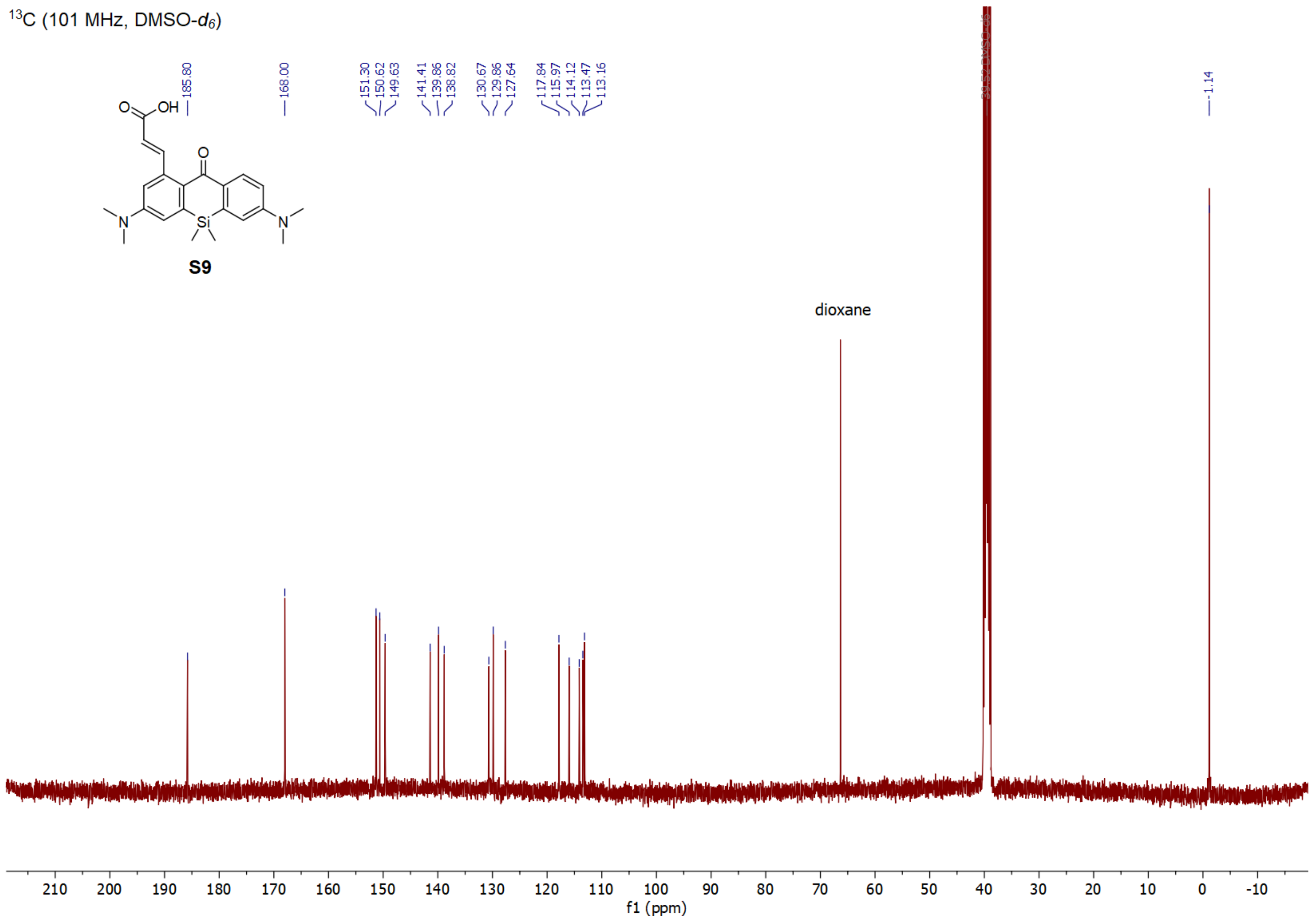

#### Compound 10 $^{1}$H

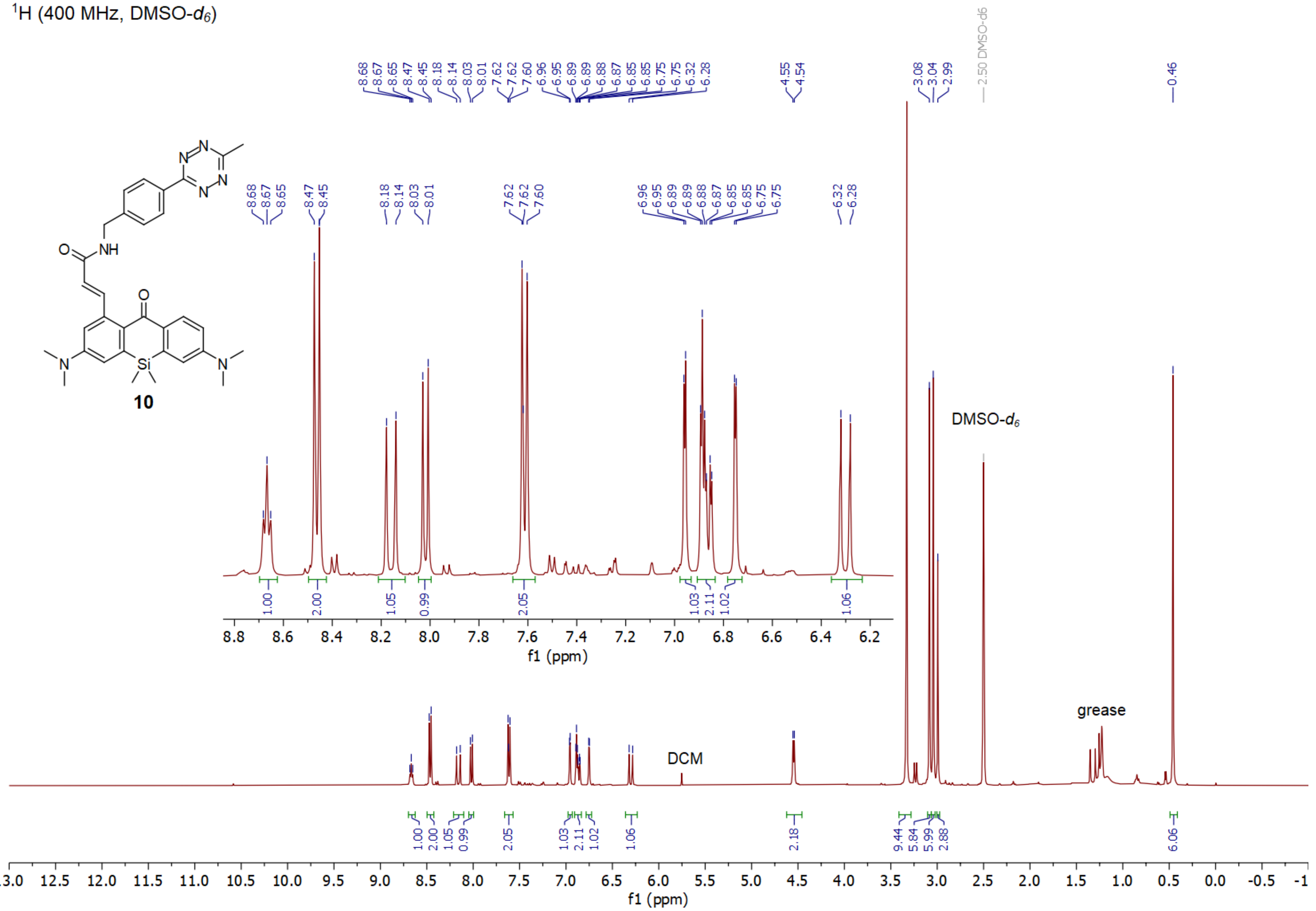

#### Compound 10 $^{13}C$

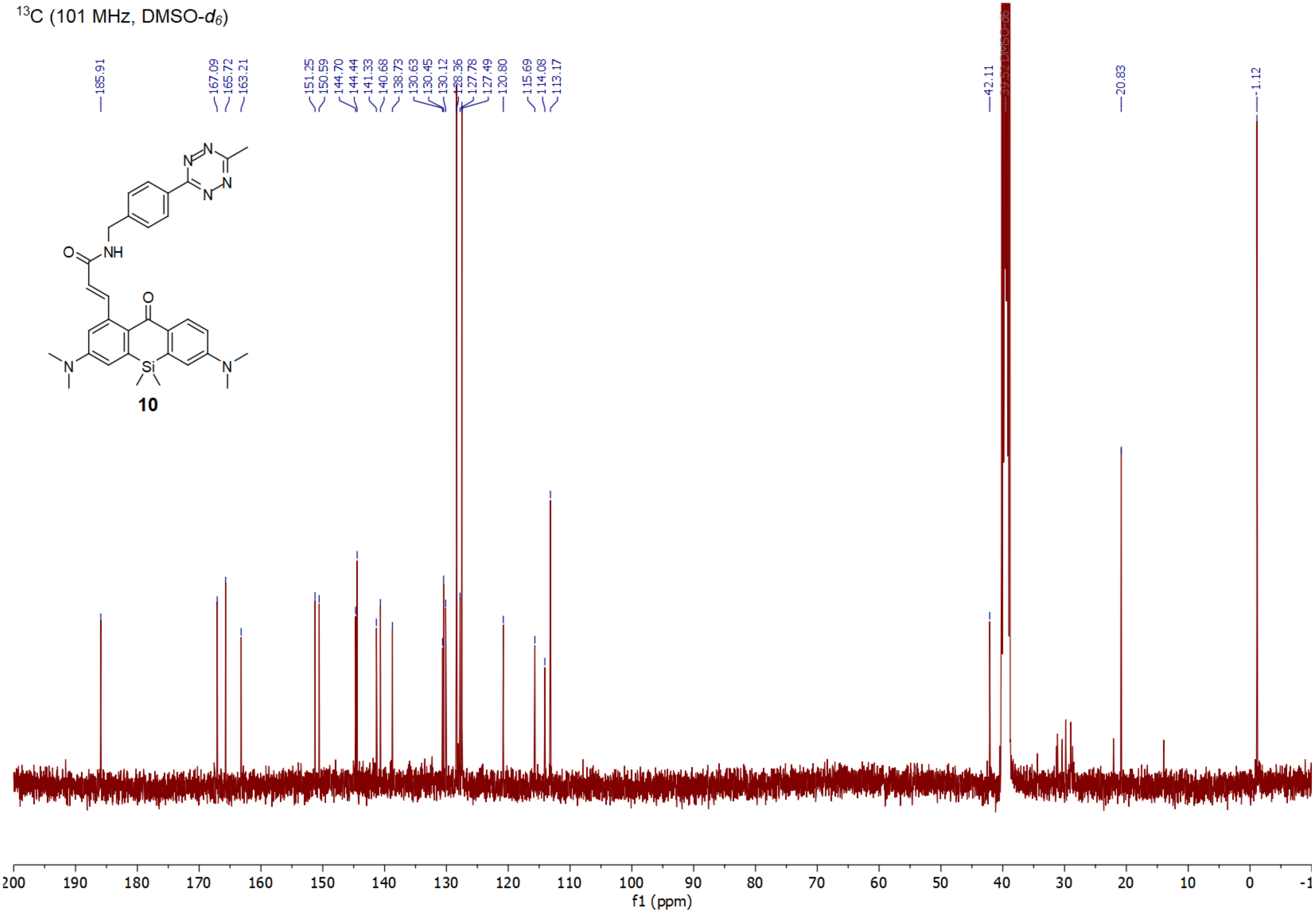

**Compound S10 $^{1}$H**

$^{1}$H (400 MHz, DMSO-$d_6$)

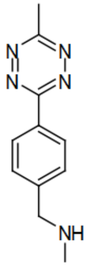

**S10**

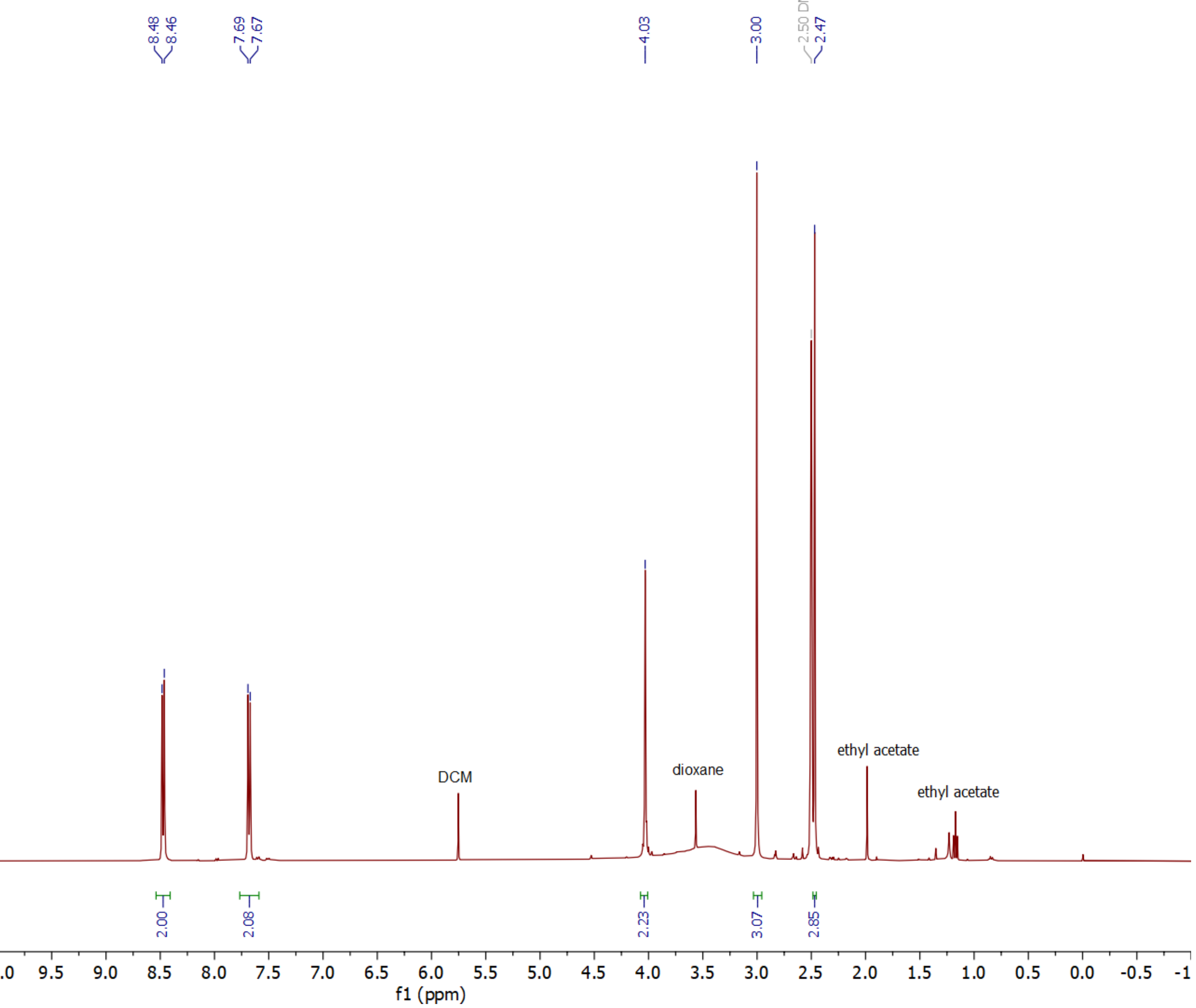

#### Compound S10 $^{13}C$

$^{13}C$ (101 MHz, DMSO-$d_6$)

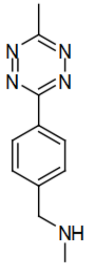

**S10**

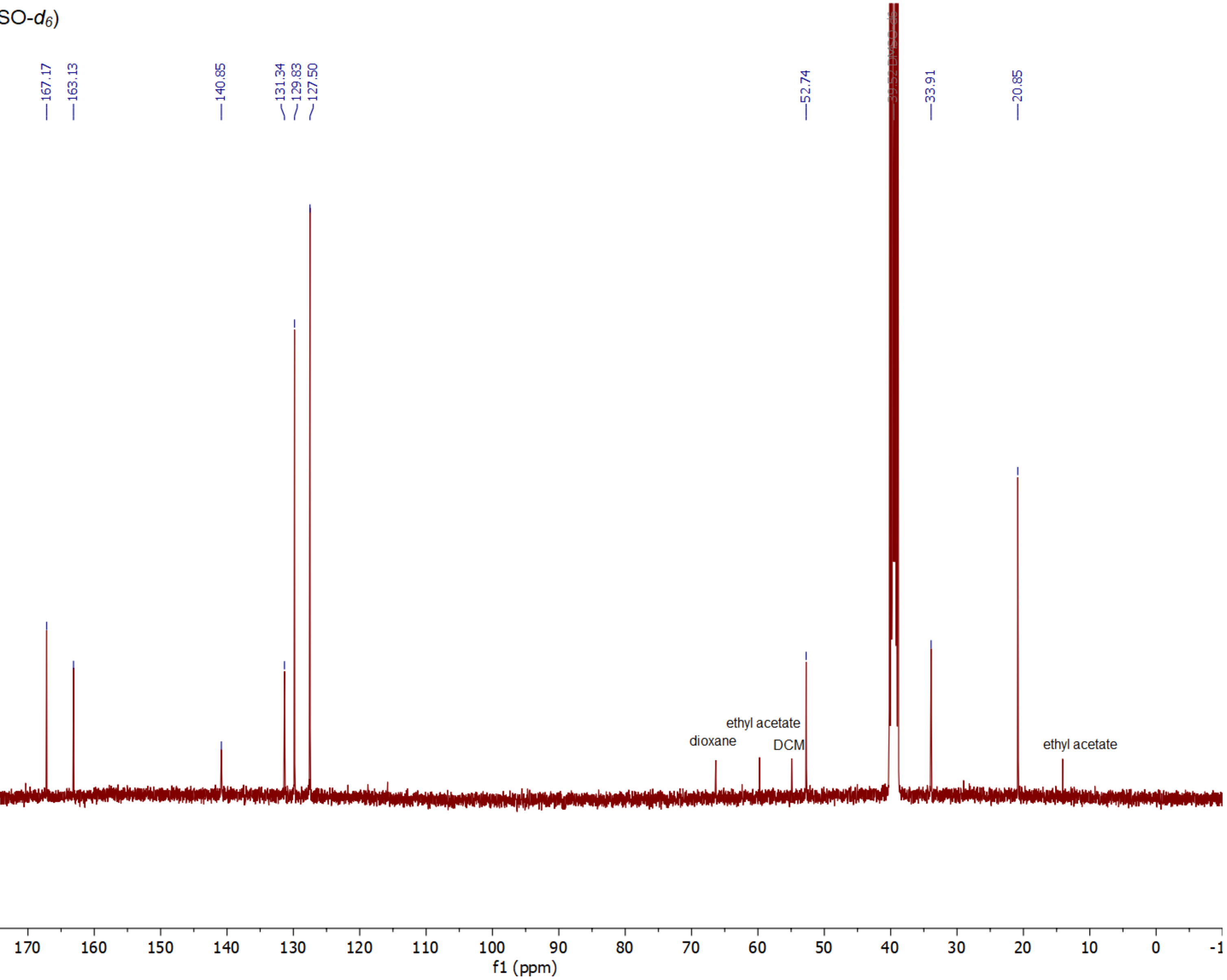

**Compound S13 $^{1}H$**

$^{1}H$ (400 MHz, $CDCl_3$)

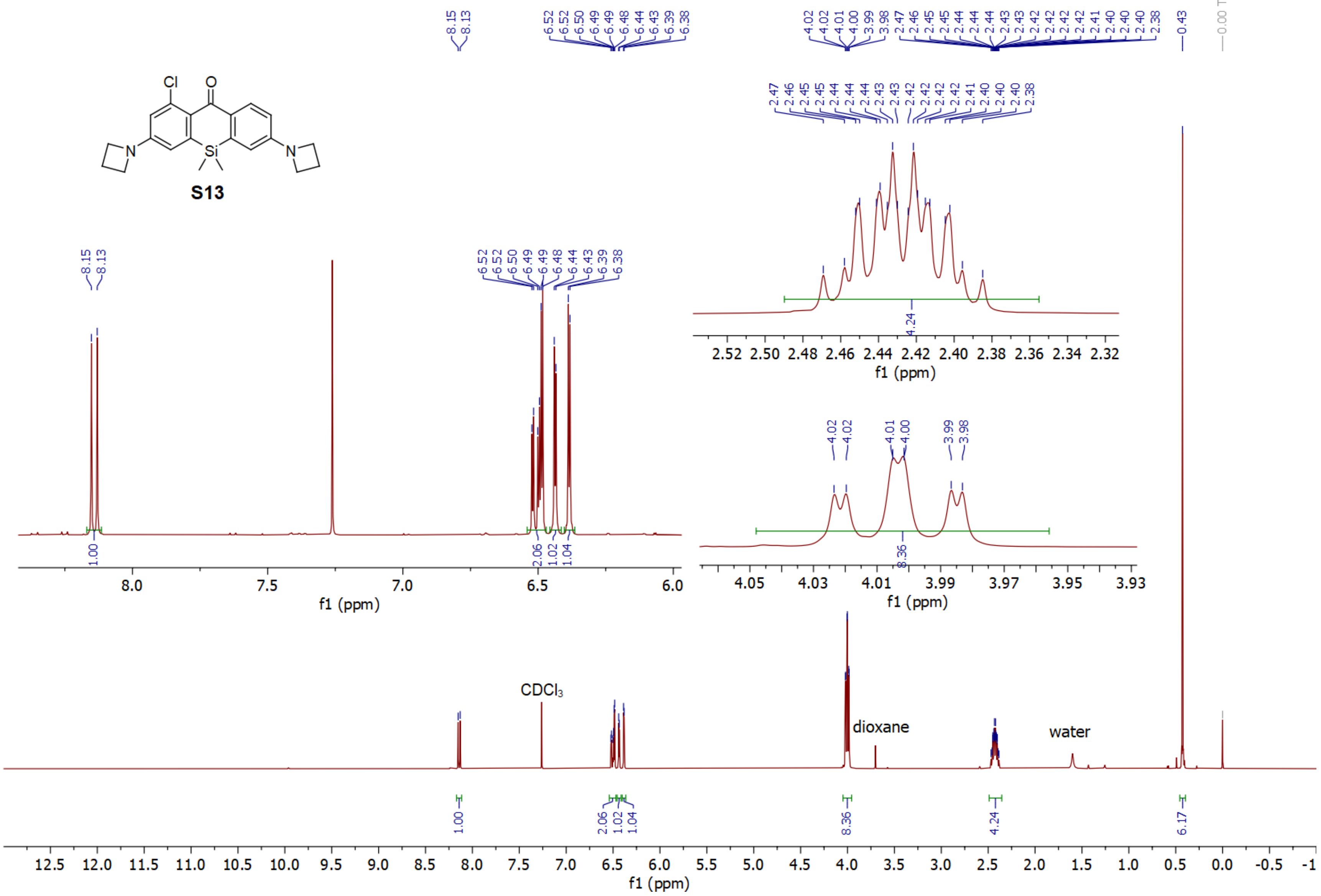

#### Compound S13 $^{13}C$

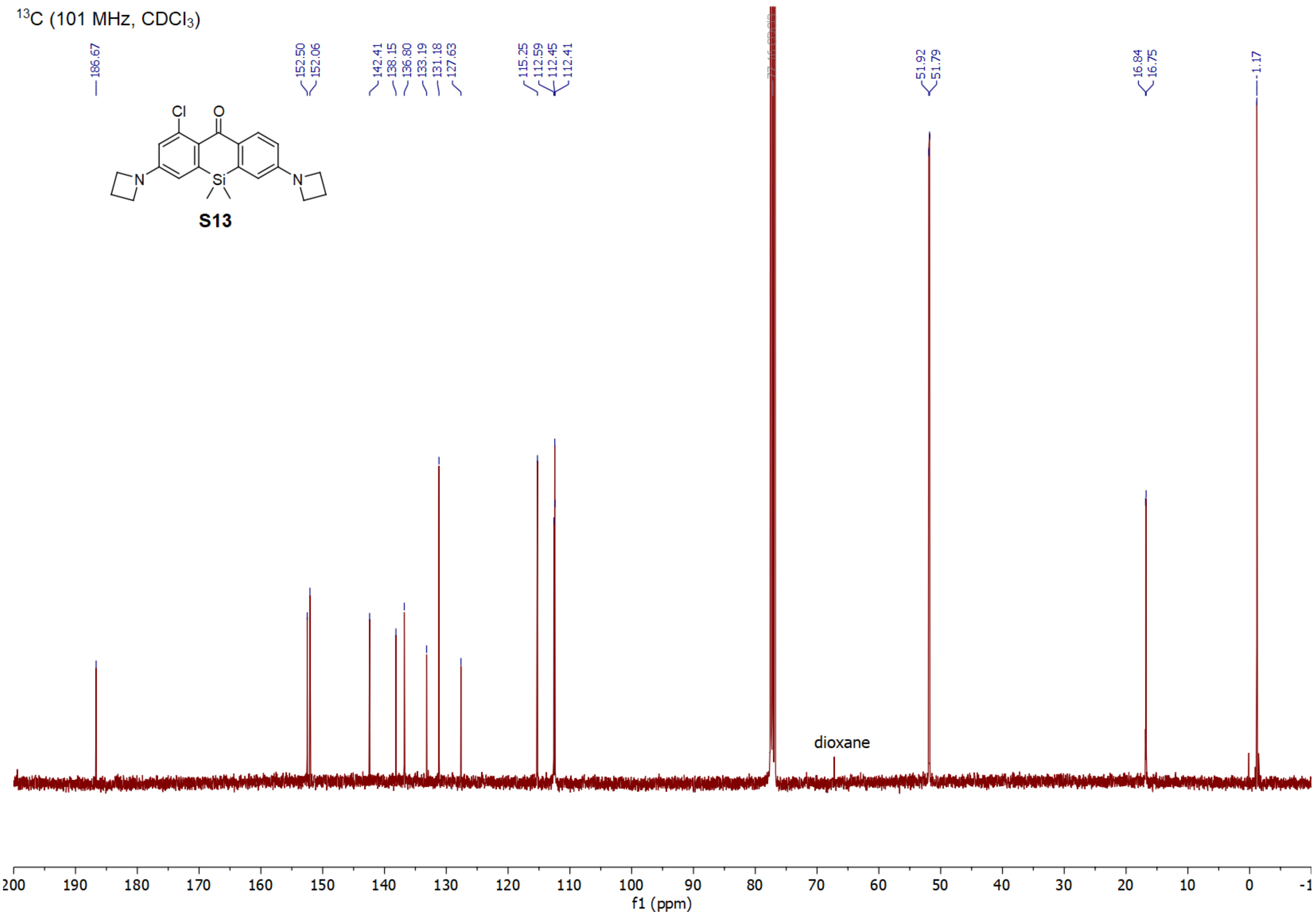

**Compound S14 $^{1}H$**

$^{1}H$ (400 MHz, DMSO-$d_6$)

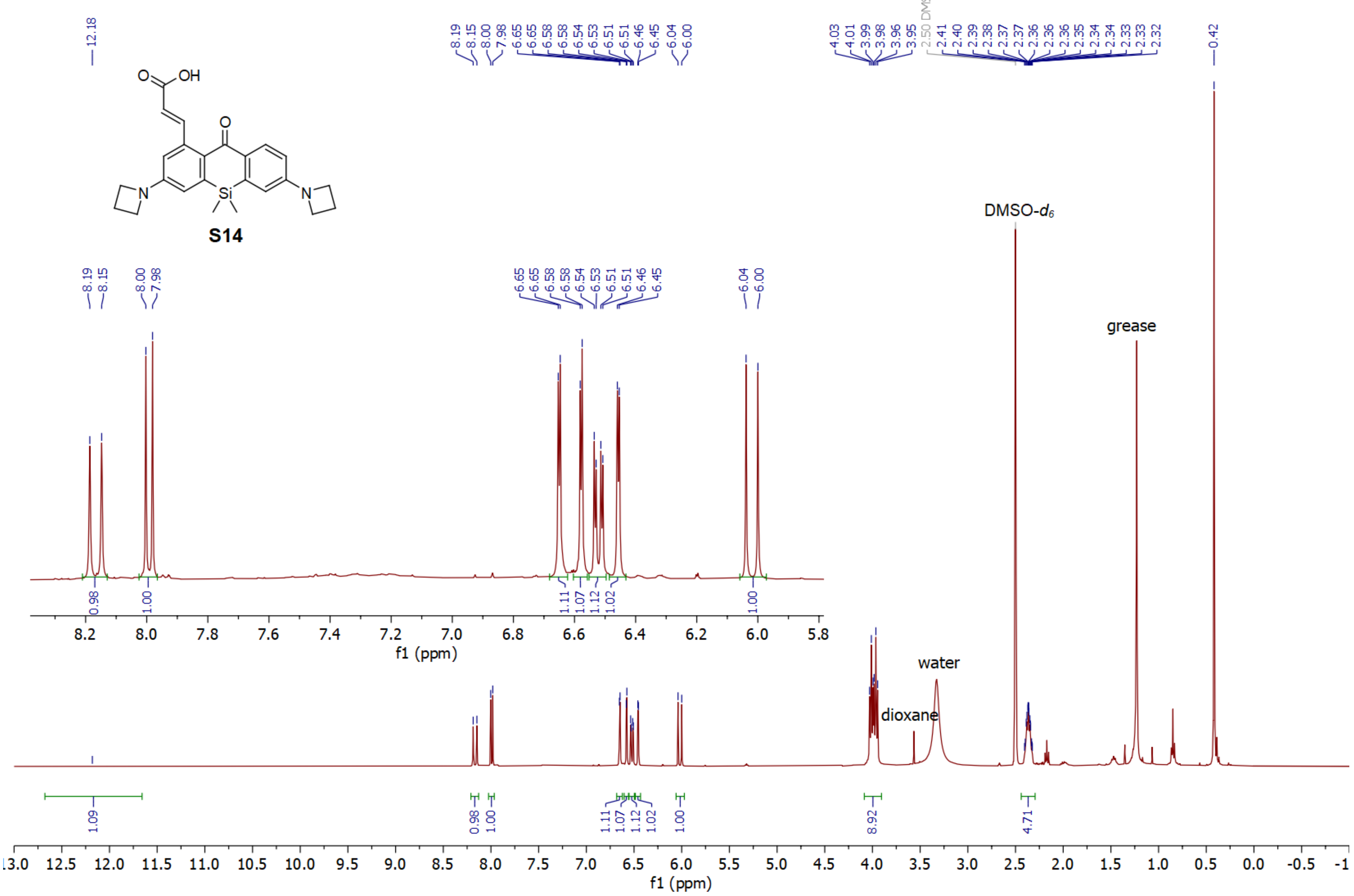

**Compound S14 $^{13}$C**

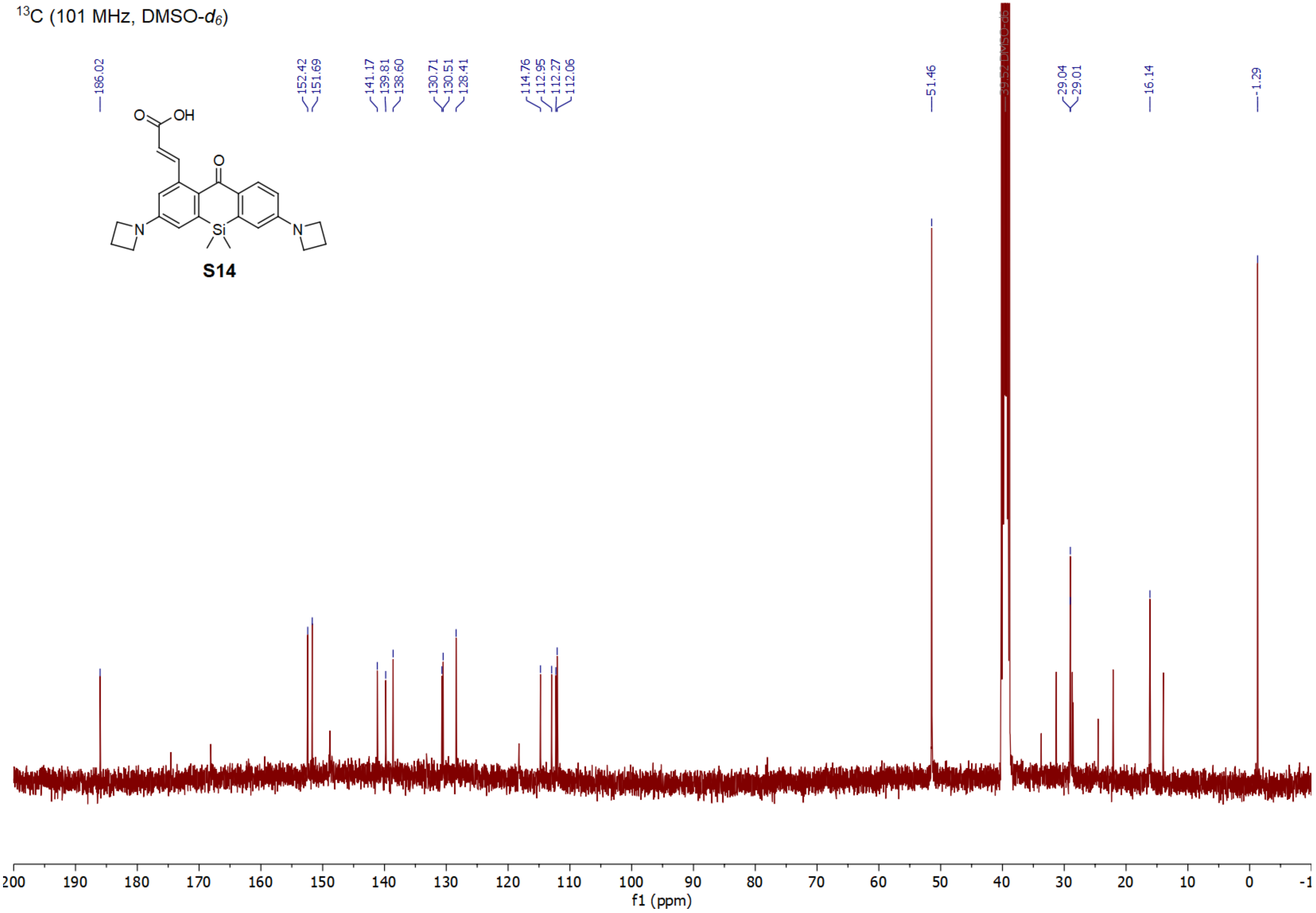
